# The winner’s curse under dependence: repairing empirical Bayes using convoluted densities

**DOI:** 10.1101/2023.09.22.558978

**Authors:** Stijn Hawinkel, Olivier Thas, Steven Maere

**Affiliations:** Department of Plant Biotechnology and Bioinformatics, Ghent University, Belgium; Center for Plant Systems Biology, VIB, Belgium; Data Science Institute, I-Biostat, Hasselt University, Belgium; National Institute for Applied Statistics Research Australia (NIASRA), University of Wollongong, Australia; Department of Applied Mathematics, Computer Science and Statistics, Ghent University, Belgium

**Keywords:** Selection bias, winner’s curse, empirical Bayes, convolution, simulation, dependence

## Abstract

The winner’s curse is a form of selection bias that arises when estimates are obtained for a large number of features, but only a subset of most extreme estimates is reported. It occurs in large scale significance testing as well as in rank-based selection, and imperils reproducibility of findings and follow-up study design. Several methods correcting for this selection bias have been proposed, but questions remain on their susceptibility to dependence between features since theoretical analyses and comparative studies are few. We prove that estimation through Tweedie’s formula is biased in presence of strong dependence, and propose a convolution of its density estimator to restore its competitive performance, which also aids other empirical Bayes methods. Furthermore, we perform a comprehensive simulation study comparing different classes of winner’s curse correction methods for point estimates as well as confidence intervals under dependence. We find a bootstrap method by Tan et al. (2015) and empirical Bayes methods with density convolution to perform best at correcting the selection bias, although this correction generally does not improve the feature ranking. Finally, we apply the methods to a comparison of single-feature versus multi-feature prediction models in predicting *Brassica napus* phenotypes from gene expression data, demonstrating that the superiority of the best single-feature model may be illusory.

## 1. INTRODUCTION

Contemporary omics technologies allow to routinely gather datasets containing many features. Commonly all these features are tested for association with a variable of interest, or some other quantity is estimated for each of these features. Scientific interest then naturally focuses on the most extreme estimates. Consequently, often only a small number of most extreme estimates or test statistics (e.g. top 10) are reported and considered for follow-up studies. Yet this subset of extreme estimates is subject to a selection bias called the *winner’s curse* (Sun et al., 2011; Zollner and Pritchard, 2007; Efron, 2011); a manifestation of the more general phenomenon of “regression towards the mean” (Galton, 1886; Bland and Altman, 1994). There are two possible reasons why the estimates are extreme: because their true estimand is extreme or because the random estimation error pushes the estimate towards the extremes. Selecting only the most extreme estimates leads to enrichment of extreme estimation errors, engendering the selection bias. More formally, if all features with estimates for a metric of interest 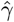 falling below a threshold *c* are selected (assuming we are interested in small values), the selection bias is defined as 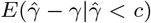. This bias is usually non-zero and negative, even when the regular estimator is unbiased 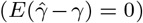. Selection bias results from using the same dataset for identifying extreme features as well as for providing the corresponding estimates, not unlike data dredging. It has been suggested as a cause for poor reproducibility of published findings and insufficient power in follow-up studies (Sun et al., 2011; Zollner and Pritchard, 2007; van Zwet et al., 2021). The winner’s curse is related to the problem of inference on model parameters after model selection (post-selection inference) (Berk et al., 2013), but here we focus purely on parameter estimation and ranking, without considering model selection.

Existing selection bias correction methods are often applied to test statistics in the realm of hypothesis testing (Ghosh et al., 2008; Zhong and Prentice, 2008; Tan et al., 2015; Ferguson et al., 2013; Faye et al., 2011; Bigdeli et al., 2016; Efron, 2011). Yet also estimation and ordering of raw effect sizes can be of interest, e.g. in cases where there is no obvious null value, like predictive modelling. Moreover, even when using test statistics, improving the ordering of top features may improve the success rate of subsequent validation efforts. Multiple testing corrections that control the false discovery rate (FDR), e.g. Benjamini-Hochberg adjustment of p-values (Benjamini and Hochberg, 1995), are widely adopted, but often not all significant features can be investigated further as resources for follow-up studies are limited. Consequently, it is common practice to only focus on a small set of top test statistics. Conversely, researchers may also consider features falling just short of the significance level when the number of discoveries is small. In such settings it is the number of findings that is fixed, rather than the rate of false findings, and the reliability of the findings might be increased by selection bias correction methods that improve the feature ranking.

Despite the many available methods for selection bias correction, and although many publications do include simulation studies (Tan et al., 2015; Faye et al., 2011; Bigdeli et al., 2016), exhaustive benchmarking studies are lacking. Also the effect of dependence among estimates on these procedures has not been thoroughly investigated. The estimates may show dependence for a host of different reasons: co-expression of genes for instance causes correlation between test statistics in differential expression testing. But also reliance on a common random quantity for the calculation of the estimates introduces dependence, e.g. a common outcome variable in prediction problems. Tweedie’s formula, an empirical Bayes method that is often used to correct for selection bias, has been found to perform poorly in the presence of dependence (Tan et al., 2015; Bigdeli et al., 2016). Tan et al. (2015) claim that dependence invalidates Tweedie’s formula, but according to Efron (2011) it remains valid and dependence merely inflates the variability of the resulting estimates. Similarly, Stephens (2017) notes the independence assumption on the estimates as a weakness of his ash method, another empirical Bayes method, but saw no obvious way to relax it. Bigdeli et al. (2016) tackle the problem of correlation for empirical Bayes methods in the context of GWAS studies by splitting the test statistics into maximally distant, and thus uncorrelated, sets of SNPs based on chromosome order, but this is not easily generalizable to other applications.

In this work we investigate the problem with empirical Bayes procedures under dependence both theoretically and empirically, and suggest a solution based on convolution. Next we benchmark the refactored empirical Bayes methods against other selection bias correction methods under dependence. Finally we apply the best performing methods to a real dataset from a *Brassica napus* field trial.

## 2. OVERVIEW OF CURRENT METHODOLOGY

Given is an *n × p* data matrix **X** with *n* samples and *p* features, possibly with a vector **y** of length *n* that is tested for association with every feature **x**_*j*_. The quantity of interest for every feature *j* is indicated by *γ*_*j*_; we assume that small values are of most interest, e.g. the features with the lowest prediction loss. We distinguish four streams of methods to tackle selection bias: 1) separate estimation, 2) conditional likelihood, 3) bootstrapping and 4) empirical Bayes.

The first method is inspired on the observation that the selection bias springs from using the same data for best feature selection as well as for parameter estimation. An obvious solution is to split the available samples at random in two sets, and use the first set for selecting the features with the smallest estimates, and the second set for providing parameter estimates of the selected features to be reported (Sun and Bull, 2005). We call this the *separate estimation* procedure.

A second method is conditional likelihood. First *γ* is estimated for all features using full maximum likelihood (ML) as 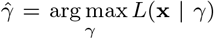 with *L*(**x** | *γ*) the likelihood of the data **x** given parameter value *γ* (omitting feature subscripts). Next conditional likelihood estimates are found for the subset of features with the smallest estimates, conditional on them falling below some threshold *c*, which may depend on the other estimates in rank-based selection: 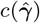 (Zollner and Pritchard, 2007; Ghosh et al., 2008). Using Bayes’ theorem, the conditional likelihood *L*_*c*_ for features passing the threshold *c* is

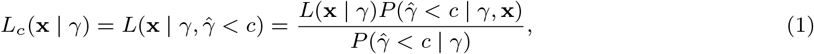

and the conditional ML estimate is 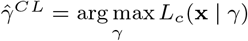. For the features concerned, 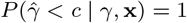. Assuming normality of the parameter estimate around the true value *γ*, 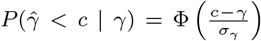with Φ the standard normal distribution function and *σ*_*γ*_ the estimate’s standard error. This probability shrinks with increasing *γ*, shifting the peak of the conditional likelihood to larger values and attenuating the selection bias of the ML estimator (Zollner and Pritchard, 2007). Conditional likelihood methods are evidently only applicable to likelihood-based estimation. Moreover, their corrected estimates depend on the choice of cut-off value *c*. A third stream of methods rely on bootstrapping to estimate the selection bias, and then correct the parameter estimates by subtracting the bias. Three distinct bootstrap methods exist: the first one, known as “BR-squared” (Faye et al., 2011), works as follows. For every *b* = 1, …, *B*:

1. Draw a bootstrap sample by sampling *n* rows with replacement from **X**, and if applicable select the corresponding elements from **y**.
2. Estimate the parameter of interest on both the bootstrap sample (in-sample), yielding 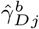 for feature *j*, and on the samples not included in the bootstrap sample (out-of-sample), yielding 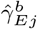.
3. Rank the in-sample estimates 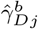 over the bootstrap sample, to yield 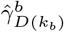, with (*k*_*b*_) indexing the feature with the k-th smallest value (k-th order statistic) in 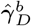. Use the same feature ordering to also order 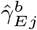 and 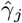 (the estimates on the real non-bootstrapped data) into 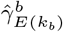 and 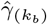, respectively.
4. Estimate the bias as 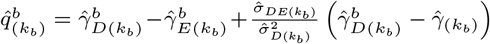. 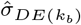 is the empirical covariance between 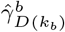and 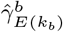, and 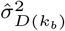 is the empirical variance of 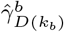, both estimated from all bootstrap samples. The last term corrects for the correlation between 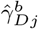 and 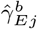 conditional on 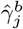 induced by their estimation from mutually exclusive sets of samples.

Finally, subtract the average bias to obtain the corrected estimate 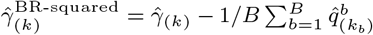 with 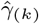 the k-th order statistic of 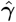. The second approach, which we will call “Tan2015” (Tan et al., 2015), works as follows:

1. Perform steps 1-3 as for BR-squared, without calculating the out-of-sample estimates.
2. Set 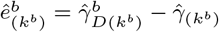 for every *b*.
3. Subtract the bias: 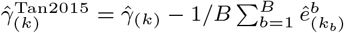

Both these bootstrap methods address dependence between estimates by performing the resampling in the first step jointly for all features, i.e. sampling entire rows of **X**. For the Tan2015 method, also parametric bootstrapping is possible if the dependence structure of the data can be estimated. A third bootstrap method is called *winnerscurse* and only generates a single, parametric bootstrap sample for every feature relying on normality of the estimates. The bias 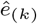 is approximated as for the Tan2015 method, and a cubic smoothing spline is fitted through the collection of 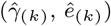 pairs. The fitted values 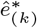 of this spline are subtracted from the raw values to correct the bias (Forde et al., 2023).

A fourth class consists of Bayesian methods, as Bayesian statistics is immune to selection bias when a common prior distribution for all features is used (Efron, 2011; Senn, 2008). For the three methods considered, some prior distribution is estimated from the data, making them *empirical* Bayes methods. Tweedie’s formula (Robbins, 1956) is the most popular one for the purpose of selection bias correction (Efron, 2011). For normally distributed estimators, its estimate of the posterior mean is:

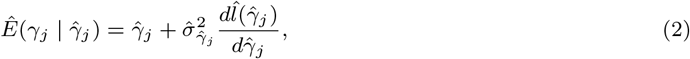

with 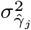 the variance of the estimator 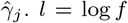 is the marginal log density, where *f* is a convolution of the prior distribution over all features *g*(*γ*) with an estimator distribution 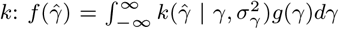 or *f* = *g* ∗ *k* for short. By exploiting the high-dimensionality of the data, the marginal log density *l* and its derivative can be estimated directly from the observed estimates 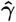, avoiding the need to specify or estimate the prior *g* explicitly (Efron, 2011). A second empirical Bayes procedure which has been applied for selection bias correction (Morrison and Simon, 2018) is adaptive shrinkage (ash) by Stephens (2017). It assumes a unimodal prior distribution *g* of which the parameters are estimated from the data, and assumes the estimates 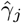 to be independently and normally distributed with variance 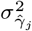. A posterior distribution 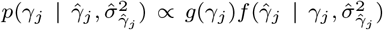 is then calculated for each feature, and the posterior mean serves as a point estimate. Note the difference between both empirical Bayes approaches: in the ash method the prior distribution of the parameters *g*(*γ*) is estimated, whereas Tweedie’s formula estimates the marginal distribution of the estimates 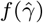, which is a convolution of the prior with the estimator sampling distribution. A third empirical Bayes method, which we will call “VanZwet2021”, estimates the distribution *f* (SNR|*z*) of the signal-to-noise-ratio (SNR) 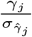 conditional on the z-value 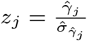 through an EM algorithm, by assuming that the prior distribution can be described by a mixture of normal distributions with mean 0. Its corrected estimate is then the posterior mean of the SNR multiplied by the standard error: 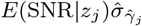 (van Zwet et al., 2021). Finally, Zuber and Strimmer (2009) construct correlation adjusted t-statistics (“cat-scores”) by decorrelating the test statistics. Although not directed against the winner’s curse, they claim that the decorrelation improves feature ranking.

Apart from point estimates, confidence intervals for the *h* smallest estimates are also desirable. The regular confidence intervals do not achieve their nominal coverage for extreme estimates, and custom interval construction is needed (Morrison and Simon, 2018). For the separate estimation method, the confidence intervals for the selected features are constructed based on the samples from the second group alone. Zollner and Pritchard (2007) and Zhong and Prentice (2008) present a 95% confidence interval for the conditional likelihood estimates consisting of all values *γ*_*j*_ for which twice the difference in log conditional likelihood with 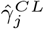 lies below the 95th percentile of a *χ*^2^-distribution with degrees of freedom equal to the number of free parameters of the model. This interval can be asymmetric. Faye et al. (2011) applied a nested bootstrap to construct a symmetric confidence interval for the BR-squared estimates using the bootstrap standard error and relying on normality of the estimator. Morrison and Simon (2018) provide the estimator by Tan et al. (2015) with a confidence interval, also employing a nested bootstrap but using the quantiles of the bootstrap distribution rather than assuming normality of the estimator. This bootstrap interval can be asymmetric to reflect the fact that small estimates are often underestimates. The Bayesian framework of Tweedie’s formula also yields an expression for a credible interval on the final estimates, relying on the normality of the posterior distribution. The posterior variance is

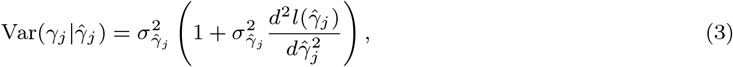

and the (1 − *α*) credible interval for *γ*_*j*_ is found as 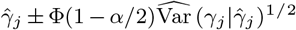 (Efron, 2011). Ferguson et al. (2013) also incorporated the variability of the estimator 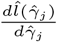 in every point 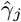 through bootstrapping. Credible intervals for the ash and VanZwet2021 methods are derived from their posterior distributions 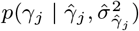 and *f* (SNR|*z*_*j*_), respectively (Stephens, 2017; van Zwet et al., 2021).

## 3. TWEEDIE’S FORMULA IS BIASED UNDER STRONG DEPENDENCE

The effect of dependence between estimates on empirical Bayes procedures is subject to debate (Tan et al., 2015; Stephens, 2017; Efron, 2011; Morrison and Simon, 2018; Bigdeli et al., 2016). Simulation studies by Tan et al. (2015) and Bigdeli et al. (2016) found that dependence negatively affects Tweedie’s formula. To elucidate why this happens, we simulated 3 Monte-Carlo instances of 1,000 independent and 1,000 dependent estimates 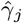 (i.e. with correlation 0 and 0.75 between all features, respectively, see Section 5.2 for details), shown in Figure 1. The distributions for independent estimates are all very similar and approximate the true marginal distribution of all estimates overlaid in blue. Dependence narrows the distribution of estimates and shifts it to higher or lower values, strongly departing from the true marginal distribution; a phenomenon known from the field of multiple testing correction (Efron, 2007; Hawinkel et al., 2022a; Azriel and Schwartzman, 2015). It is easy to envision how this behaviour upsets the performance of Tweedie’s formula (2): a naive density estimator will yield wrong derivatives of the log-density, leading to incorrect shrinkage. Analogously, the ash and VanZwet2021 methods will be affected as the width of their prior and z-value distribution, respectively, is underestimated.

**Fig. 1.**
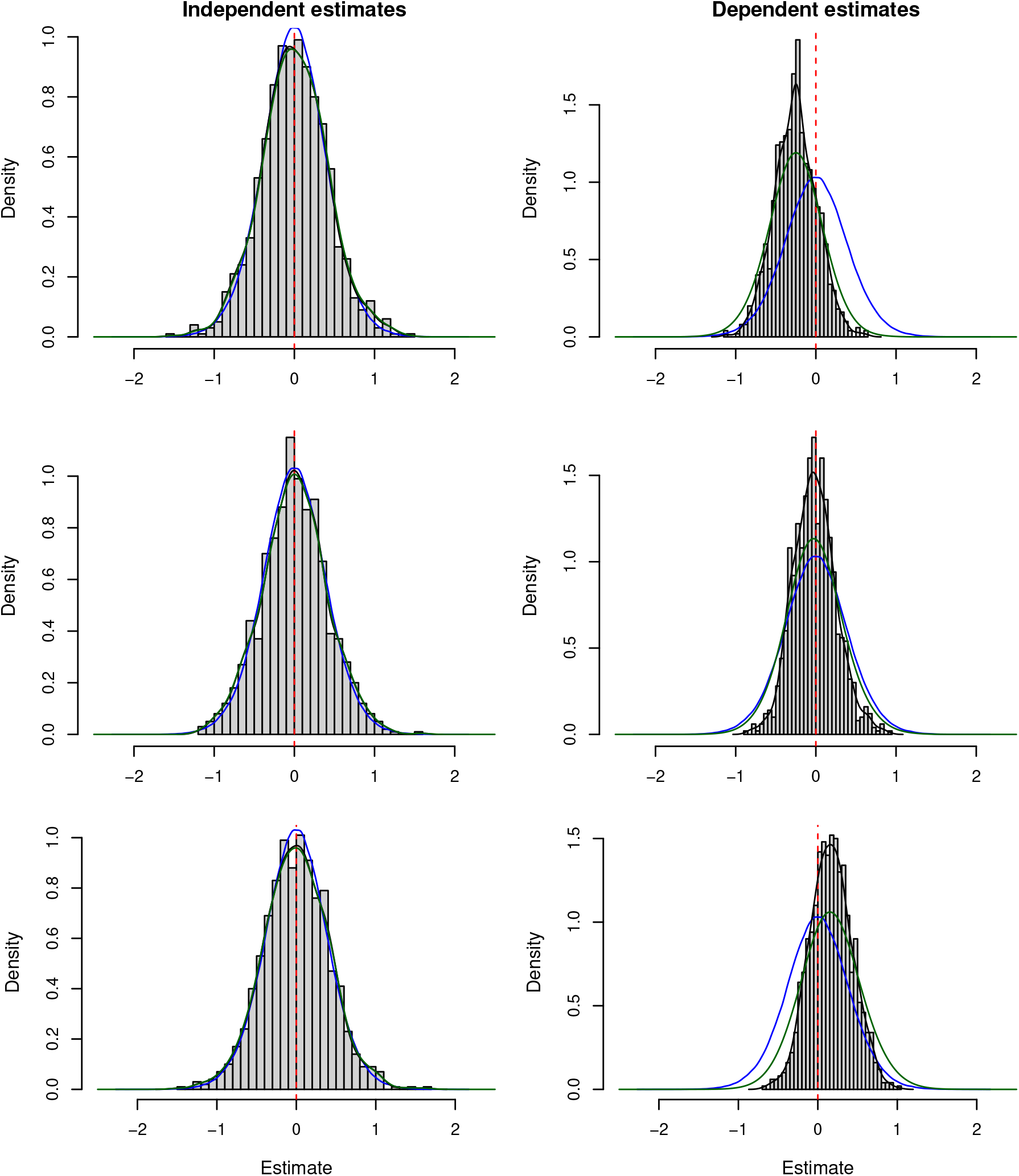
Histograms of independent estimates (left) and dependent estimates (right), see text for details. The red dotted line indicates 0. The naive density estimator is overlaid as a black line, the true marginal distribution as a blue line, and the convoluted density estimate (see Section 4) as a green line. In the left column the black, blue and green lines largely overlap. Convolution of the estimated densities under dependence yields estimates of similar spread as the true densities, even though they are still shifted.

To investigate this phenomenon more formally in line with existing theory (Schwartzman, 2010; Efron, 2011; Azriel and Schwartzman, 2015), we normalize all estimates 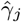 to z-values: 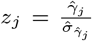 The generative model underlying Tweedie’s formula is then:

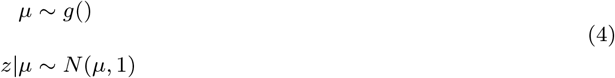

The observed distribution *f* (*z*) is a convolution of a prior distribution *g* of the expectations *µ* with an estimator distribution 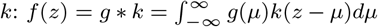. Since all z-values have the same variance 1, *k* is a mixture of *p* standard normal distributions with equal mixing proportions, which equals *ϕ*(*z*): the standard normal density. We assume for a moment that *g* is a point density, so all means *µ* are the same, and *k* and *f* coincide. Azriel and Schwartzman (2015) have shown that a naive estimator 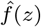 based on an observed sample **z** only does not converge to *f* (*z*) as *p* → ∞ in presence of strong dependence. Strong dependence means that the average absolute correlation between estimators does not vanish as *p* → ∞. This occurs for instance when a set of genes is tested for differential expression with respect to some phenotype in an observational study, if a confounder influences the expression of multiple genes as well as the phenotype. The resulting test statistics will then be dependent. Another example of strong dependence is the prediction loss of a common outcome variable predicted by many different models (see nonparametric simulation and case study below). Adding more features with the same dependence structure will not cause the average correlation to vanish in these cases. On the contrary, weak correlation occurs when the average absolute pairwise correlation does tend to zero as *p* tends to infinity. Weak correlation holds e.g. for autoregressive models (Azriel and Schwartzman, 2015) but also for GWAS settings with only local correlation due to linkage disequilibrium, or for gene correlation networks, as most gene pairs are not correlated. Yet strong and weak dependence are theoretical notions describing behaviour as *p* → ∞. For a given experiment, *p* is usually fixed, and could be imagined to grow in different ways, yet the analysis strategy cannot depend on such hypothetical experiments. In practice, for finite *p*, accounting for strong dependence is often indicated when the observed average correlation is away from zero. Schwartzman (2010) demonstrates how strong correlation narrows and shifts 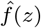, making it behave as a random function even as *p* → ∞. Following the Gram-Charlier expansion in Schwartzman (2010), more precisely his equation (8), this random function can be described as:

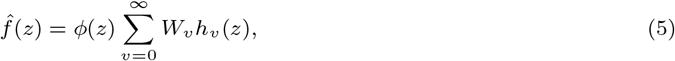

with 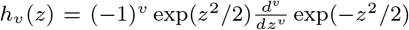 the v-th Hermite polynomial (*h*_0_ (*z*) = 1, *h*_1_ (*z*) = *z, h*_2_ (*z*) = *z*^2^ − 1, *h*_3_(*z*) = *z*^3^ − 3*z*, …). *W*_0_ = 1 and the other *W* ‘s are random variables with mean 0 and variance-covariance structure

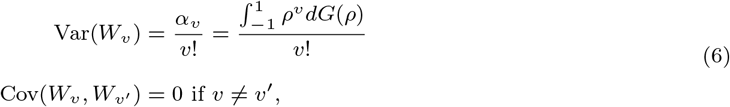

with *G*(*ρ*) the distribution function of pairwise correlations *ρ* between z-values, and *α*_*v*_ the v-th moment of the correlation distribution. Repeated experiments correspond to repeated draws **W** = (*W*_0_, *W*_1_, …, *W*_∞_)^*t*^. The term *W*_1_*h*_1_(*z*) is responsible for the shift in location of 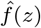, whereas the terms *W*_1_*h*_1_(*z*) and *W*_2_*h*_2_(*z*) drive its width, usually narrowing 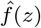 with respect to *f* (*z*) (Schwartzman, 2010). These departures from f(z) upset empirical Bayes methods, even though the correlation does not affect the expectation over different experiments: 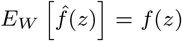. Hence 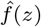 is an unbiased estimator for f(z) but not consistent in the sense that is does not converge to *f* (*z*) as *p* → ∞ (Schwartzman, 2010).

Tweedie’s formula relies on the first derivative of the logarithm of the marginal density function, so we investigate the behaviour of this random function too. Taking the logarithm and derivative of (5) yields

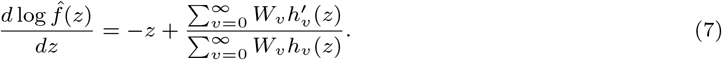

We now take the expectation over random realizations of **W**, so not over z as in equation (11) in Schwartzman (2010). For ease of notation we set 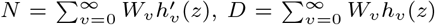, both of which are functions of **W** and z, with *E*_*W*_ (*N*) = 0 and *E*_*W*_ (*D*) = 1. The term *N/D* is in general not zero and causes bias in the estimation of the shrinkage term in (2). Its second order Taylor expansion (Seltman, 2019) provides the following approximation:

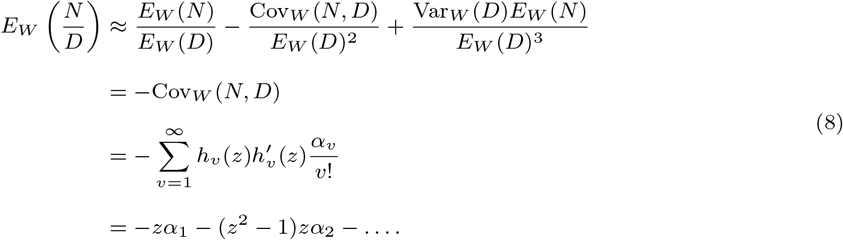

Hence even though 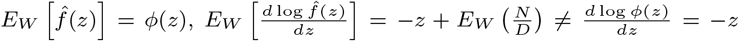 because of the dependence. A special case is z=0: since the Hermite polynomials contain only even or only odd powers of z,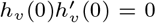 for all *v* and thus (8) equals 0. In other words, the naive estimator for the derivative of the log-density is biased for values other than z=0, with bias growing in absolute value as z moves away from 0. Next we approximate the variance of expression (7), which we rewrite as *b*(**W**) = −*z* + *N/D*, through the first-order delta method (Gauss, 1823) as

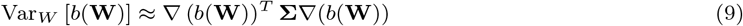

The variance-covariance matrix **Σ** is diagonal, see (6). The *v*-th element of ∇ (*b*(**W**)) is given by

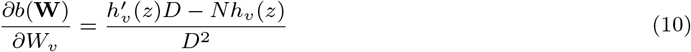

Evaluated in the expected value *E*(**W**) = (1, **0**)^*t*^, with **0** a vector of zeroes of infinite length, this equals

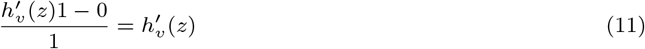

Plugging this into (9) yields (since Var(*W*_0_) = 0)

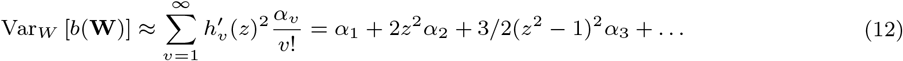

Hence the variance grows as *z* moves away from zero. Ferguson et al. (2013) already observed that the variability of the estimator 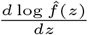 is larger in the tails because there are fewer observations, causing inaccurate shrinkage. This is sampling variability that is expected to disappear as *p* → ∞. Yet our results (8) and (12) uncover two more problems under correlation: 1) the regular estimator for 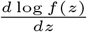 is biased for *z* ≠ 0, and 2) 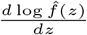 is itself a random function with a variance of at least *α*_1_ (the average pairwise correlation), even as *p* → ∞. Both absolute value of the bias and variance grow as *z* moves away from 0. Problem 1) adds nuance to the assertions by Efron (2011) that Tweedie’s formula remains valid under dependence: it is biased when *z*Σ 0 under strong dependence in combination with a naive estimator for *f* (*z*) that assumes independent z-values.

In general, *g* is not a point density, such that *f* is a mixture of normal distributions with differing means and only 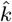 follows (5) with expectation *ϕ*(*z*) as 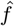 takes on other shapes. The derivative of the marginal log-density then becomes

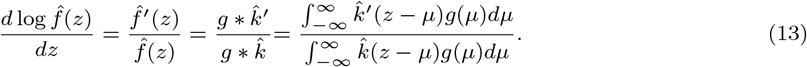

This implies that the shape of the bias function depends on the prior *g*. When the spread of *g* is the main driver of the spread of *f*, which happens e.g. when *n* → ∞, the change in shape due to the correlation is negligible and the derivative of the log-density is approximately unbiased. Yet in that scenario, the estimator variances 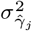 are small compared to the variability of the true values *γ*_*j*_, and the effect of winner’s curse would be minor. Correcting for winner’s curse is most relevant when 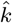 is the dominant source of variance of the estimates 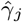, of which *g* being a point density is the most extreme case.

In the field of multiple testing correction, precise control or estimation of the false discovery rate while classifying the ensemble of all features into significant and non-significant is often the goal. This means that the significance of a certain feature depends on all other features’ test statistics, and the distribution of a given experiment 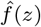 is of primary concern (Efron, 2007; Hawinkel et al., 2022b; Azriel and Schwartzman, 2015). Yet when estimation of the parameters *γ* is the main purpose, the estimates of other features are not of direct interest as estimation occurs feature-by-feature. The other features’ estimates simply provide a way to estimate the prior distribution *g* or the true marginal distribution *f* (*z*), which are needed for Bayesian shrinkage. Yet we have proven that a naive estimator 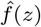 is not a good choice under strong dependence, so we need a better estimator for *f* (*z*).

## 4. CONVOLUTION FOR IMPROVED DENSITY ESTIMATION UNDER DEPENDENCE

Schwartzman (2010) has shown that 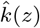 has expected variance 1 − *α*_1_ over repeated experiments **W**:

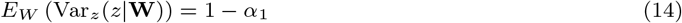

with *α*_1_ the average pairwise correlation between the *z*_*j*_’s. Azriel and Schwartzman (2015) add that 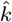 can be described by a mixture of *M* normal distributions:

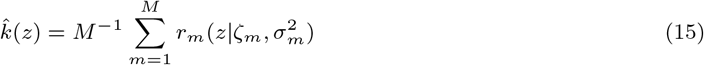

with *r*(.|*ζ, σ*^2^) the normal density with random mean *ζ* (with expectation 0) and variance *σ*^2^. In our case study (see Section 6), a common outcome variable is causing a compound symmetric correlation structure (see Supplementary Figure S21), such that 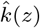 can be described using a single normal distribution (M=1) with random mean 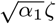 (with *ζ* standard normal) and variance 1 − *α*_1_ (Azriel and Schwartzman, 2015). The derivative of its log-density equals 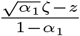 with expected value 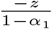. For small *α*_1_ this is similar to (7) with the approximation in (8) truncated after one term, which equals −*z*(1 + *α*_1_), proving again that Tweedie’s formula is biased when combined with a naive density estimator.

To counter this bias, knowing that correlation narrows 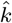 with respect to *k*, we propose to widen 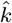 by convoluting it with a zero-mean normal distribution with variance *α*_1_. We choose a normal distribution because of its straightforward, analytical convolution with other normal distributions. In the theoretical model (15), this would mean to convolute every component with *r*(*z*|0, *α*_1_), yielding

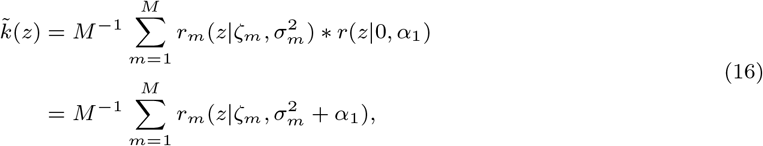

with expected variance (1 − *α*_1_) + *α*_1_ = 1. Consequently, we could use 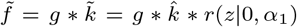 as an estimator of *f*. Yet choosing *M* and estimating the parameters *ζ*_*m*_ and 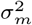 of 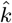 is difficult as all we observe are the estimates **z**, reflecting variability of both *g* and 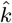. Sidestepping the deconvolution of *g* and *k* is a key advantage of Tweedie’s formula, so in practice we do not calculate (16). In this spirit, since our estimates 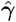 are drawn from 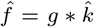, we could estimate 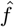 and then convolute it with *r*(*z*|0, *α*_1_). Yet even simpler, and avoiding density estimation, is to convolute the observed distribution, formed by point masses of 1*/p* at each *z*_*j*_, with *r*(*z*|0, *α*_1_) and estimate 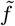 using the following mixture:

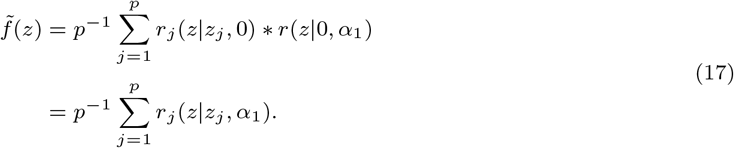

This expression (17) is conceptually equivalent to (16), but much easier to calculate. If M=1, unbiasedness of the estimator for the derivative of the log-density in (13) based on (17) can be proven for the case where *g* is a point density or a Gaussian distribution (see Supplementary Section 1.2), but not for arbitrary shape of *g*. For more complex strong correlation structures (M*>*1) and/or for arbitrary *g*, 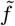 will still have the correct expected variance but unbiasedness of the estimator for the derivative of the log-density can no longer be guaranteed for all values of *z*. Still, numerical studies suggest a reduced estimation bias for the example of block correlation too (Supplementary section 2.1.2.2). Hence our estimator 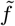 should be seen as a working solution that captures the bulk of the dependence structure, similar to the empirical normal null by Efron (2010).

On the scale of the original estimates 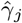, an analogous reasoning leads to the following estimator:

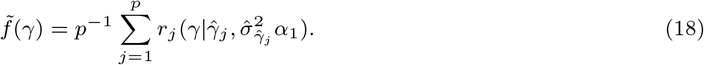

*α*_1_ is estimated as the average of all pairwise correlations between 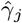’s (or a subsample thereof), which are themselves estimated either through likelihood theory or by the bootstrap. Simulations show that the convolution estimate 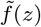 under dependence is on average as wide as *f* (*z*) but remains shifted with respect to it (see Figure 1). There is no way of correcting this, as from a single realization **z** we have got no way of knowing in which direction it is shifted. In case 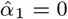, no convolution is necessary and the regular Tweedie’s formula can be employed. If 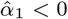, it cannot be smaller than 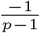 due to non-negative definiteness constraints on the covariance matrix (Schwartzman, 2010), and convolution is also not needed. In practice, since *p* is finite, small positive values of 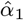 may lead to irregular estimates 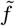, so we do not apply convolution when 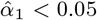. Instead, to stabilize the density estimation in the tails, we obtain bootstrap estimates of the *γ*’s by resampling samples with replacement from the data, and estimating *γ* from them. The density is then estimated on the collection of all bootstrapped estimates, a paradigm known as bootstrap aggregating or bagging (Bourel and Cugliari, 2019; Breiman, 1996). The workflow of the modified Tweedie’s formula with convolution or bagging is then as follows:

1. Obtain estimate 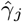 and its standard error 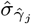 for every feature *j*.
2. Estimate *α*_1_ as the average pairwise correlation between estimates 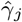_*j*_.
3. If 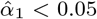, estimate the marginal log density with density bagging. We use Lindsey’s method for density estimation, as this allows a direct estimate of the log density and its derivative (Lindsey, 1974) and is less biased than nonparametric density estimators (Efron and Tibshirani, 1996). If 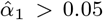, estimate the first and second order derivative of the log density based on (18) (see Supplementary section 1.1 for the derivatives) by plugging in 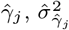 and 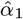.
4. Plug the estimates from steps 1 and 3 into (2) and (3) to obtain empirical Bayes estimates and the associated credible intervals.

Like Tweedie’s formula, the ash method assumes the estimates 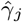 to be drawn from a convolution of the prior *g* with a normal sampling distribution *k*. Yet in reality *g* is convoluted with 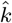, leading ash to underestimate the width of *g* and hence also of the posterior. As a solution we estimate ash’s prior using random samples from the distributions 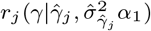; a sample size of 10 per feature suffices to obtain an estimate 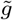 with spread similar to that of the true prior (see Supplementary Figures S9 and S11). The posterior distributions are then found as 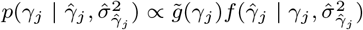. Analogously, we modify the VanZwet2021 method by estimating the distribution of z-values from the same random samples from 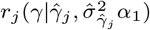 divided by 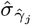, which leads to a more accurate estimation of the width of the distribution of the z-values under dependence (see Supplementary Figures S10-S11). No instability due to small *α*_1_ was observed for the ash or VanZwet2021 methods, so the convolution was applied for all cases in which 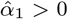.

## 5. EMPIRICAL VALIDATION

In the following sections, we first validate the theoretical findings from Sections 3 and 4 empirically (Section 5.1). Next, we investigate the performance of the 4 methods for selection bias correction (separate estimation, conditional likelihood, bootstrapping and empirical Bayes) in a parametric simulation study in which feature means are estimated (Section 5.2) and a nonparametric simulation study in which the predictive performance for a common outcome variable is estimated for many features (Section 5.3). Finally all methods are applied to real dataset from a *B. napus* field trial (Section 6). All analyses are run in R 4.4.0, see Supplementary Section 7 for the package versions.

### 5.1. Empirical validation of theoretical properties

In parametric simulations, multivariate Gaussian data **X** of dimension *n × p* are drawn. A mean vector ***µ*** of length *p* is generated either a) through p independent draws from a normal distribution with mean 0 an variance *η*^2^ (the dense scenario) or b) the first 0.9p entries are set to 0 and only the last 0.1p elements are drawn from a normal distribution with mean -1 (as we focus on the lowest estimates) and variance *η*^2^ (the sparse scenario). The variance vector for the columns of **X, *σ***^2^, is generated by *p* independent draws from a uniform distribution on [*υ* _1_, *υ* _2_]. The correlation *ρ* is identical for all feature pairs features and set to either 0, 0.25, 0.5 or 0.75. The estimand *γ*_*j*_ is the mean *µ*_*j*_ for every feature. Its ML estimate, under the assumption of homoscedastic normality, is 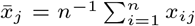, referred to as the raw estimate. Its standard error 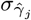 is estimated as 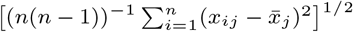.

We validate the theoretical results on the derivative of log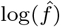 from Section 3 numerically. We use the dense parametric scenario, setting *n* = 1, *η*^2^ = 0 and *υ* _1_ = *υ* _2_ = 1, so *g* is a point density and the estimates 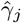 are standard normal, making the true *f* (*z*) also standard normal. Of course, the simulation results will also reflect sampling variability, so we choose *p* ∈ (500; 1, 000; 5, 000) to have it decrease. For each combination of *ρ* and *p* we perform 5,000 Monte-Carlo simulations to approximate the bias, variance and mean squared error (MSE) with respect to 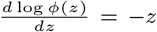 across a range of z-values, using Lindsey’s method as density estimator (Lindsey, 1974). The approximate bias is shown in Figure 2 and confirms the results from (8): the bias is close to 0 for *ρ* = 0, but grows in absolute value with increasing *z* when *ρ* ≠ 0. For low correlations, the growth is linear as it is dominated by the first term in (8) (shown as dashed line), but for larger correlations also the higher order terms kick in. The variance and MSE are depicted in Supplementary Figure S1. The variance is inevitably larger than predicted by (12), as it also includes sampling variability and the first-order approximation of the variance through the delta method may be inaccurate, but as expected a larger *p* yields more precise estimates. The second order term provides a reasonable approximation for the shape of the variance function.

**Fig. 2.**
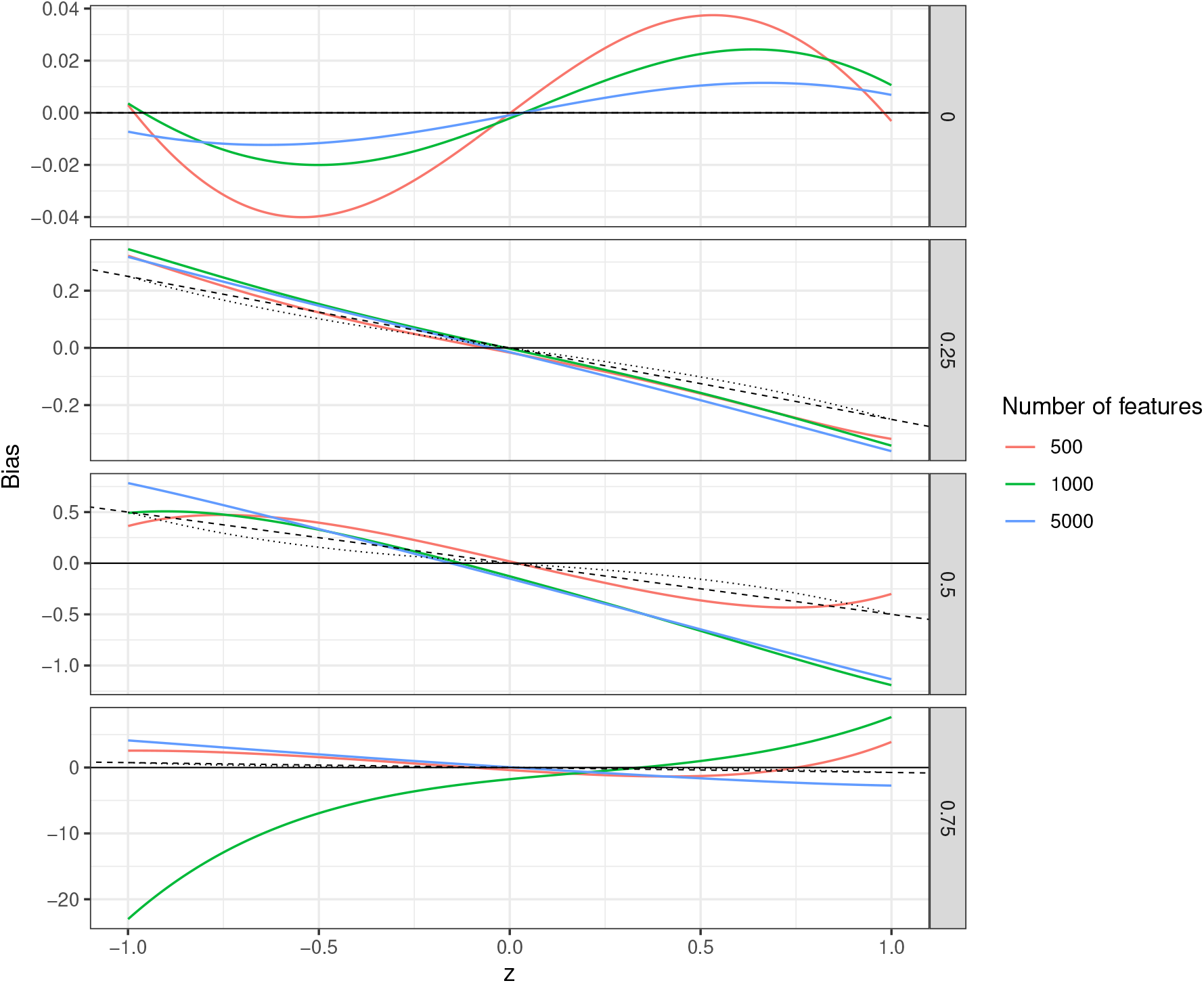
Bias in the estimation of the derivative of the log density of *f* (y-axis) as a function of z-value (x-axis), correlation between features *ρ* (rows) and number of features *p* (colour). The horizontal solid line indicates 0 (no bias), the dashed and dotted lines indicate first order and second order approximations to the bias as in (8) respectively. Notice the different scales of the y-axis; the bias grows in absolute value with *ρ*.

Next, we test the convolution and bagging estimators for the derivative of the log-density from Section 4 in a second simulation study, with the same settings as before except that now *n* = 10, *υ* _1_ = 2 and *υ* _2_ = 10. Setting *n* = 10 allows for estimating *α*_1_ as the average Pearson correlation between the different columns of **X**. Figure 3 reveals that the convoluted density yields an unbiased estimator of the derivative of the log marginal density for large *p*, although it still suffers from small sample bias in the tails when *ρ* = 0. The bagging estimator largely remedies this bias, although it becomes biased itself in the tails as *ρ* grows (see Supplementary Figure S2). Supplementary Figures S3 and S4 show that the convolution and bagging estimators also have lower variability than the naive estimator.

**Fig. 3.**
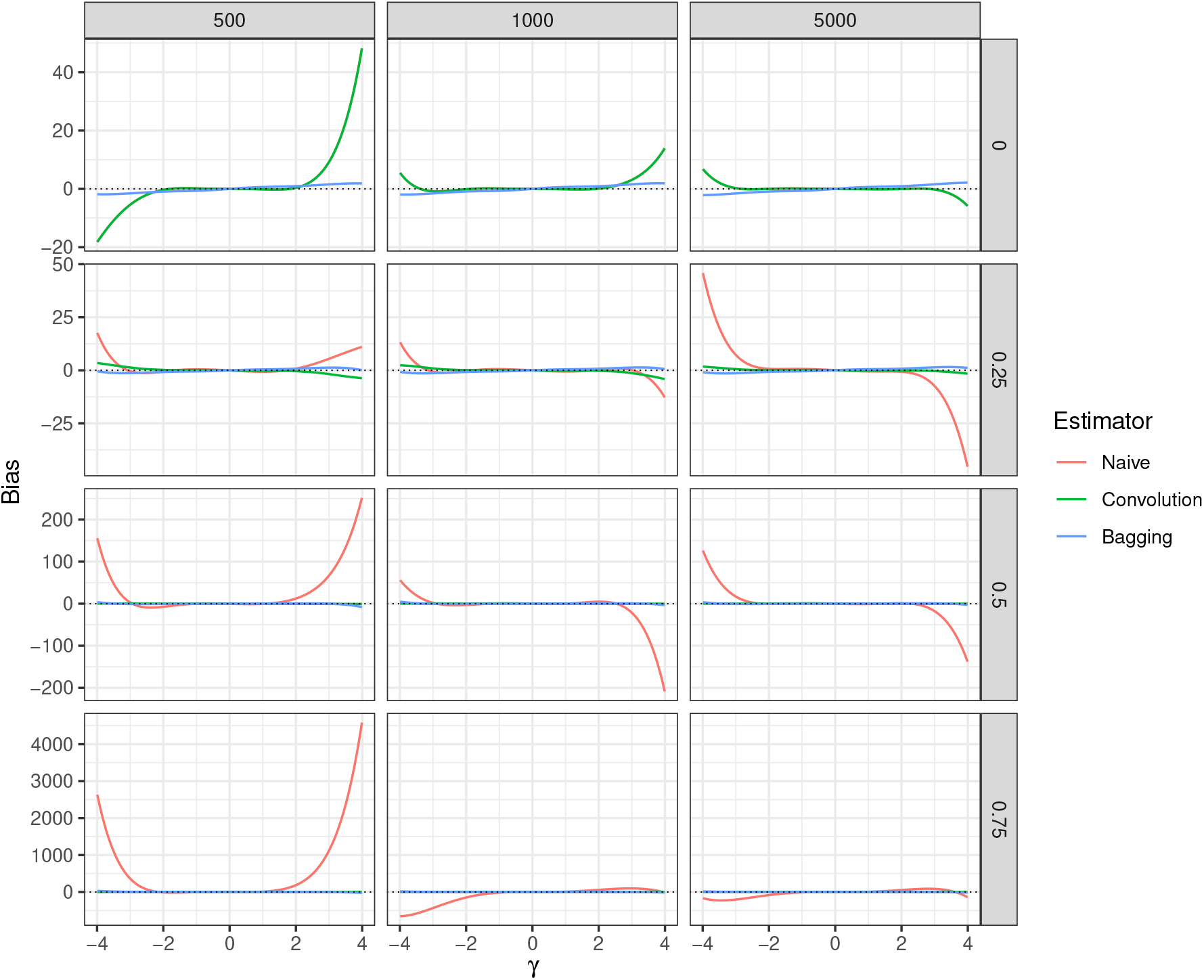
Bias in the estimation of the derivative of the log density of *f* (y-axis) as a function of estimand *γ* (x-axis), correlation between features (rows), number of features *p* (columns) and estimator (colour). The horizontal dotted line indicates 0 (no bias). For a correlation of 0, most often no convolution is applied such that the red and green lines overlap.

### 5.2 Empirical comparison of correction methods: parametric simulations

In a parametric simulation on mean estimation evaluating the correction for winner’s curse, we set *n* = 50, *p* = 1, 000, *η*^2^ = 0.04, *v*_1_ = 2 and *v*_2_ = 10, and use both the dense and sparse scenarios outlined in Section 5.1. 500 Monte-Carlo instances are generated per correlation strength and scenario. A naive confidence interval for the raw estimates is constructed as 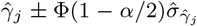 with *α* = 0.05. Separate estimation proceeds by randomly assigning the samples to two equally sized groups, and estimating the *γ*_*j*_’s and 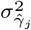 ‘s in every group separately. For the conditional likelihood estimation, the cut-off value *c* is chosen at the *h*-th smallest raw estimate with *h* = 1, …, 30. For all features falling below the cut-off, the corresponding conditional likelihood (1) is optimised over *γ*_*j*_ to yield the final estimate. A *χ*^2^-distribution with 2 degrees of freedom is used to construct the confidence interval. For the methods relying on bootstrapping, nonparametric bootstrapping proceeds by sampling rows with replacement from **X**, retaining correlation between parameter estimates. For parametric bootstrapping for the Tan2015 method, a multivariate normal distribution was fit to the data using maximum likelihood, from which *n* samples were drawn to form the bootstrapped matrix **X**^*b*^. In both cases *B*=200 bootstraps are performed. Tweedie’s formula with and without convolution was implemented.

Following Ferguson et al. (2013) we also incorporate the variability of the estimator 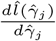 in every point 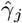 through bootstrapping. This is done by sampling *p* feature indices with replacement, and estimating the density from all bootstrap estimates 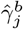 of the corresponding features. This process is repeated 100 times; the variance of the derivative of the density estimate at 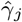 over all repeats is used as an estimate of 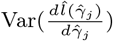. Since the error on 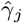 and on 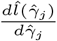 are approximately independent, this leads to a credible interval

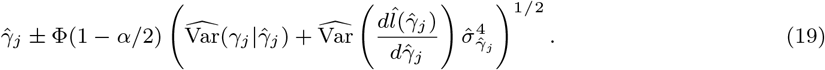

The ash method was applied using the *ash* function in the *ashr* R-package (Stephens et al., 2022), both on raw estimates only and using convolution. The *ashci* function was used to construct credible intervals. Since we focus on estimation, VanZwet2021’s assumption of the z-values (and SNRs) to have zero mean was relaxed, and the z-values are instead modeled as a mixture of normals with estimated means. This mixture is estimated through an EM algorithm similar to the original method, provided by Erik van Zwet in personal communication. The resulting algorithm is very similar to the one proposed by Ferguson and Chang (2020) to improve feature ranking, with only slight differences in the prior specification. In addition, the raw estimates were mean centered prior to estimation, and the means were re-added to the corrected estimates and their credible intervals after the VanZwet2021 correction.

For the purpose of feature ranking, also test statistics for testing the null hypothesis *H*_0_ : *γ*_*j*_ = 0 are calculated as 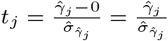. The rationale is that the inclusion of estimator variance may improve the feature ranking. The same correction methods as for the raw estimates were applied and the smallest *h* corrected test statistics were considered as discoveries. In addition, the correlation adjusted t-statistics (cat scores) implemented in the *shrinkcat*.*stat* function in the *st* R-package (Opgen-Rhein et al., 2021) were calculated. The *ash* method applied to test statistics with equal variances leaves the feature ordering unchanged, regardless of how the prior is estimated, and was therefore omitted from feature ranking comparisons.

As performance measures, we calculate the following quantities by averaging over all Monte-Carlo instances: average estimation error (as approximation of the selection bias), and the average MSE of the 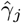’s for the *h* = 1, …, 30 features with smallest estimates. In addition, the true discovery rate (TDR) is approximated as the average proportion of the *h* features with smallest estimates 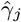 or test statistics *t*_*j*_ that are part of the *h* features with smallest true *γ*_*j*_ values. The feature ranking is thus always evaluated with respect to the true raw values *γ*_j_, rather than the true test statistics 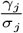. Further the coverage and width of the confidence and credible intervals for the *h* smallest point estimates were calculated. Two different notions of coverage exist for a series of selected and ranked estimates. One is the false coverage rate (FCR) by Benjamini and Yekutieli (2005), defined as the expected proportion of intervals not covering their true value among an entire set of *h* selected features. We refer to 1-FCR simply as the coverage, and this is shown unless mentioned otherwise. Yet a method controlling the FCR may still show undercoverage for the intervals around the most extreme estimates, and overcoverage for the less extreme estimates. As an alternative, Morrison and Simon (2018) present the concept of rank conditional coverage (RCC), which is the expected proportion of confidence intervals covering the true value for the feature with rank *k* point estimate. The empirical Bayes credible intervals are also evaluated according to these frequentist definitions of coverage. Finally, the average MSE of all estimates is calculated per Monte-Carlo instance, and its distribution over the Monte-Carlo instances is shown as a boxplot. Except for the RCC and the MSE of all estimates, all performance measures are thus calculated cumulatively over the *h* smallest estimates. For the separate estimation method, TDR is shown for the feature selection part of the data, the bias, MSE and confidence intervals are calculated for the final estimates based on the estimation part.

The effect of convolution on the empirical Bayes methods is illustrated in Figure 4 for the dense scenario and in Supplementary Figure S8 for the sparse scenario. In both scenarios, all empirical Bayes methods degrade in performance as the correlation between features increases, especially Tweedie’s formula. ash is particularly affected by the correlation in the dense scenario in terms of bias and coverage, whereas VanZwet2021 performs comparatively worse in the sparse scenario, presumably reflecting stronger departures from their assumptions on the prior distribution. Yet all empirical Bayes methods attain similar or better performance over all performance metrics after the inclusion of convolution (or bagging for Tweedie’s formula). Still, at high correlation, also with convolution these methods tend to overcorrect the selection bias and their credible intervals do not reach a coverage of 95%. Next we compare the other methods for correcting winner’s curse with the empirical Bayes methods, for the latter we only consider the versions with convolution (or bagging) from now on. The results in Figure 5 and Supplementary Figures S12-S18 show how the selection bias of the raw estimates decreases with the number of top features and with increasing correlation between features. This latter effect has been proven by Tan et al. (2015). Also, the winner’s curse is weaker in the sparse than in the dense scenario. The coverage of the naive confidence intervals falls short of the nominal level of 95%. The separate estimation method reduces the MSE of the top estimates with respect to the raw estimates in most cases in the dense scenario, but not in the sparse scenario, and its confidence intervals achieve the nominal coverage in both scenarios. In the dense scenario, it is immune to selection bias and its performance is not affected by dependence, but in the sparse scenario it still suffers from some selection bias at high correlations. Moreover it has wide confidence intervals and low TDR. Conditional likelihood estimation greatly reduces selection bias of top estimates, but overcorrects when the correlation between the estimates is high. Its estimates are increasingly imprecise under increasing correlation and its confidence intervals are wider than those of the separate estimation method, and slightly overcover. The BR-squared correction increases selection bias and MSE in case of independence between features; its feature ranking performance is also very bad in this case and its confidence intervals are narrow but undercover. As the correlation between features grows higher, its performance improves over all these performance measures, becoming good under high correlation. The Tan2015 bootstrap correction works well irrespective of correlation strength, reducing selection bias and MSE of the smallest estimates, and the coverage of its confidence intervals hovers around the nominal level under independence but drops under dependence. Its feature ranking is often slightly worse than that of the raw estimate. The *winnerscurse* method performs reasonably well at correcting selection bias in the dense but not in the sparse scenario.

**Fig. 4.**
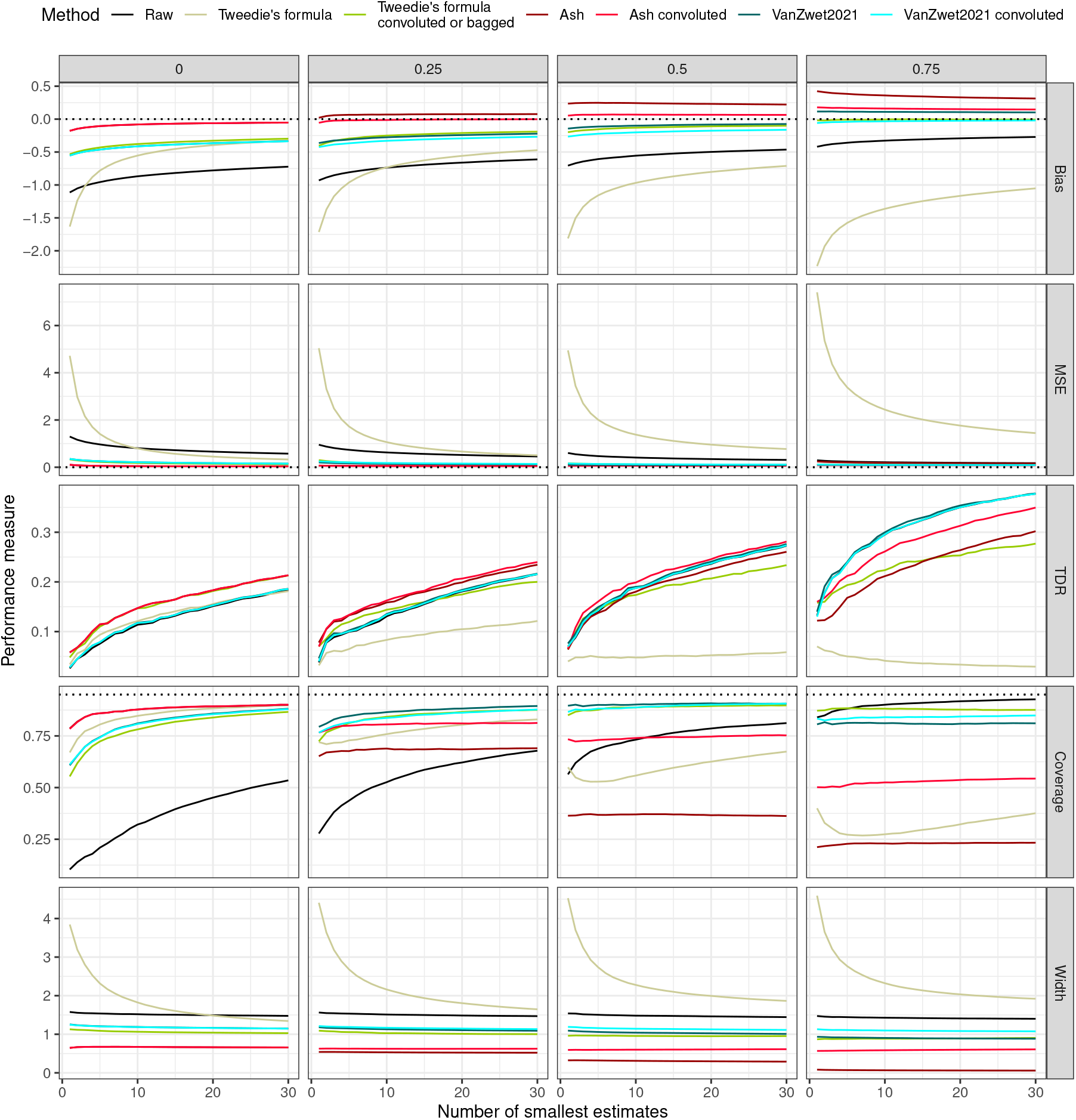
Parametric simulation results in the dense scenario for different performance measures (side panels) for the raw estimates and naive and convolution empirical Bayes methods (colour) for increasing correlation between features (top panels). The x-axis shows the number of features with smallest estimates considered. The dotted horizontal line indicates zero bias or MSE or the nominal coverage of 95%, respectively. For zero correlation, the convoluted versions mostly overlap with the naive versions.

**Fig. 5.**
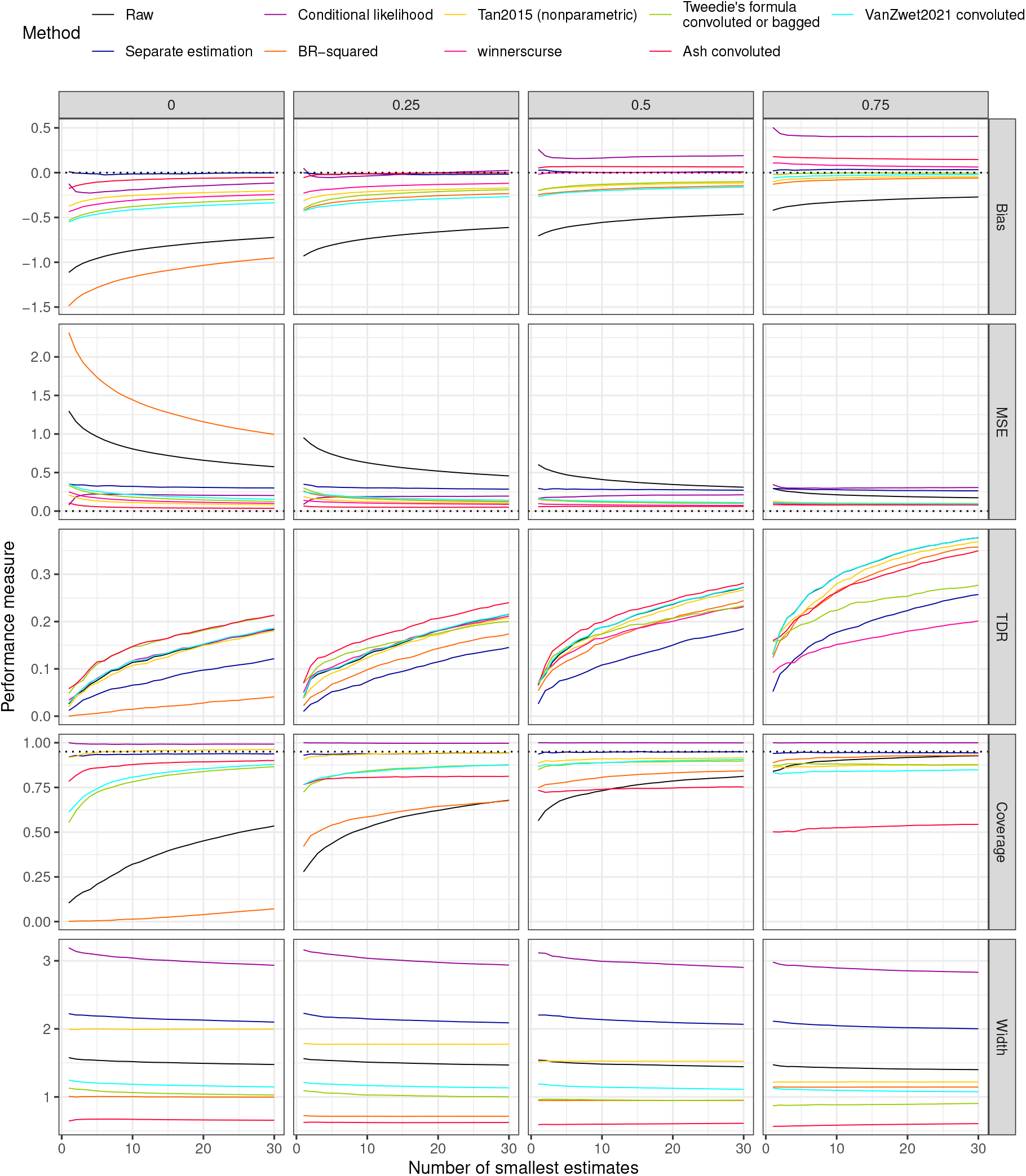
Parametric simulation results in the dense scenario for different performance measures (side panels) as a function of number of features with smallest estimates considered (x-axis) for different methods (colour) and increasing correlation between features (top panels). The feature ranking is the same for conditional likelihood as for the raw estimates, so in this case only the latter are shown. The feature ranking of VanZwet2021 is very similar to the one of the raw estimates, so their lines largely overlap. The dotted horizontal line indicates zero bias or MSE or the nominal coverage of 95%.

Feature ranking using test statistics works better than using only raw estimates for most methods in the dense scenario, but not in the sparse scenario (see Supplementary Figure S16). The raw test statistics are mostly among the best performers in terms of TDR, only under the highest correlation in the dense scenario do the cat scores provide a slight improvement (see Supplementary Figure S14). Bringing the corrected test statistics back to the scale of the raw estimates by multiplying by the standard errors as suggested by several authors (Ferguson et al., 2013; Ghosh et al., 2008) did not offer a consistent improvement across performance measures and scenarios for any method (Supplementary Figures S14-S17). When features are selected based on significance after FDR-control rather than selecting the smallest *h* features (see Supplementary Section 2.2.5), all methods perform much better in the sparse scenario than in the dense scenario. Yet the methods for correcting winner’s curse offer most improvement in the dense scenario. This is likely because the 0.1*p* features with mean away from 0 in the sparse scenario can easily be selected as significant, whereas ranking the features in the dense scenario is much harder.

### 5.3. Empirical comparison of correction methods: nonparametric simulations

In a nonparametric simulation study, the predictive performance for a common outcome variable (a plant phenotype) is being estimated for many features (gene expression values), from data nonparametrically simulated based on a real dataset. For this simulation we use a dataset on 62 field-grown *Brassica napus* plants of the same genetic background, for which the leaf transcriptome was measured in November 2016, and a number of phenotypes were measured at harvest in June 2017 (De Meyer et al., 2023). Six phenotypes where chosen to be predicted from the transcriptome: leaf 8 width and length, leaf count, total seed count, plant height and total shoot dry weight. The gene expression measurements were *rlog* transformed to stabilize their variance (Love et al., 2014), and only the 1,000 most expressed genes were retained as predictors. The quantity of interest *γ*_*j*_ is the out-of-sample root mean squared error (RMSE), which quantifies predictive performance (Hastie et al., 2009). The RMSE and its standard error are estimated through 10-fold cross-validation (CV) (Bates et al., 2023) (see Supplementary Section 4 for details). The prediction model applied is a linear model estimated by generalised least squares (GLS) with Gaussian autocovariance structure as implemented in the *gls* function in the *nlme* R-package (Pinheiro et al., 2021) to account for spatial autocovariance across the field. In the simulation, the samples of the *B. napus* data are split at random into two equally sized groups, and the *γ*_*j*_ of every feature is estimated separately in both groups. Next the selection bias correction methods are applied to the estimates from the first group, and their accuracy is assessed by considering the corresponding estimates from the second group as ground truth (Tan et al., 2015). We repeat this splitting procedure 20 times, which allows us to assess the performance metrics bias, MSE, TDR and width of the confidence interval of top estimates on real data. Estimates from the first and second group exhibit correlation as they are estimated from mutually exclusive sets of samples, and also estimates from the second group exhibit sampling variability. Hence the estimates of bias, MSE and TDP in this nonparametric simulation study rely on an imperfect ground truth and should be considered as approximations, with the purpose of comparing the performance of methods correcting for winner’s curse. The same modification to the VanZwet2021 method was made as in the parametric simulation. The estimators for *γ* and its standard error covary, which leads to overshrinkage of some of the large raw estimates 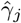 by Tweedie’s formula, even into ranges smaller than the smallest raw estimates. Since we are only interested in small *γ* values, as a solution only the 50% smallest raw 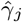’s are shrunk according to Tweedie’s formula, the rest are left unchanged. We will call this the truncated Tweedie estimates, and results are shown only for them. For the purpose of feature ranking, the estimates were decorrelated in the style of the cat scores by premultiplying the estimates by the inverse square root of their estimated correlation matrix. This correlation matrix was estimated on the matrix of bootstrapped estimates using the *crossprod*.*powcor*.*shrink* function in the *st* R-package (Opgen-Rhein et al., 2021).

Figure 6 and Supplementary Figure S20 reveal that the separate estimation method has a (slight) positive bias and low TDR; its MSE of top estimates and of all estimates is higher than for the raw estimates. The Tan2015, Tweedie’s formula and VanZwet2021 corrections have the lowest bias and MSE for the top estimates, and ash and VanZwet2021 achieve the lowest overall MSE. Also the truncated Tweedie estimates improve upon the raw estimates for all phenotypes in terms of bias and MSE of top estimates. The BR-squared method reduces bias and MSE for most outcome variables, but not as much as the other methods. The *winnerscurse* method performs poorly overall. Convolution improves the performance of the empirical Bayes methods, although ash’s credible intervals remain unlikely narrow even with convolution. The credible intervals of the convoluted Tweedie’s formula and VanZwet2021 are narrower than the confidence intervals for the raw estimate, but appear more believable as they are comparable in width to those of the raw values. Decorrelating the estimates degrades feature ranking accuracy for most phenotypes.

**Fig. 6.**
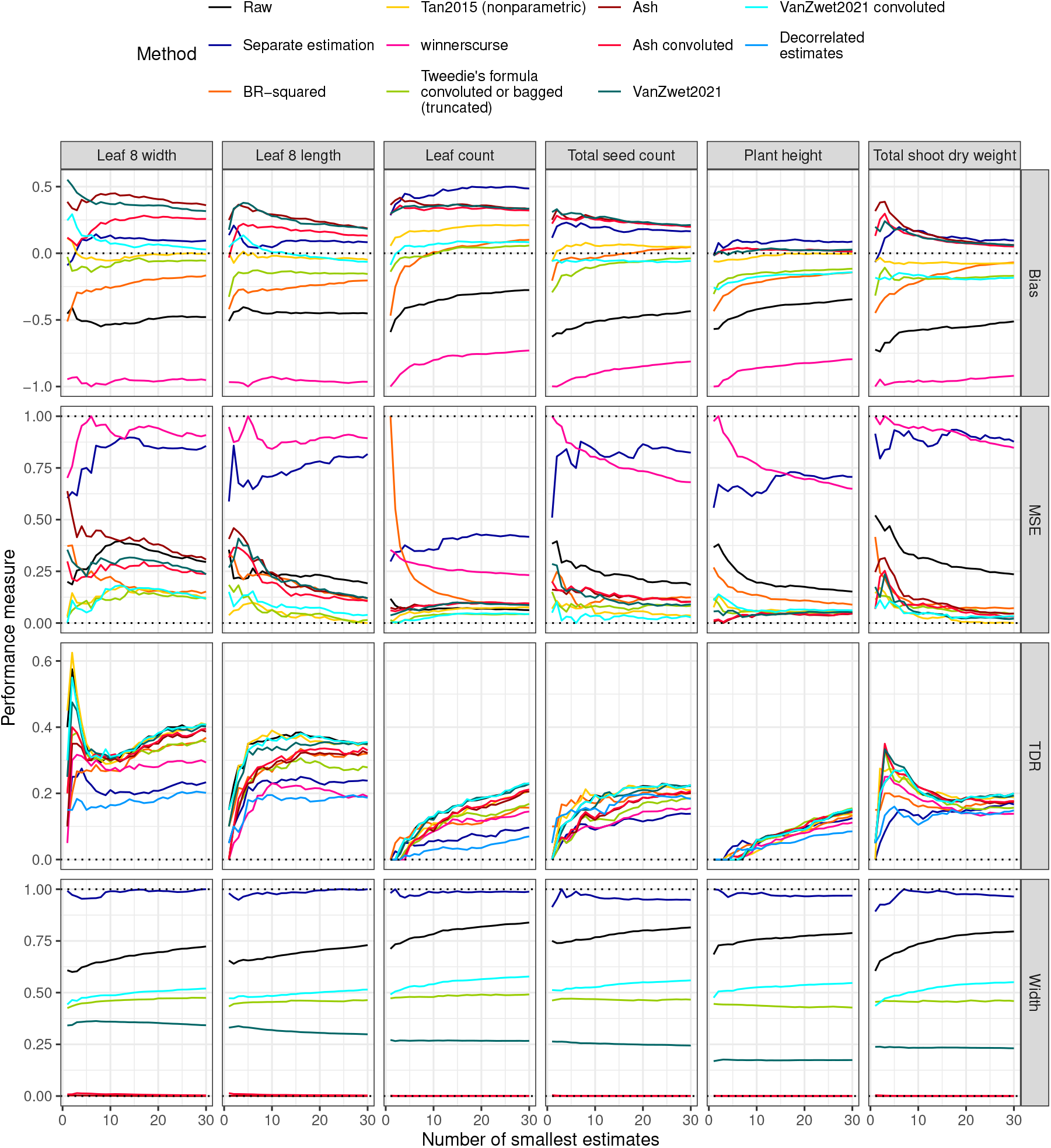
Nonparametric simulation results based on *B. napus* dataset for different methods (colour), performance measures (side panels) and phenotypes (top panels). The x-axis shows the number of top features considered, the y-axis shows the performance measure. The bias (top panels) was scaled to the [-1,1] range and the MSE and width of the confidence interval (second and bottom row) to the [0,1] range per phenotype for legibility (see case study below for an impression of their absolute values). The dotted horizontal lines indicate 0 for all performance measures and 1 for MSE and width of the confidence interval.

## 6. CASE STUDY ON B. NAPUS FIELD TRIAL

A relevant application for selection bias correction in presence of dependence is in predictive modeling with omics data. Often, several prediction models are built for the same outcome variable, including single-feature models and multi-feature models. It then often seems as though the best single-feature model outperforms the multi-feature model in terms of predictive power (Zhou et al., 2021; Cruz et al., 2020). This is also observed in a study by De Meyer et al. (2023), where leaf gene expression and several phenotypes were measured for individual field-grown *Brassica napus* (rapeseed) plants. The aim was to build prediction models based on leaf gene expression in autumn for phenotypes measured the following spring, and estimate the genes’ predictive performance. Supplementary Figure S22 shows the root mean squared error (RMSE) of all single-gene prediction models for predicting six selected outcome phenotypes. Because of common dependence on a single vector of outcome values, these estimates are correlated (see Supplementary Figure S21). In addition, a multi-gene linear model predicting the phenotype concerned from the expression of all genes combined was estimated through elastic net (EN) (see Supplementary Section 3.2 for methodological details). For four out of six phenotypes considered, namely leaf 8 width, leaf count, plant height and total shoot dry weight, the best single-gene model has a lower estimated RMSE than the multi-gene model. This is a plausible result, as even though the multi-gene model is given at least as much information as the single-gene model, its higher model complexity may lead to worse predictive performance compared to the best single-feature model. It is also a desirable result, as a single marker feature is easier to measure than the ensemble of all features. Yet it may also be a case of winner’s curse, so we applied the correction methods investigated to the single-gene RMSEs. Figure 7 shows the smallest raw and corrected single-gene RMSE estimates as well as the multi-gene estimate, with confidence intervals. For leaf count, plant height and total shoot dry weight, the smallest single-gene RMSE point estimate obtained by most selection bias correction methods is no longer smaller than the EN estimate, but for leaf 8 width it still is. Yet also for phenotypes for which the smallest raw RMSE estimate was not smaller than the EN estimate (leaf 8 length and total seed count), the smallest corrected RMSE values are larger than the raw estimate, corroborating our initial suspicion that the latter ones are underestimates. The RMSE estimates of the separate estimation method are mostly the largest with the widest confidence intervals; both observations are due to the smaller sample size on which the RMSEs are estimated. The empirical Bayes credible intervals are narrower than the raw confidence intervals, although often unlikely narrow for the ash method. Often, the feature with the smallest raw RMSE is no longer the smallest one after correction, i.e. the feature ranking has changed (Supplementary Table S1).

**Fig. 7.**
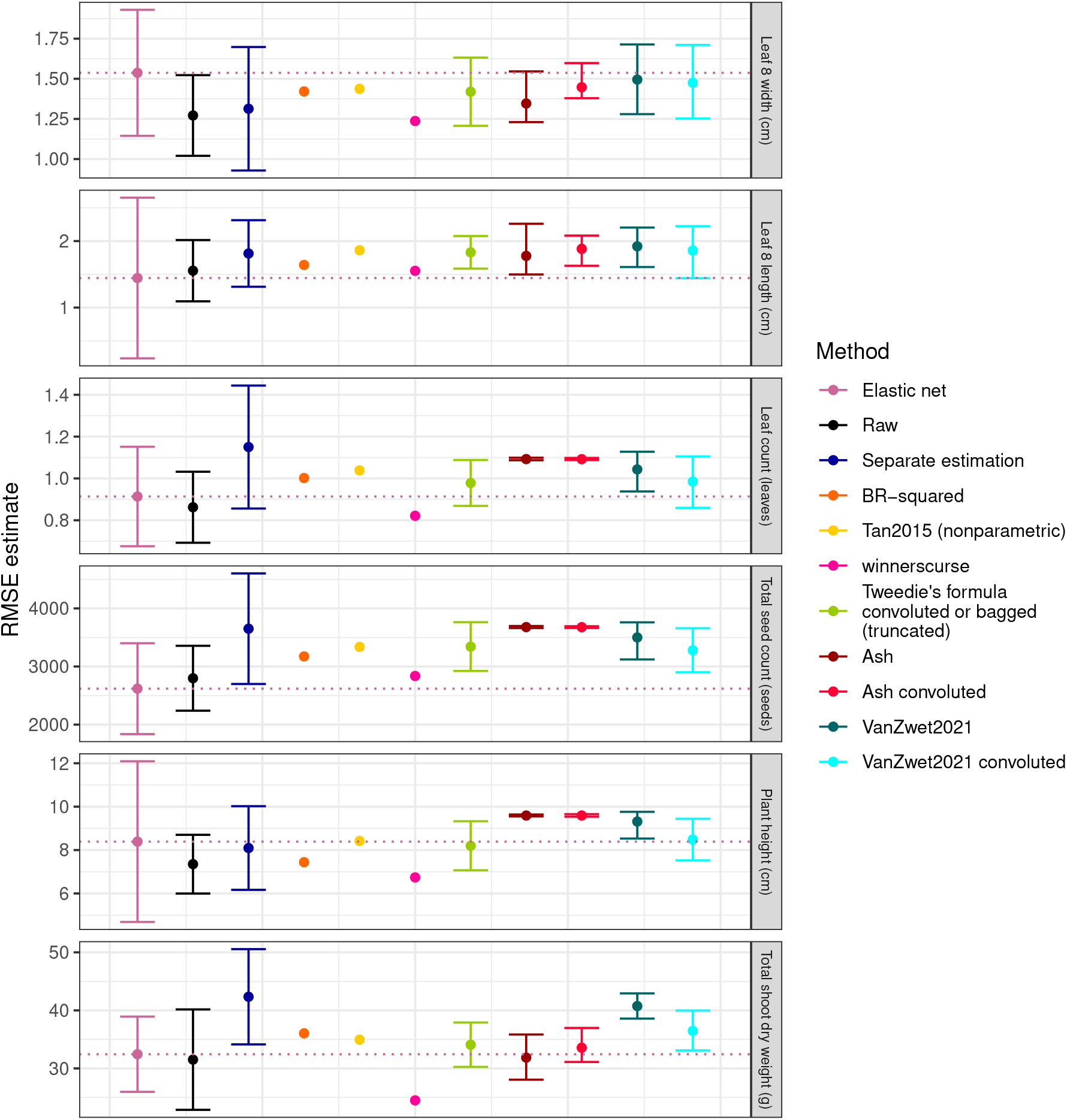
Point estimates of the RMSEs (dots) and corresponding 95% confidence or credible intervals (error bars) for the gene with smallest RMSE for different phenotypes of the *B. napus* dataset (side panels) and correction methods (colour); units are shown between brackets. The RMSE estimates of the multi-gene elastic net models are also shown, the dotted horizontal line corresponds to its point estimate. For the BR-squared and Tan2015 bootstrap methods, no confidence intervals were calculated as this would be too computationally demanding, *winnerscurse* does not provide confidence intervals.

## 7. DISCUSSION

Empirical Bayes methods have previously been found to perform poorly when applied to dependent estimates (Tan et al., 2015; Bigdeli et al., 2016; Ferguson et al., 2013), leading to claims that Tweedie’s formula is invalid in this case (Tan et al., 2015) and to concerns over the performance of ash (Stephens, 2017). Here we prove that these performance issues originate from the random nature of the observed marginal distribution under dependence, causing the width of the true marginal distribution to be underestimated. This underestimation introduces bias into Tweedie’s formula and leads ash and VanZwet2021 to underestimate prior and posterior variability. We provide a remedy in the form of a simple convolution scheme that only requires an estimate of the average correlation between estimators to counter most of the dependence effect. After this modification, all three empirical Bayes methods regained a competitive performance in our simulations.

The collection of most promising estimates in a screening experiment are often biased towards extreme values. Although ubiquitous, this winner’s curse is often ignored, possibly because of lack of awareness or because existing remedies are not widely known. We have benchmarked four classes of correction methods for selection bias in parametric as well as non-parametric simulation: separate estimation, conditional likelihood estimation, three bootstrap methods (BR-squared, Tan2015 and *winnerscurse*) and three empirical Bayes methods (Tweedie’s formula, ash and VanZwet2021), see Table S3 for an overview. Separate estimation is the most drastic solution to selection bias, tackling its root cause by using different parts of the dataset for feature selection and for estimation. It eradicates selection bias, achieves nominal coverage of the confidence intervals and is robust to dependence between estimates. Yet this comes at a high cost in estimator variance and feature ranking accuracy, and it may still be biased under high correlation or when the estimate depends on the sample size, as for prediction models. In addition, the result of the analysis is random as a result of the single data split, which is undesirable. Although this approach may be commendable when the sample size is large, splitting the dataset in two is no substitute for confirmation from an independent experiment. Conditional likelihood is only available for maximum likelihood estimation, but there it greatly reduces selection bias, although its estimates are imprecise and it tends to overcorrect when the estimates are correlated. The three bootstrap methods considered in our comparison, BR-squared, Tan2015 and *winnerscurse*, yielded different results. The BR-squared correction performs poorly in absence of correlation between estimates, but recovers in presence of such correlation. In the original publication, it was only applied to SNPs which are naturally correlated, but it appears not to generalise well to settings without correlation. Also its confidence intervals relying on normality of the estimator do not reach the nominal coverage. The bootstrap method by Tan et al. (2015), on the other hand, is a robust method that effectively combats selection bias both in presence and absence of dependence, and its confidence intervals achieve reasonable coverage, although often still falling short of the nominal level. Presumably because this work is only available as a preprint, it has received little attention in the literature. Both these methods use the average difference of the ranked estimate of a bootstrap sample with a reference quantity to estimate the selection bias. The BR-squared method uses the estimate from a small, random set of out-of-sample observations as reference, whereas the Tan2015 method uses the estimate from the fixed, larger set of observations from the real dataset. The latter option requires fewer parameters to be estimated, is theoretically better justified and performs best in our simulation study, and hence it should be preferred. Both bootstrap methods do not require an estimate of the standard error and make few distributional assumptions. On the downside, these methods do not work for small sample sizes and they can be computationally demanding, especially for the construction of confidence intervals. The *winnerscurse* method is much faster as it only requires a single bootstrap sample and does not rely on the original dataset, yet it demands an estimate of the standard error, has a changeable selection bias correction performance and does not provide a confidence interval. Unlike the other methods considered, empirical Bayes methods were not developed specifically for combatting selection bias of extreme estimates, but simply provide a posterior distribution for all estimates, which is easily coupled to downstream analyses. Yet we found them to be a fast and reliable alternative to bootstrap methods for correcting winner’s curse, if our convolution scheme is used to counter dependence effects. The VanZwet2021 method and ash are preferable when some knowledge on the prior distribution is available; otherwise Tweedie’s formula may be best as it is agnostic to the shape of the prior distribution. On the downside, empirical Bayes methods rely on accurate estimation of estimator variance and densities, and on distributional assumptions, which limit computing time but also make them dependent on high-dimensionality and less robust than bootstrap methods. When a clear null value and an estimator for the standard error are available, conversion of the raw estimates to test statistics can improve ranking of the features and hence also feature selection. In presence of high correlation, also decorrelating the raw estimates or test statistics can improve feature ranking. On the contrary, methods that work well for reducing selection bias can exhibit a poorer feature ranking than the raw estimates, which suggests a bias-variance trade-off.

In a case study on a *B. napus* field trial, we have demonstrated how the apparent superiority of the best single-gene prediction model over the multi-gene prediction model for some phenotypes was most likely a case of selection bias. This result calls for caution in marker selection: a correction for winner’s curse is needed. The effect of winner’s curse decreases as the estimator variability shrinks, or as the differences between the features become more pronounced, as e.g. in our sparse simulation scenario. Yet in practice it is difficult to rule out winner’s curse, so that it is safer to always correct for it. Based on our results, we recommend the Tan2015 bootstrap and empirical Bayes methods, where applicable with our convolution scheme, for this purpose.

## Supporting information

Data, R-code and style file

## Author contributions statement

S.H. conceived and conducted the experiments, S.H., O.T. and S.M. wrote the manuscript. S.M. supervised the study.

## Acknowledgments

We thank Erik van Zwet for pointing us to the work by Azriel and Schwartzman (2015), and for useful discussions.

