## Supplementary material for "The winner’s curse under dependence: repairing empirical Bayes using convoluted densities": Data, R-code and style file: bootCV.R

```
bootCV = function(dat, nOuterFolds, bootReps, cvSplits, nCores = 1, mseOut = FALSE,
                  paramBoot = FALSE, verbose = FALSE, loc = NULL, ...){
    id0 = seq_along(dat$y)
    folds = sample(rep(unFolds <- seq_len(nOuterFolds), length.out = NROW(dat$x)))
    Fit = if(paramBoot) getMeanSigma(dat) else NULL
    bootSplitReps = mclapply(mc.cores = nCores, seq_len(bootReps), function(cvs){
        if(verbose && (cvs %% 10)==0) cat("Bootstrap instance", cvs, "\t")
        #Retain dependence
        if(!paramBoot){
            id = sample(id0, replace = TRUE)
            y = dat$y[id]; x = dat$x[id, , drop = FALSE]
        } else {
            x = dat$x
        }
        t(vapply(FUN.VALUE = double(2+mseOut), seq_len(NCOL(x)), function(j){
            x = x[, j, drop = FALSE]
           if(paramBoot){
               y = rnorm(length(id0), Fit[[j]]$Mean, Fit[[j]]$Sigma) #No dependence
            }
            MSE = mean(vapply(FUN.VALUE = double(1), integer(cvSplits), function(jj){
                folds = sample(folds)
                mean(unlist(sapply(unFolds, function(uf){
                    idTrain = folds!=uf
                    (predLin(x[idTrain,, drop = FALSE], y[idTrain], x[!idTrain,,drop = FALSE ], loc = loc[idTrain,], ...)-y[!idTrain])^2
                })))
            }))
            if(!paramBoot && mseOut){
                idOut = id0[-id]
                xOut = dat$x[idOut,j,drop = FALSE];yOut = dat$y[idOut]; locOut = loc[idOut, ]
                MSEout = mean(vapply(FUN.VALUE = double(1), integer(cvSplits), function(jj){
                    folds = sample(rep(unFoldsOut <- seq_len(nOuterFolds/2), length.out = length(idOut)))
                    mean(unlist(lapply(unFoldsOut, function(uf){
                        idTrain = folds!=uf
                        predTest = predLin(xOut[idTrain,, drop = FALSE], yOut[idTrain], xOut[!idTrain, ], loc = locOut[idTrain,], ...)
                        (predTest-yOut[!idTrain])^2
                    })))
                }))
            } else {MSEout = NULL}
            c("MSE" = MSE, "margVar" = var(y), "MSEout" = MSEout)
        }))
    })
}
varN = function(x) mean((x-mean(x))^2)
bootstrap = function(x, y, id, Mean, Sigma, paramBoot){
    if(paramBoot){
        y = rnorm(length(y), Mean, Sigma)
    } else {
        id = sample(id, replace = TRUE)
        y = y[id];x=x[id, ,drop = FALSE]
    }
    return(list("x" = x, "y" = y, "id" = id))
}
```
