## Supplementary material for "The winner’s curse under dependence: repairing empirical Bayes using convoluted densities": Data, R-code and style file: bootNonParam.R

```
bootNonParam = function(mat, bootReps, ciReturnBoot){
    n = nrow(mat);id = seq_len(n)
    lapply(integer(bootReps), function(b){
        id0 = sample(id, replace = TRUE)
        estsIn = colMeans(matIn <- mat[id0,])
        estsOut = colMeans(matOut <- mat[-id0,])
        out = cbind("estsIn" = estsIn, "estsOut" = estsOut,
                    "varEstsIn" = rowMeans((t(matIn) - estsIn)^2)/(n - 1),
                    "varEstsOut" = rowMeans((t(matOut) - estsOut)^2)/(n - 1))
        outIn = if(ciReturnBoot){
            vapply(seq_len(bootReps), FUN.VALUE = matrix(0, ncol(mat), 4), function(b){
                idIn = sample(id, replace = TRUE)
                estsIn = colMeans(matIn <- mat[id0,][idIn,])
                estsOut = colMeans(matOut <- mat[id0,][-idIn,])
                cbind("estsIn" = estsIn, "estsOut" = estsOut,
                      "varEstsIn" = rowMeans((t(matIn) - estsIn)^2)/(n - 1),
                      "varEstsOut" = rowMeans((t(matOut) - estsOut)^2)/(n - 1))
            })
        }
        list("out" = out, "outIn" = outIn)
    })
}
```
