## Supplementary material for "The winner’s curse under dependence: repairing empirical Bayes using convoluted densities": Data, R-code and style file: bootParam.R

```
bootParam = function(mat, bootReps, rawEsts, ciReturnBoot, method = "eigen"){
    n = nrow(mat)
    paramBootMat = mvtnorm::rmvnorm(n = n*bootReps, mean = rawEsts, sigma = cov(mat), method = method)
    lapply(seq_len(bootReps), function(i){
        estsIn = colMeans(matIn <- paramBootMat[seq_len(n)+(i-1)*n,])
        out = cbind("estsIn" = estsIn,
                    "varEstsIn" = rowMeans((t(matIn) - estsIn)^2)/(n - 1))
        outIn = if(ciReturnBoot){
            paramBootMatIn = mvtnorm::rmvnorm(n = n*bootReps, mean = out[, 'estsIn'], sigma = cov(matIn), method = method)
            vapply(seq_len(bootReps), FUN.VALUE = matrix(0, ncol(mat), 2), function(b){
                estsIn = colMeans(matIn <- paramBootMatIn[seq_len(n)+(b-1)*n,])
                cbind("estsIn" = estsIn,
                      "varEstsIn" = rowMeans((t(matIn) - estsIn)^2)/(n - 1))
            })
        }
        list("out" = out, "outIn" = outIn)
    })
}
```
