## Supplementary material for "The winner’s curse under dependence: repairing empirical Bayes using convoluted densities": Data, R-code and style file: BR_ss_edit.R

```
BR_ss_edit = function (summary_data, seed_opt = FALSE, seed = 1998, allowInflate)
{
    stopifnot(all(c("rsid", "beta", "se") %in% names(summary_data)))
    stopifnot(!all(is.na(summary_data$rsid)) && !all(is.na(summary_data$beta)) &&
                  !all(is.na(summary_data$se)))
    stopifnot(nrow(summary_data) > 5)
    stopifnot(is.numeric(summary_data$beta) && is.numeric(summary_data$se))
    stopifnot(!any(duplicated(summary_data$rsid)))
    summary_data <- dplyr::arrange(summary_data, dplyr::desc((summary_data$beta/summary_data$se)))
    N <- nrow(summary_data)
    if (seed_opt == TRUE) {
        set.seed(seed)
    }
    beta_boot <- matrix(stats::rnorm(1 * N, mean = rep(summary_data$beta,
                                                       1), sd = rep(summary_data$se, 1)), nrow = N, ncol = 1,
                        byrow = FALSE)
    beta_mat <- matrix(rep(summary_data$beta, 1), nrow = N,
                       ncol = 1, byrow = FALSE)
    se_mat <- matrix(rep(summary_data$se, 1), nrow = N, ncol = 1,
                     byrow = FALSE)
    beta_oob <- beta_mat
    ordering <- apply(beta_boot/se_mat, 2, order, decreasing = TRUE)
    bias_correct <- matrix(nrow = N, ncol = 1)
    bias_correct[, 1] <- (beta_boot[ordering[, 1], 1] - beta_oob[ordering[,
                                                                          1], 1])/summary_data$se[ordering[, 1]]
    z <- summary_data$beta/summary_data$se
    bias_correct <- stats::predict(stats::smooth.spline(z, bias_correct)$fit,
                                   z)$y
    beta_BR_ss <- summary_data$beta - summary_data$se * bias_correct[rank(-1 *
                                                                              summary_data$beta/summary_data$se)]
    beta_BR_ss[sign(beta_BR_ss) != sign(summary_data$beta)] <- 0
    summary_data <- cbind(summary_data, beta_BR_ss)
    if(!allowInflate){
        for (i in 1:N) {
            if (abs(summary_data$beta[i]) < abs(summary_data$beta_BR_ss[i])) {
                summary_data$beta_BR_ss[i] <- summary_data$beta[i]
            }
        }
    }
    #Leave out this correction for RMSE without clear null value
    summary_data <- dplyr::arrange(summary_data, dplyr::desc(abs(summary_data$beta/summary_data$se)))
    return(summary_data)
}
```
