## Supplementary material for "The winner’s curse under dependence: repairing empirical Bayes using convoluted densities": Data, R-code and style file: buildCredInt.R

```
#Build the credible interval for Tweedie's formula
buildCredInt = function(tweeEsts, vars, quants, addVarDens = FALSE, bootEsts = NULL, bootReps = 1e2, varVar = FALSE, n, ...){
    VarsNew = vars*(1 + vars*tweeEsts["densEstsLogDeriv2",])
    VarsNew[VarsNew<0] = vars[VarsNew<0]
    Ests = tweeEsts["tweedieCorEsts",]
    if(addVarDens){
        #add variance due to density estimation
        if(is.null(bootEsts)){
            #Poisson approximation: probably not very reliable since bootstraps are not independent
            degree = ncol(tweeEsts$vcov)-1L
            Polymatrix = seq_len(degree)*t(cbind(1, polym(Ests, degree = degree, raw = TRUE)[, -degree]))
            zVec = outer(Ests, seq_len(degree), FUN = "^")
            Vcov = tweeEsts$vcov[-1,-1]
            VarDens = diag(crossprod(Polymatrix, Vcov %*% Polymatrix))
        } else {
            bootDensEsts = vapply(integer(bootReps), FUN.VALUE = Ests, function(i){
                id = sample(length(Ests), replace = TRUE)
                deriv1 = LindseysMethod(zObs = Ests, z = c(bootEsts[, id]), ...)$deriv1
            })
            VarDens = apply(bootDensEsts, 1, var)
        }
        if(varVar){#Include uncertainty on the variance
            varVars = vars^2*2/(n-1)
            varDensVar = varVars*VarDens + VarDens*vars^2 + varVars*VarDens^2
            #Single features has little impact on density estimation => Assume independence
        } else {
            varDensVar = VarDens*vars^2 #For MSE: ignore this variance
        }
        VarsNew = VarsNew + varDensVar
    }
    Ests + outer(sqrt(VarsNew), quants)
}
```
