## Supplementary material for "The winner’s curse under dependence: repairing empirical Bayes using convoluted densities": Data, R-code and style file: condML.R

```
# The conditional maximum likelihood following Zollner2007
condML = function(mu, dat, thresh, lower.tail, Sum = TRUE){
    sd = sqrt(length(dat)/(length(dat)-1)*mean((dat-mu)^2))
    tmp = sum(dnorm(dat, mean = mu, sd = sd, log = TRUE)) -
       pnorm(thresh-mu, sd = sd/sqrt(length(dat)), lower.tail = lower.tail, log.p = TRUE)
    #"The numerator is maximized at the naive penetrance estimates."
    -tmp
}
estCondML = function(init,  dat, thresh, lower.tail){
    optim(init, fn = condML, dat = dat, thresh = thresh,
          lower.tail = lower.tail, method = "Brent", lower = min(dat), upper = max(dat))$par
}
findConfReg = function(dat, se, thresh, lower.tail, quants, condMLest, numSEs = 5, numPoints = 2e3, sigLevel = 0.05, Df = 2){
    #Evaluate densities over a grid
    zGrid = seq(condMLest-numSEs*se, condMLest+numSEs*se, length.out = numPoints)
    evalDens = vapply(zGrid, FUN.VALUE = double(1), condML,
                      dat = dat, thresh = thresh, lower.tail = lower.tail)
    #Zollner2007 and Zhong2008: Chi-square
    condMLopt = condML(mu = condMLest, dat = dat, thresh = thresh, lower.tail = lower.tail)
    diffLL =  evalDens - condMLopt #Difference in likelihoods
    chisqPerc = qchisq(p = 1-sigLevel, df = Df) #The 95th percentile, two parameter model being tested for (mean and variance)
    confRegChisq = range(zGrid[which(2*diffLL < chisqPerc)])
    names(confRegChisq) = paste0(names(quants), "chisq")
    return(confRegChisq)
}
#Ghosh2008 talk about quantiles of the conditional likelihood, but is this not even a proper density
condMLstat = function(mu, z, thresh, lower.tail){
     - dnorm(z - mu, log = TRUE) + pnorm(thresh - mu, lower.tail = lower.tail, log.p = TRUE)
}
estCondMLstat = function(init, thresh, lower.tail, z, lower, upper){
    optim(init, fn = condMLstat, thresh = thresh, z = z, lower.tail = lower.tail,
          method = "Brent", lower = lower, upper = upper)$par
}
findConfRegStat = function(raw, thresh, lower.tail, quants, condMLest, numSEs = 5, numPoints = 2e3, sigLevel = 0.05, Df = 1){
    #Evaluate densities over a grid
    zGrid = seq(condMLest-numSEs, condMLest+numSEs, length.out = numPoints)
    evalDens = vapply(zGrid, FUN.VALUE = double(1), condMLstat,
                      z = raw, thresh = thresh, lower.tail = lower.tail)
    #Zollner2007 and Zhong2008: Chi-square
    condMLopt = condMLstat(mu = condMLest, z = raw, thresh = thresh, lower.tail = lower.tail)
    diffLL =  evalDens - condMLopt #Difference in likelihoods
    chisqPerc = qchisq(p = 1-sigLevel, df = Df) #The 95th percentile, one parameter model being tested for
    confRegChisq = range(zGrid[which(2*diffLL < chisqPerc)])
    names(confRegChisq) = paste0(names(quants), "chisq")
    return(confRegChisq)
}
corrCondML = function(raw, vars, dat, extremes, ciReturn, quants = c(0.025, 0.975), testStatistic = FALSE){
    p = length(raw)
    names(quants) = quants
    sortRaw = sort(raw)
    if(testStatistic){
        lapply(extremes, function(e){
            lapply(c("small" = FALSE, "large" =  TRUE), function(t){
                ext = if(t) sortRaw[p-e+1] else sortRaw[e]
                id = which(if(t) raw >= ext else raw <= ext);names(id) = names(raw)[id]
                condMLests = vapply(id, FUN.VALUE = double(1), function(i){
                    estCondMLstat(init = raw[i], thresh = ext, lower.tail = !t, z = raw[i],
                                  lower = if(t) 0 else raw[i], upper = if(t) raw[i] else 0)
                })
                condMLcr = if(ciReturn){
                    vapply(seq_along(id), FUN.VALUE = double(2), function(i){
                        findConfRegStat(raw = raw[id[i]], thresh = ext, lower.tail = !t,
                                    quants = quants, condMLest = condMLests[i])
                    })
                }
                cbind("condMLests" = condMLests, if(ciReturn){t(condMLcr)} else NULL)
            })
        })
    } else {
        ses = sqrt(vars) #Standard errors
        lapply(extremes, function(e){
            lapply(c("small" = FALSE, "large" =  TRUE), function(t){
                ext = if(t) sortRaw[p-e+1] else sortRaw[e]
                id = which(if(t) raw >= ext else raw <= ext);names(id) = names(raw)[id]
                condMLests = vapply(id, FUN.VALUE = double(1), function(i){
                    estCondML(init = mean(dat[, i]), dat = dat[, i], thresh = ext, lower.tail = !t)
                })
                condMLcr = if(ciReturn){
                    vapply(seq_along(id), FUN.VALUE = double(2), function(i){
                        findConfReg(dat = dat[, id[i]], se = ses[id[i]], thresh = ext, lower.tail = !t,
                               quants = quants, condMLest = condMLests[i])
                    })
                }
                cbind("condMLests" = condMLests, if(ciReturn){t(condMLcr)} else NULL)
            })
        })
    }
}
```
