## Supplementary material for "The winner’s curse under dependence: repairing empirical Bayes using convoluted densities": Data, R-code and style file: densNM.R

```
anaDerivLog = function(z, mu ,sd){
    sNorm = sum(dnorm(z,mu, sd))
    sNorm2 = sum(dnorm(z, mu, sd)*(mu-z)/sd^2)
    deriv1 = sNorm2/sNorm
    deriv2 = sum(dnorm(z,mu, sd)*(((mu-z)/sd^2)^2-1/sd^2))/sNorm - (sNorm2/sNorm)^2
    c("deriv1" = deriv1, "deriv2" = deriv2)
}
evalMixNorm = function(z, mu, sd){
    p = length(mu)
    rowMeans(vapply(seq_len(p), FUN.VALUE = z, function(j){
        dnorm(z, mu[j], sd[j])
    }))
}
```
