## Supplementary material for "The winner’s curse under dependence: repairing empirical Bayes using convoluted densities": Data, R-code and style file: errorLarge.R

```
errorLarge = function(ests, trueVec, ext, p, ests2rank = ests, power = 1, outSEs = NULL){
    rankMat = rank(ests2rank)
    c("small" = mean((ests[rankMat <= ext]- trueVec[rankMat <= ext])^power - (if(!is.null(outSEs) && power == 2) outSEs[rankMat <= ext]^2 else 0)),
      "large" = mean((ests[rankMat > p-ext] - trueVec[rankMat > p-ext])^power -  (if(!is.null(outSEs) && power == 2) outSEs[rankMat > p-ext]^2 else 0)))
}
runningTDP = function(ests, trueRanks, ext, p, ests2rank = ests){
    rankObs = rank(ests2rank)
    c("small" = mean(names(ests)[rankObs <= ext] %in% names(trueRanks)[trueRanks <= ext]),
      "large" = mean(names(ests)[rankObs > p-ext] %in% names(trueRanks)[trueRanks > p-ext]))
}
```
