## Supplementary material for "The winner’s curse under dependence: repairing empirical Bayes using convoluted densities": Data, R-code and style file: evalFitLogDeriv.R

```
#Evaluate the fit for the derivative of the logarithm
evalFitLogDeriv = function(Coef, z, expansion){
    degree = length(Coef)-1
    if(missing(expansion)){
        expansion <- polym(z, degree = degree, raw = TRUE)
    }
    c(cbind(1, expansion[, -degree]) %*% (Coef[-1]*seq_len(degree)))
}
evalFitLog = function(Coef, z, expansion){
    degree = length(Coef)-1
    if(missing(expansion)){
        expansion <- polym(z, degree = degree, raw = TRUE)
    }
    c(cbind(1, expansion) %*% Coef)
}
```
