## Supplementary material for "The winner’s curse under dependence: repairing empirical Bayes using convoluted densities": Data, R-code and style file: evalZwetDens.R

```
#Evaluate the mixture density of the zwet normal mixture
evalZwetDens = function(dens, z){
    dens = rowSums(apply(dens, 2, function(x){
        dnorm(z, x["mean"], x["sigma"])*x["probs"]
    }))
}
#Get the overal variance
zwetOverallVar = function(dens){
    ovMean = sum(dens["mean", ]*dens["probs", ])
    meanVar = sum((dens["mean",]-ovMean)^2*dens["probs", ])
    resVar = sum(dens["sigma",]^2*dens["probs", ])
    meanVar + resVar
}
```
