## Supplementary material for "The winner’s curse under dependence: repairing empirical Bayes using convoluted densities": Data, R-code and style file: FayeCorrect.R

```
#Apply the Faye2011 correction
FayeCorrect = function(rawEsts, bootEstsIn, bootEstsOut, ciReturn = FALSE, fullBootObj, zQuants, testStatistic = FALSE){
    rankRaw = rank(rawEsts)
    rankIn = t(apply(bootEstsIn, 1, rank)) # All ranking is based on in-bootstrap samples
    # "we apply the exact same selection criteria as in the original sample" = thresholding in our case, always yields hits
    orderIn = apply(bootEstsIn, 1, order)
    seqBoot <- seq_len(nBoots <- nrow(bootEstsIn))
    betaCor = vapply(FUN.VALUE = double(nBoots),
                     seq_along(rawEsts), function(k){ #Loop over ranks
        bootEstInK = bootEstsIn[idri <- rankIn==k]
        bootEstOutK = bootEstsOut[idri] #Same feature for out of sample error
        sigma2Di = var(bootEstInK)
        sigmaDEi = cov(bootEstInK, bootEstOutK) #Is this what they mean? Covariance over the bootstraps
        sigRat = sigmaDEi/sigma2Di
        vapply(FUN.VALUE = double(1), seqBoot, function(i){
            bootEstOut = bootEstsOut[i, id <- orderIn[k, i]] #Use k-ranked SNP in bootstrap sample. This means IN!
            #Remember orderIn[k, i] = which(rankIn[i,]==k)
            bootEstIn = bootEstsIn[i, id]
            betaNik = rawEsts[id]
            bootEstIn - (bootEstOut - sigRat*(bootEstIn-betaNik))
        })
    })
    out = rawEsts - colMeans(betaCor)[rankRaw]
    ciFaye = if(ciReturn){
        outIn = vapply(FUN.VALUE = rawEsts, seq_len(dim(fullBootObj[[1]]$outIn)[3]), function(i){
            FayeCorrect(rawEsts = bootEstsIn[i,],
                        bootEstsIn = t(fullBootObj[[i]]$outIn[,"estsIn",]/(if(testStatistic) sqrt(fullBootObj[[i]]$outIn[,"varEstsIn",]) else 1)),
                    bootEstsOut = t(fullBootObj[[i]]$outIn[,"estsOut",]/(if(testStatistic) sqrt(fullBootObj[[i]]$outIn[,"varEstsOut",]) else 1)))$est[order(bootEstsIn[i,])]
        })
        sdK = apply(outIn, 1, sd) #Standard deviation of k-ranked estimates
        ci = out + outer(sdK[order(out)], zQuants)
        names(ci) = names(out)
        ci
    }
    return(list("est" = out, "ci" = ciFaye))
}
```
