## Supplementary material for "The winner’s curse under dependence: repairing empirical Bayes using convoluted densities": Data, R-code and style file: filterFeat.R

```
#Returns indices of features to filter out
filterFeat = function(mat, criterion, out, spearThreshP = 0.01){
    if(criterion == "spearman"){
        spearCorsP = apply(mat, 2, function(x) cor.test(x, out, method = "spearman")$p.value)
        which(p.adjust(spearCorsP, method = "BH")>spearThreshP)
    }
}
filterFun = function(mat, out, ...){
    id = filterFeat(mat, out, criterion = 'spearman', ...)
    mat[, -id, drop = FALSE]
}
```
