## Supplementary material for "The winner’s curse under dependence: repairing empirical Bayes using convoluted densities": Data, R-code and style file: forde.R

```
# Bootstrap method from Forde2023
forde = function(estimates, varsEstimates, allowInflate = FALSE){
    df = data.frame("rsid" = names(estimates), "beta" = estimates,
                    "se" = sqrt(varsEstimates))
    BRres = BR_ss_edit(df, allowInflate = allowInflate)
    BRres[names(estimates),]$beta_BR_ss #Order is changed internally
}
```
