## Supplementary material for "The winner’s curse under dependence: repairing empirical Bayes using convoluted densities": Data, R-code and style file: genDat.R

```
genDat = function(n, p, sdY = 1, betas = integer(p), sdX = rep(1, p), ...){
    x = matrix(rnorm(n*p, sd = rep(sdX, each = n)), n, p)
    y = rnorm(n, mean = x %*% betas, sd = sdY)
    list("x" =  x, "y" = y)
}
```
