## Supplementary material for "The winner’s curse under dependence: repairing empirical Bayes using convoluted densities": Data, R-code and style file: getConvAsh.R

```
#Build convolution density for ash
getConvAsh = function(raws, ses, alpha1, numSamAsh = 10, ashObj, mixcompdist = "uniform", ...){
    if(alpha1 > 0){
        p = length(raws)
        samConvDens = rnorm(p*numSamAsh, raws, ses*sqrt(alpha1))
        ashObjConvTmp = ash(samConvDens, repVars <- rep(ses, length.out = p*numSamAsh),
                            method = "shrink", mode = "estimate", mixcompdist = mixcompdist, ...)
        ash(raws, ses, method = "shrink", g = ashObjConvTmp$fitted_g, fixg = TRUE, ...)
    } else {ashObj}
}
```
