## Supplementary material for "The winner’s curse under dependence: repairing empirical Bayes using convoluted densities": Data, R-code and style file: getCorrectedEsts.R

```
#'Extract bias corrected matrices
getCorrectedEsts = function(resultsList, breaksNorm = 4e1, breaksBoot = 2e2, zOracle = NULL, quants = c(0.025, 0.975),
                            RMSE = FALSE, varOracle = NULL, nCores = 1, Quant = 0.5, ciReturn = FALSE, numSamAsh = 10,...){
    names(quants) = quants;zQuants = qnorm(quants)
    mclapply(mc.cores = nCores, seq_along(resultsList), function(tx){
        cat(tx, "\t")
        rawEst = t(sapply(resultsList[[tx]]$seEsts, function(z) z["Bates", ]))
        raws = rawEst[, "MSEhat"];p = length(raws)
        names(raws) = seq_along(raws)
        bootEstsNonparam = sapply(resultsList[[tx]]$bootEsts, function(z) z[,"MSE"])
        bootEstsParam = sapply(resultsList[[tx]]$bootEstsParam, function(z) z[,"MSE"])
        bootEstOut = sapply(resultsList[[tx]]$bootEsts, function(z) z[,"MSEout"])
        if(RMSE){
            raws = sqrt(raws)
            rawEst[, "SE"] = rawEst[, "SE"]/(2*raws)
            bootEstsNonparam = sqrt(bootEstsNonparam);bootEstsParam = sqrt(bootEstsParam);bootEstOut = sqrt(bootEstOut)
        }
        VarsBoot = apply(bootEstsNonparam, 1, var);VarsBootParam = apply(bootEstsParam, 1, var)
        rawsStand = raws/rawEst[, "SE"]
        # Do not reestimate standard errors in the bootstrap, too much computation
        alpha1 = getAlpha1(t(bootEstsNonparam))
        tweedieEst = tweedieForm(estimates = raws, varsEstimates = Vars <- rawEst[, "SE"]^2,
                                  analytical = FALSE)["tweedieCorEsts",]
        tq = quantile(raws, Quant)
        tweedieEstsTrunc = tweedieEst;tweedieEstsTrunc[raws > tq] = raws[raws > tq]
        tweedieConvEstsList = tweedieForm(estimates = raws, varsEstimates = Vars,
                                       alpha1 = alpha1)
        tweedieConvEsts = tweedieConvEstsList["tweedieCorEsts",]
        tweedieConvEstsBootList = tweedieForm(estimates = raws, varsEstimates = VarsBoot,
                                          alpha1 = alpha1)
        tweedieConvEstsBoot = tweedieConvEstsBootList["tweedieCorEsts",]
        tweedieConvEstsTrunc = tweedieConvEstsList;tweedieConvEstsTrunc["tweedieCorEsts",][raws > tq] = raws[raws > tq]
        tweedieConvEstsTruncBoot = tweedieConvEstsBootList;tweedieConvEstsTruncBoot["tweedieCorEsts",][raws > tq] = raws[raws > tq]
        tweedieEstStand = rawEst[, "SE"]*tweedieForm(estimates = rawsStand, varsEstimates = 1, breaks = 40, analytical = FALSE)["tweedieCorEsts",]
        tweedieConvEstsStand = rawEst[, "SE"]*tweedieForm(estimates = rawsStand, varsEstimates = 1, breaks = 40, alpha1 = alpha1)["tweedieCorEsts",]
        TanCorrectEst = TanCorrect(raws, t(bootEstsNonparam), ciReturn = FALSE, quants = quants)
        TanCorrectEstParam = TanCorrect(raws, t(bootEstsParam), ciReturn = FALSE, quants = quants)
        FayeCorrectEst = FayeCorrect(raws, t(bootEstsNonparam), t(bootEstOut), ciReturn = FALSE)
        #ash
        ashObj = ash(raws, s<-sqrt(Vars), method = "shrink", mode = "estimate")
        bootCorrectForde = forde(raws, rawEst[, "SE"]^2, allowInflate = TRUE)
        zwetRes = zwetWrapper(z = (raws - (mr <- mean(raws)))/s, s = s)
        zwetResConv = if(alpha1 < 0){
            zwetRes
        }else {
            zEsts = rnorm(p*10, raws, s*sqrt(alpha1))
            zwetWrapper(z = (raws - mr)/s, s = s, (zEsts-mr)/s)
        }
        ashObjConv = getConvAsh(raws, sqrt(Vars), alpha1 = alpha1, numSamAsh = numSamAsh, ashObj = ashObj)
        #cat scores
        rawsCatBoot = crossprod.powcor.shrink(t(bootEstsNonparam), raws-mean(raws), alpha = -1/2, verbose = FALSE)+mean(raws)
        if(is.null(zOracle)){
            tweedieEstOracleDens = tweedieEstOracleDensStand = NULL
            } else {
                estForDensOracle = if(RMSE) sqrt(zOracle) else zOracle
                estForDensOracleStand = if(RMSE) sqrt(zOracle)/sqrt(varOracle) else zOracle/sqrt(varOracle)
                tweedieEstOracleDensList = tweedieForm(estimates = raws, varsEstimates = Vars,
                            breaks = breaksBoot, estimatesForDens = estForDensOracle, analytical = FALSE)
                tweedieEstOracleDens = tweedieEstOracleDensList["tweedieCorEsts",]
                tweedieEstOracleDensStand = rawEst[, "SE"]*tweedieForm(estimates = rawsStand, analytical = FALSE, varsEstimates = 1,
                                                breaks = breaksBoot, estimatesForDens = estForDensOracleStand)["tweedieCorEsts",]
             }
        if(is.null(varOracle)) {
            tweedieEstOracleVar = tweedieEstOracleVarParam = tweedieEstOracleVarStand = tweedieEstOracleVarParamStand = NULL
        } else {
            tweedieEstOracleVarList = tweedieForm(estimates = raws, varsEstimates = varOracle,
                                               breaks = breaksBoot, alpha1 = alpha1)
            tweedieEstOracleVar = tweedieEstOracleVarList["tweedieCorEsts",]
            tweedieEstOracleVarStand = rawEst[, "SE"]*tweedieForm(estimates = rawsStandOracle <- raws/sqrt(varOracle),
                                               varsEstimates = 1, breaks = breaksBoot, alpha1 = alpha1)["tweedieCorEsts",]
        }
        if(is.null(zOracle) || is.null(varOracle)) {
            tweedieEstOracleDensVar = tweedieEstOracleDensVarStand = NULL
            } else {
                tweedieEstOracleDensVarList = tweedieEstOracleDensVar = tweedieForm(estimates = raws,
                            varsEstimates = varOracle, breaks = breaksBoot, analytical = FALSE,
                            estimatesForDens = estForDensOracle)
                tweedieEstOracleDensVar = tweedieEstOracleDensVarList["tweedieCorEsts",]
                tweedieEstOracleDensVarStand = tweedieForm(estimates = rawsStandOracle, varsEstimates = 1, analytical = FALSE,
                                                           breaks = breaksBoot, estimatesForDens = estForDensOracleStand)["tweedieCorEsts",]
            }
        outSplit = cbind("estInSplit" = sapply(resultsList[[tx]]$estInSplit, function(x) mean(unlist(x))),
                         "estOutSplit" = sapply(resultsList[[tx]]$estOutSplit, function(z) z["Bates", "MSEhat"]))
        outSE = sapply(resultsList[[tx]]$estOutSplit, function(z) z["Bates", "SE"])
        if(RMSE){
            outSplit = sqrt(outSplit); outSE = outSE/(2*outSplit[, "estOutSplit"])
        }
        resMat =  cbind("rawEst" = raws, "tweedieEst" = tweedieEst, "tweedieEstsTrunc" = tweedieEstsTrunc,
                        "tweedieEstStand" = tweedieEstStand, "tweedieConvEsts" = tweedieConvEsts,
                     "tweedieConvEstsTrunc" = tweedieConvEstsTrunc["tweedieCorEsts",], "tweedieConvEstsBoot" = tweedieConvEstsBoot,
                     "tweedieConvEstsTruncBoot" = tweedieConvEstsTruncBoot["tweedieCorEsts",],
                     "tweedieConvEstsStand" = tweedieConvEstsStand,
              "TanCorrectEst" = TanCorrectEst$est, "TanCorrectEstParam" = TanCorrectEstParam$est, "FayeCorrectEst" = FayeCorrectEst$est,
              "tweedieEstOracleDens" = tweedieEstOracleDens, "tweedieEstOracleVar" = tweedieEstOracleVar,
             "tweedieEstOracleDensVar" = tweedieEstOracleDensVar,
              "tweedieEstOracleDensStand" = tweedieEstOracleDensStand, "tweedieEstOracleVarStand" = tweedieEstOracleVarStand,
              "tweedieEstOracleDensVarStand" = tweedieEstOracleDensVarStand,
             outSplit, "ash" = ashObj$result$PosteriorMean, "ashConv" = ashObjConv$result$PosteriorMean,
              "rawsCatBoot" = c(rawsCatBoot), "bootCorrectForde" = bootCorrectForde,
             "VanZwet2021" = zwetRes$res$betaHats+mr, "VanZwet2021conv" = zwetResConv$res$betaHats+mr)
        ciList = if(ciReturn){
            list("rawEst" = raws + outer(sqrt(Vars), zQuants),
                 "estOutSplit" = outSplit[, "estOutSplit"] + outer(outSE, zQuants),
                 "tweedieConvEsts" = buildCredInt(tweedieConvEstsList, n = n, Vars, zQuants),
                 "tweedieConvEstsBoot" = buildCredInt(tweedieConvEstsList, n = n, VarsBoot, zQuants),
                 "tweedieConvEstsTrunc" = buildCredInt(tweedieConvEstsTrunc, n = n, Vars, zQuants),
                 "tweedieEstOracleVar" = buildCredInt(tweedieEstOracleVarList, n = n, varOracle, zQuants),
                 "tweedieEstOracleDens" = buildCredInt(tweedieEstOracleDensList, n = n, Vars,
                                                       zQuants, bootEsts = estForDensOracle, addVarDens = FALSE),
                 "tweedieEstOracleDensVar" = buildCredInt(tweedieConvEstsList, n = n, varOracle,
                                                          zQuants, bootEsts = estForDensOracle, addVarDens = FALSE),
                 "ash" = ashci(ashObj, level = nomCoverage <- 1-quants[1]*2, trace = FALSE),
                 "ashConv" = ashci(ashObjConv, level = nomCoverage, trace = FALSE),
                 "rawsCatBoot" = c(rawsCatBoot) + outer(sqrt(Vars), zQuants),
                 "VanZwet2021" = zwetRes$confInts + mr, "VanZwet2021conv" = zwetResConv$confInts + mr
            )
        }
       return(list("resMat" = resMat, "ciList" = ciList))
    })
}
```
