## Supplementary material for "The winner’s curse under dependence: repairing empirical Bayes using convoluted densities": Data, R-code and style file: getCorrectedEstsIndep.R

```
#'Extract bias corrected matrices
getCorrectedEstsIndep = function(resultsList, breaksNorm = 1e2, breaksBoot = 1e3, zOracle = NULL,
                            RMSE = FALSE, varOracle = NULL, Quant = 0.5, nCores = 1, ...){
    nullIndepEstsMat = mclapply(mc.cores = nCores, seq_along(resultsList), function(tx){
        cat(tx, "\t")
        rawEst = t(sapply(resultsList[[tx]], function(z) z$seEsts))
        p = nrow(rawEst)
        bootEsts = sapply(resultsList[[tx]], function(z) sapply(z$bootEsts, function(y) y[,"MSE"]))
        alpha1 = getAlpha1(bootEsts)
        bootEstsParam = sapply(resultsList[[tx]], function(z) sapply(z$bootEstsParam, function(y) y[,"MSE"]))
        MSEout = sapply(resultsList[[tx]], function(z) sapply(z$bootEsts, function(y) y[,"MSEout"]))
        if(RMSE){
            rawEst[, "MSEhat"] = sqrt(rawEst[, "MSEhat"])
            rawEst[, "SE"] = rawEst[, "SE"]/(2*rawEst[, "MSEhat"])
            bootEsts = sqrt(bootEsts);MSEout = sqrt(MSEout); bootEstsParam = sqrt(bootEstsParam)
        }
        raws = rawEst[, "MSEhat"];names(raws) = seq_along(raws)
        tweedieEst = tweedieForm(estimates = rawEst[, "MSEhat"], varsEstimates = Vars<- rawEst[, "SE"]^2,
                                  analytical = FALSE)["tweedieCorEsts", ]
        tweedieConvEsts = tweedieForm(estimates = rawEst[, "MSEhat"], alpha1 = alpha1, varsEstimates = Vars, estimatesForDens = c(bootEsts))["tweedieCorEsts", ]
        tq = quantile(raws, Quant)
        tweedieEstsTrunc = tweedieEst;tweedieEstsTrunc[raws > tq] = raws[raws > tq]
        tweedieConvEstsTrunc = tweedieConvEsts;tweedieConvEstsTrunc[raws > tq] = raws[raws > tq]
        TanCorrectEst = TanCorrect(raws, bootEsts)$est
        TanCorrectEstParam = TanCorrect(raws, bootEstsParam)$est
        FayeCorrectEst = FayeCorrect(raws, bootEsts, MSEout)$est
        estForDensOracle = (if(RMSE) sqrt(zOracle) else zOracle)
        bootCorrectForde = forde(raws, rawEst[, "SE"]^2, allowInflate = TRUE)
        zwetRes = zwetWrapper(z = (raws - (mr <- mean(raws)))/rawEst[, "SE"], s = rawEst[, "SE"], int = 1)
        tweedieEstOracleDens = tweedieForm(estimates = rawEst[, "MSEhat"], analytical = FALSE, varsEstimates = rawEst[, "SE"]^2,
                                           breaks = breaksBoot, Plot = FALSE, topQuant = 1, estimatesForDens = estForDensOracle)["tweedieCorEsts", ]
        tweedieEstOracleVar = tweedieForm(estimates = rawEst[, "MSEhat"], analytical = FALSE,
                                          varsEstimates = varOracle, breaks = breaksBoot, Plot = FALSE,
                                          topQuant = 1)["tweedieCorEsts", ]
        tweedieEstOracleVarParam = tweedieForm(estimates = rawEst[, "MSEhat"], analytical = FALSE,
                                               varsEstimates = varOracle, breaks = breaksBoot, Plot = FALSE,
                                               topQuant = 1)["tweedieCorEsts", ]
        tweedieEstOracleDensVar = tweedieForm(estimates = rawEst[, "MSEhat"], analytical = FALSE,
                                              varsEstimates = varOracle, breaks = breaksBoot, Plot = FALSE, topQuant = 1,
                                              estimatesForDens = estForDensOracle)["tweedieCorEsts", ]
        ashObj = ash(raws, rawEst[, "SE"], method = "shrink", mode = "estimate")
        ashObjConv = getConvAsh(raws, sqrt(Vars), alpha1 = alpha1, ashObj = ashObj)
        out = list("resMat" = cbind("rawEst" = raws, "tweedieEst" = tweedieEst, "tweedieConvEsts" = tweedieConvEsts,
                                    "tweedieConvEstsTrunc" = tweedieConvEstsTrunc, "tweedieEstsTrunc" = tweedieEstsTrunc,
                     "TanCorrectEst" = TanCorrectEst, "TanCorrectEstParam" = TanCorrectEstParam, "FayeCorrectEst" = FayeCorrectEst,
                     "tweedieEstOracleDens" = tweedieEstOracleDens, "tweedieEstOracleVar" = tweedieEstOracleVar,
                     "bootCorrectForde" = bootCorrectForde, "VanZwet2021" = zwetRes$res$betaHats + mr,
                     "tweedieEstOracleVarParam" = tweedieEstOracleVarParam, "tweedieEstOracleDensVar" = tweedieEstOracleDensVar,
                     "ash" = ashObj$result$PosteriorMean, "ashConv" = ashObjConv$result$PosteriorMean[seq_len(p)]))
        return(out)
    })
}
```
