## Supplementary material for "The winner’s curse under dependence: repairing empirical Bayes using convoluted densities": Data, R-code and style file: getCVests.R

```
getCVests = function(trainDat, nOuterFolds, cvSplits, cvSplitsBoot, bootReps, nCores = 1){
    n = length(trainDat$y);
    seEsts = nestedCV(trainDat, nOuterFolds, cvSplits = cvSplits, nCores = nCores)
    bootEsts = bootCV(trainDat, n, bootReps, cvSplits = cvSplitsBoot,
                      mseOut = TRUE, nCores = nCores)
    bootEstsParam = bootCV(trainDat, n, bootReps, cvSplits = cvSplitsBoot,
                           paramBoot = TRUE, nCores = nCores)
    idIn = sample(n, n/2)
    trainDatIn = trainDatOut = trainDat
    trainDatIn$y = trainDatIn$y[idIn];trainDatIn$x = trainDatIn$x[idIn,, drop = FALSE]
    trainDatOut$y = trainDatOut$y[-idIn];trainDatOut$x = trainDatOut$x[-idIn,, drop = FALSE]
    estInSplit = simpleCV(trainDatIn, nOuterFolds);
    estOutSplit = nestedCV(trainDatOut, nOuterFolds, cvSplits = cvSplits, nCores = nCores)
    list("seEsts" = seEsts, "bootEsts" = bootEsts, "bootEstsParam" = bootEstsParam,
         "estInSplit" = estInSplit, "estOutSplit" = estOutSplit)
}
```
