## Supplementary material for "The winner’s curse under dependence: repairing empirical Bayes using convoluted densities": Data, R-code and style file: getDensEsts.R

```
#' Estimate the density and its derivative
#'
#' @param estimates matrix of estimates
#' @param ... passed on to the density estimation
getDensEsts = function(estimates, binned = TRUE,...){
    densEstObj = kde(estimates, binned = binned, ...)
    densDerivEstObj = kdde(estimates, binned = binned, deriv.order = 1, ...)
    return(list("densEst" = stats::predict(densEstObj, x = estimates ),
                "densEstDeriv" = stats::predict(densDerivEstObj, x = estimates )))
}
```
