## Supplementary material for "The winner’s curse under dependence: repairing empirical Bayes using convoluted densities": Data, R-code and style file: getMeanSigma.R

```
#Get mean and sigma for parametric bootstrap
getMeanSigma = function(dat){
    lapply(seq_len(ncol(dat$x)), function(j){
        Fit = lm.fit(x=cbind(1,dat$x[, j]), y=dat$y)
        list("Sigma" = sqrt(sum(Fit$residuals^2)/(length(dat$y)-2)),
             "Mean" = cbind(1,dat$x[, j]) %*% Fit$coef, "Coef" = Fit$coef)
    })
}
```
