## Supplementary material for "The winner’s curse under dependence: repairing empirical Bayes using convoluted densities": Data, R-code and style file: getSignifId.R

```
getSignifId = function(z, sigLevel){
    quants = pnorm(z)
    id <- quants>0.5
    quants[id] = pnorm(z[id], lower.tail = FALSE)
    pAdj = p.adjust(2*quants, method = "BH")
    which(pAdj<sigLevel)
}
```
