## Supplementary material for "The winner’s curse under dependence: repairing empirical Bayes using convoluted densities": Data, R-code and style file: getSimpleEstsMat.R

```
getSimpleEstsMat = function(mat, bootReps, bootRepsParam = bootReps, meanVec, VarsTrue, id = seq_len(nrow(mat)),
                            ciReturn = FALSE, varVar = TRUE,
                            extremes, quants = c(0.025, 0.975), parametric = TRUE, method = "eigen",
                            condml = TRUE, ciReturnBoot = TRUE, Ash = TRUE, numSamAsh = 10){
    names(quants) = quants;zQuants = qnorm(quants)
    p = ncol(mat);n = nrow(mat)
    ## Raw estimation ##
    rawEsts = colMeans(mat)
    #Data splitting
    idSplit = sample(nrow(mat), size = nrow(mat)/2)
    estInSplit = colMeans(matIn <- mat[idSplit,]);estOutSplit = colMeans(matOut <- mat[-idSplit,])
    # Bootstrap corrections
    bootEsts = bootNonParam(mat, bootReps, ciReturnBoot)
    bootEstsIn = t(sapply(bootEsts, function(x) x$out[,"estsIn"]))
    bootEstsOut = t(sapply(bootEsts, function(x) x$out[,"estsOut"]))
    bootCorrectFaye = FayeCorrect(rawEsts, bootEstsIn, bootEstsOut, ciReturn = ciReturnBoot, fullBootObj = bootEsts, zQuants = zQuants)
    bootCorrectTan = TanCorrect(rawEsts, bootEstsIn, ciReturn = ciReturnBoot, fullBootObj = bootEsts, quants = quants)
    bootCorrectForde = try(forde(rawEsts, Vars <- apply(mat, 2, var)/n))
    ses = sqrt(Vars)
    # Parametric bootstrap
    if(parametric){
        bootEstsParam = bootParam(mat, bootReps, rawEsts, ciReturnBoot, method = method)
        bootEstsInParam = t(sapply(bootEstsParam, function(x) x$out[,"estsIn"]))
        bootParamCorrectTan = TanCorrect(rawEsts, bootEstsInParam, ciReturn = ciReturnBoot,
                                         fullBootObj = bootEstsParam, quants = quants)
    } else {
        bootParamCorrectTan = NULL
    }
    #Empirical Bayes
    alpha1 = getAlpha1(mat) #unlikely to be zero due to non-negative definiteness
    #Naive Tweedie
    tweedieCorrectList = tweedieForm(rawEsts, varsEstimates = Vars, analytical = FALSE)
    #Convoluted tweedie
    tweedieCorrectConvList = tweedieForm(rawEsts, varsEstimates = Vars, alpha1 = alpha1, estimatesForDens = bootEstsIn)
    tweedieCorrectConvListSmooth = tweedieForm(rawEsts, varsEstimates = Vars, alpha1 = alpha1, smooth = TRUE, minAlpha1 = 0)
    #Ash
    if(Ash){
        ashObj = ash(rawEsts,ses, method = "shrink", mode = "estimate")
        #Convoluted density
        ashObjConv = getConvAsh(rawEsts,ses, alpha1, numSamAsh, ashObj)
    } else {ashObj = ashObjConv = NULL}
    #Van Zwet
    zwetRes = zwetWrapper(z = (rawEsts-(mr <- mean(rawEsts)))/ses, s =ses)
    zwetResConv = if(alpha1 < 0){
        zwetRes
    }else {
        zEsts = rnorm(p*10, rawEsts, ses*sqrt(alpha1))
        zwetWrapper(z = (rawEsts-mr)/ses, s = ses, zEsts = (zEsts-mr)/ses)
    }
    #Conditional likelihood
    condMLests = if(condml) corrCondML(raw = rawEsts, vars = Vars, dat = mat, extremes, ciReturn = ciReturn) else NULL
    if(!is.null(VarsTrue)){
        tweedieCorrectBootOracleVar = (tweedieCorrectBootOracleVarList <- tweedieForm(rawEsts, varsEstimates = VarsTrue, alpha1 = alpha1, trueVars = Vars))["tweedieCorEsts",]
        tweedieCorrectBootOracleDens = (tweedieCorrectBootOracleDensList <- tweedieForm(rawEsts, varsEstimates = Vars,
            alpha1 = 1, trueMeans = meanVec, trueVars = VarsTrue))["tweedieCorEsts",]
        tweedieCorrectBootOracleVarDens = (tweedieCorrectBootOracleVarDensList <-tweedieForm(rawEsts, varsEstimates = VarsTrue,
            alpha1 = 1, trueMeans = meanVec, trueVars = VarsTrue))["tweedieCorEsts",]
    } else{
        tweedieCorrectBootOracleVar = tweedieCorrectBootOracleDens = tweedieCorrectBootOracleVarDens = NULL
    }
    resMat = cbind("rawEst" = rawEsts, "FayeCorrectEst" = bootCorrectFaye$est,
                   "TanCorrectEst" = bootCorrectTan$est, "tweedieEst" = tweedieCorrectList["tweedieCorEsts",],
                   "VanZwet2021" = zwetRes$res$betaHats + mr,
                   "VanZwet2021conv" = zwetResConv$res$betaHats + mr,
                   "tweedieConvEsts" = tweedieCorrectConvList["tweedieCorEsts",],
                   "TanCorrectEstParam" = bootParamCorrectTan$est,
                   "bootCorrectForde" = bootCorrectForde,
                   "tweedieEstOracleVar" = tweedieCorrectBootOracleVar,
                   'tweedieEstOracleDens' = tweedieCorrectBootOracleDens,
                   'tweedieEstOracleDensVar' = tweedieCorrectBootOracleVarDens,
                   "meanVec" = meanVec, "VarsTrue" = VarsTrue, "Vars" = Vars,
                   "estInSplit" = estInSplit, "estOutSplit" = estOutSplit,
                   "ash" = ashObj$result$PosteriorMean,
                   "ashConv" = ashObjConv$result$PosteriorMean,
                   "tweedieConvEstsSmooth" = tweedieCorrectConvListSmooth["tweedieCorEsts",])
    ciList = if(ciReturn){
        sdOut = apply(mat[-idSplit,], 2, sd)/sqrt(nrow(mat)/2)
        ashCI = if(Ash) ashci(ashObj, level = nomCoverage <- 1-quants[1]*2, trace = FALSE)
        ashCIconv = if(Ash) ashci(ashObjConv, level = nomCoverage, trace = FALSE)
        rownames(ashCI) = rownames(ashCIconv) = names(rawEsts)
        CIestOutSplit = estOutSplit + outer(sdOut, zQuants)#Use z quantiles here for comparability
        list("rawEst" = rawEsts + outer(sqrt(Vars), zQuants),
             "tweedieEst" = buildCredInt(tweedieCorrectList, n = n, Vars, zQuants, varVar = varVar),
             "tweedieConvEsts" = buildCredInt(tweedieCorrectConvList, n = n, Vars, zQuants, bootEsts = bootEstsIn, varVar = varVar),
             "tweedieEstOracleVar" = buildCredInt(tweedieCorrectBootOracleVarList, n = n, VarsTrue, zQuants, varVar = varVar, bootEsts = bootEstsIn, trueVars = Vars),
             "tweedieEstOracleDens" = buildCredInt(tweedieCorrectBootOracleDensList, n = n, varVar = varVar, Vars, zQuants, addVarDens = FALSE, trueVars = VarsTrue),
             "tweedieEstOracleDensVar" = buildCredInt(tweedieCorrectBootOracleVarDensList, varVar = varVar, n = n, VarsTrue, zQuants, addVarDens = FALSE),
             "estOutSplit" = CIestOutSplit, "FayeCorrectEst" = bootCorrectFaye$ci,
             "VanZwet2021" = zwetRes$confInts + mr,
             "VanZwet2021conv" = zwetResConv$confInts + mr,
             "TanCorrectEst" = bootCorrectTan$ci, "TanCorrectEstParam" = bootParamCorrectTan$ci,
             "ash" = ashCI, "ashConv" = ashCIconv)
    }
    ## Test statistics estimation ##
    rawEstsZ = rawEsts/sqrt(Vars)
    bootEstsInZ = t(sapply(bootEsts, function(x) x$out[,"estsIn"]/sqrt(x$out[,"varEstsIn"])))
    bootEstsOutZ = t(sapply(bootEsts, function(x) x$out[,"estsOut"]/sqrt(x$out[,"varEstsOut"])))
    bootCorrectFayeZ = FayeCorrect(rawEstsZ, bootEstsInZ, bootEstsOutZ, ciReturn = ciReturnBoot,
                                   fullBootObj = bootEsts, zQuants = zQuants, testStatistic = TRUE)
    bootCorrectTanZ = TanCorrect(rawEstsZ, bootEstsInZ, ciReturn = ciReturnBoot,
                                fullBootObj = bootEsts, quants = quants, testStatistic = TRUE)
    tweedieCorrectListZ = tweedieForm(rawEstsZ, varsEstimates = 1, analytical = FALSE)
    tweedieCorrectConvListZ = tweedieForm(rawEstsZ, varsEstimates = 1, alpha1 = alpha1)
    #Conditional likelihood
    bootCorrectFordeZ = forde(rawEstsZ, 1)
    condMLestsZ = if(condml) corrCondML(raw = rawEstsZ, testStatistic = TRUE, extremes = extremes, ciReturn = ciReturn) else NULL
    #Parametric bootstrap
    if(parametric){
        bootEstsInParamZ = t(sapply(bootEstsParam, function(x) x$out[,"estsIn"]/sqrt(x$out[,"varEstsIn"])))
        bootParamCorrectTanZ = TanCorrect(rawEstsZ, bootEstsInParamZ, ciReturn = ciReturnBoot,
                                         fullBootObj = bootEstsParam, quants = quants, testStatistic = TRUE)
    } else {
        bootEstsParamCorZ = bootParamCorrectTanZ = NULL
    }
    #Data splitting
    estInSplitZ = estInSplit/sqrt(rowMeans((t(matIn) - estInSplit)^2)/(nrow(matIn) - 1))
    estOutSplitZ = estOutSplit/sqrt(rowMeans((t(matOut) - estOutSplit)^2)/(nrow(matOut) - 1))
    #Ash
    if(Ash){
        #Keep mode at estimate for comparability
        ashObjZ = ash(rawEstsZ, rep(1, p), method = "shrink", mode = "estimate")
        ashObjConvZ = getConvAsh(rawEstsZ, 1, alpha1, numSamAsh, ashObjZ)
    } else {ashObjZ = ashObjConvZ = NULL}
    #Cat scores
    catScores = shrinkcat.stat(mat, L = rep(0, n), verbose = FALSE)
    resMatZ = cbind("rawEst" = rawEstsZ, "FayeCorrectEst" = bootCorrectFayeZ$est,
                   "TanCorrectEst" = bootCorrectTanZ$est, "tweedieEst" = tweedieCorrectListZ["tweedieCorEsts", ],
                   "tweedieConvEsts" = tweedieCorrectConvListZ["tweedieCorEsts",],
                   "TanCorrectEstParam" = bootParamCorrectTanZ$est,
                   "VanZwet2021" = zwetRes$res$betaHats/zwetRes$res$s,
                   "VanZwet2021conv" = zwetResConv$res$betaHats/zwetResConv$res$s,
                   "bootCorrectForde" = bootCorrectFordeZ*sqrt(Vars),
                   "estInSplit" = estInSplitZ, "estOutSplit" = estOutSplitZ,
                   "ash" = ashObjZ$result$PosteriorMean, "ashConv" = ashObjConvZ$result$PosteriorMean,
                   "catScores" = catScores)
    ciListZ = if(ciReturn){
        ashCI = if(Ash) ashci(ashObjZ, level = nomCoverage, trace = FALSE)
        ashCIconvZ = if(Ash) ashci(ashObjConvZ, level = nomCoverage,
                                  trace = FALSE)
        rownames(ashCI) = rownames(ashCIconv) = names(rawEstsZ)
        CIestOutSplitZ = outer(estOutSplitZ, zQuants, FUN = "+")#Use z quantiles here for comparability
        list("rawEst" = outer(rawEstsZ, zQuants, FUN = "+"),
             "tweedieEst" = buildCredInt(tweedieCorrectListZ, n = n, 1, zQuants, varVar = varVar),
             "tweedieConvEsts" = buildCredInt(tweedieCorrectConvListZ, n = n, 1, zQuants, varVar = varVar),
             "estOutSplit" = CIestOutSplitZ, "FayeCorrectEst" = bootCorrectFayeZ$ci,
             "TanCorrectEst" = bootCorrectTanZ$ci, "TanCorrectEstParam" = bootParamCorrectTanZ$ci,
             "ash" = ashCI, "ashConv" = ashCIconvZ)
    }
    idSignif = getSignifId(resMatZ[, "rawEst"], sigLevel = sigLevel)
    condMLestsFdr = if(condml) corrCondML(raw = rawEstsZ, testStatistic = TRUE, dat = mat,
                                          extremes = sum(resMatZ[idSignif, "rawEst"]<0), ciReturn = ciReturn) else NULL
    list("condMLests" = condMLests, "resMat" = resMat, "ciList" = ciList, "condMLestsFdr" = condMLestsFdr,
         "condMLestsZ" = condMLestsZ, "resMatZ" = resMatZ, "ciListZ" = ciListZ, "mat" = mat)
}
```
