## Supplementary material for "The winner’s curse under dependence: repairing empirical Bayes using convoluted densities": Data, R-code and style file: getTestMSE.R

```
#Get test MSE given training and test data
getTestMSE = function(trainDat, testDat, highDim = FALSE,...){
    if(highDim){
        multiMod = cv.glmnet(x = trainDat$x, y = trainDat$y, ...)
        mean((predict(multiMod, testDat$x)-testDat$y)^2)
    } else {
        vapply(seq_len(NCOL(trainDat$x)), FUN.VALUE = double(1), function(j){
            predTest = predLin(trainDat$x[, j], trainDat$y, testDat$x[, j])
            mean((predTest-testDat$y)^2)
        })
    }
}
#Generate data and get test MSE
genDataAndgetTestMSE = function(n, p, betas = integer(p), testDat, ...){
    trainDat = genDat(n, p, betas = betas, ...)
    if(missing(testDat))
        testDat = genDat(n, p, betas = betas, ...)
    MSE = getTestMSE(trainDat, testDat, ...)
    list("MSE" = MSE, "margVar" = var(testDat$y))
}
```
