## Supplementary material for "The winner’s curse under dependence: repairing empirical Bayes using convoluted densities": Data, R-code and style file: invSqrt.R

```
invSqrt = function (A)
{
    ei <- eigen(A, symmetric = TRUE)
    d <- pmax(ei$values, 10^-12)
    d2 <- 1/sqrt(d)
    d2[d == 0] <- 0
    ei$vectors %*% (if (length(d2) == 1)
        d2
        else diag(d2)) %*% t(ei$vectors)
}
```
