## Supplementary material for "The winner’s curse under dependence: repairing empirical Bayes using convoluted densities": Data, R-code and style file: LindseysMethod.R

```
#Lindseys method in 1D
LindseysMethod = function(z, zObs = z, breaks = 8e1, degree = 7, maxIter = 1e2,
                          Plot = FALSE, topQuant = 1, returnFull = FALSE, returnFit = FALSE,...){
    iter = 1
    while((iter==1 || inherits(fitpois, "try-error")) && iter <=maxIter){
        breaks = breaks+1; iter = iter+1
        histObj <- hist(z, breaks = breaks, plot = FALSE)
        # getting the middle of the bin, for later
        #The 2.5% features with highest MSE are ignored, densities are set to 0
        largeZ = quantile(z, probs = topQuant)
        midPoints <- histObj$mids; Counts = histObj$counts
        idZ = z > largeZ
        # Defining the degree of the polynomial expansion
        # Easy function to do that is with polym
        ply_expansion <- polym(midPoints, degree = degree, raw = TRUE)
        ply_expansionObs <- polym(zObs, degree = degree, raw = TRUE)
        # Fit the poisson regression
        offS = rep(log(length(z)*diff(range(z))/breaks), length(Counts))
        fitpois <- try(glm(Counts ~ ply_expansion , family = poisson(), offset = offS), silent = TRUE)
    }
    if(returnFit){
        return(coef(fitpois))
    }
    manualLogDeriv = evalFitLogDeriv(coef(fitpois), expansion = ply_expansionObs)
    manualLogDeriv[zObs > largeZ] = 0
    manualLogDens = cbind(1,ply_expansionObs) %*% coef(fitpois)
    #Second order derivative
    manualLogDeriv2 = c(cbind(1, ply_expansionObs[, -(degree+c(0,-1))]) %*% (coef(fitpois)[-c(1,2)]*seq_len(degree)[-1]*seq_len(degree-1)))
    Order = order(zObs)
    if(Plot){
        parTmp = par(no.readonly = TRUE)
        par(mfrow = c(1,3))
        plot(zObs[Order], manualLogDens[Order], xlab = "Statistic", ylab = "Log density", type = "l")
        plot(zObs[Order], manualLogDeriv[Order], xlab = "Statistic", ylab = "Derivative of log density", type = "l")
        plot(zObs[Order], manualLogDeriv2[Order], xlab = "Statistic", ylab = "Second order derivative of log density", type = "l")
        abline(h = 0, lty = "dotted")
        par(parTmp)
    }
    if(returnFull){
        return(cbind("zObs" = zObs[Order], "logDens" = manualLogDens[Order],
                     "logDensDeriv" = manualLogDeriv[Order], "manualLogDeriv2" = manualLogDeriv2[Order]))
    } else {
        return(rbind("deriv1" = manualLogDeriv, "deriv2" = manualLogDeriv2))
    }
}
```
