## Supplementary material for "The winner’s curse under dependence: repairing empirical Bayes using convoluted densities": Data, R-code and style file: makeBlockFolds.R

```
makeFolds = function(nfolds, data, cvType){
    folds = sample(nfolds, nrow(data), replace = TRUE)
    while(any(table(folds)<=1)){
        folds = makeFolds(nfolds, data, cvType)
    }
    return(folds)
}
```
