## Supplementary material for "The winner’s curse under dependence: repairing empirical Bayes using convoluted densities": Data, R-code and style file: makeMean.R

```
#Construct a mean vector
makeMean = function(p, sd, effSize, effFrac, mode){
    if(mode=="dense"){
        rnorm(p, sd = sdMean)
    } else if(mode=="sparse"){
        c(integer(p*(1-effFrac)), rnorm(p*effFrac, effSize, sd = sd))
    }
}
```
