## Supplementary material for "The winner’s curse under dependence: repairing empirical Bayes using convoluted densities": Data, R-code and style file: nestedCV.R

```
#' Nested CV per feature
nestedCV = function(dat, nOuterFolds, cvSplits, nInnerFolds = nOuterFolds - 1, nCores = 1,loc = NULL, verbose = FALSE, saveFolder = NULL,
                    toBeSaved = FALSE, mc.preschedule = FALSE,...){
    out = lapply(seq_len(ncol(dat$x)), function(j){
        if(verbose && (j %% 50)==0) cat("Feature", j, "\t")
        if(!toBeSaved ||  (!is.null(saveFolder) && !file.exists(saveFile <- paste0(saveFolder, "/cvSplitReps", j, ".RData")))){
            #Repeat splitting into folds
            cvSplitReps = mclapply(mc.cores = nCores, seq_len(cvSplits), function(cvs){
                folds = sample(rep(unFolds <- seq_len(nOuterFolds), length.out = nrow(dat$x)))
                lapply(unFolds, function(uf){
                    idTrain = folds!=uf
                    yTrain = dat$y[idTrain]; xTrain <- dat$x[idTrain,j];locTrain = loc[idTrain,]
                    predTest = predLin(xTrain, yTrain, dat$x[!idTrain, j], loc = locTrain, ...)
                    eOut = (predTest-dat$y[!idTrain])^2
                    eBarOut = mean(eOut)
                    b = var(eOut)/length(eOut)
                    inFolds = sample(rep(unFoldsIn <- seq_len(nInnerFolds), length.out = sum(idTrain)))
                    errHatTilde = mean(vapply(FUN.VALUE = double(1), unFoldsIn, function(inf){
                        idTrainIn = inFolds!=inf
                        predTestIn = predLin(xTrain[idTrainIn], yTrain[idTrainIn], xTrain[!idTrainIn], loc = locTrain[idTrainIn,],...)
                        mean((predTestIn-yTrain[!idTrainIn])^2)
                    }))
                    a = (errHatTilde-eBarOut)^2
                    list("a" = a, "b" = b, "errHatTilde" = errHatTilde, "eOut" = eOut, "margVar" = var(dat$y[!idTrain]))
                })
            })
            seNested = getSEsNested(cvSplitReps, nOuterFolds = nOuterFolds, n = nrow(dat$x))
            if(toBeSaved){save(seNested, file = saveFile)}
        } else if(toBeSaved){
            load(saveFile)
        }
        seNested
    })
    names(out) = colnames(dat$x)
    return(out)
}
#Build confidence intervals
getSEsNested = function(cvSplitReps, nOuterFolds, n){
    ErrNCV = mean(unlist(sapply(cvSplitReps, function(y) sapply(y, function(x) if(is.list(x)) x[["errHatTilde"]] else NA))), na.rm = TRUE)
    MSEhat = mean(unlist(sapply(cvSplitReps, function(y) sapply(y, function(x) if(is.list(x)) x[["a"]] else NA))), na.rm = TRUE) -
        mean(unlist(sapply(cvSplitReps, function(y) sapply(y, function(x) if(is.list(x)) x[["b"]] else NA))), na.rm = TRUE)
    errOuter0 = lapply(cvSplitReps, function(y) lapply(y, function(x) if(is.list(x)) x[["eOut"]] else NA))
    mseOuter = sapply(errOuter0, function(w) sapply(w, mean, na.rm = TRUE))
    errOuter = unlist(errOuter0)
    margVars = sapply(cvSplitReps, function(y) sapply(y, function(x) if(is.list(x)) x[["margVar"]] else NA))
    SEest = sqrt(max(0, nOuterFolds/(nOuterFolds-1)*MSEhat))
    naiveRMSE = sd(errOuter, na.rm = TRUE)/sqrt(n)
    maxMSE = naiveRMSE * sqrt(nOuterFolds)
    if(is.na(SEest) || (SEest < naiveRMSE)){ #See below equation (17), prevent implausible values
        SEest = naiveRMSE
    } else if (SEest > maxMSE){
        SEest = maxMSE
    }
    #Correct the bias
    ErrCV = mean(errOuter, na.rm = TRUE)
    Bias = (1+(nOuterFolds-2)/nOuterFolds)*(ErrNCV-ErrCV)
    ErrNCVBC = ErrNCV - Bias#Bias correction
    cbind("MSEhat" = c("Naive" = ErrCV, "Bates" = ErrNCVBC),
          "SE" = c("Naive" = naiveRMSE, "Bates" = SEest))
}
nestedCVglmnet = function(dat, nOuterFolds, cvSplits, nInnerFolds = nOuterFolds - 1,
                          nCores = 1, alpha, loc = NULL, filterFun = NULL, saveFolder = NULL,...){
    #Repeat splitting into folds
    cvSplitReps = lapply(seq_len(cvSplits), function(cvs){
        saveFile = paste0(saveFolder, "/rep", cvs, ".RData")
        if(!file.exists(saveFile)){
            cat("cvSplit", cvs, "\t")
            folds = sample(rep(unFolds <- seq_len(nOuterFolds), length.out = nrow(dat$x)))
            tmp = mclapply(mc.cores = nCores, unFolds, function(uf){
                idTrain = folds!=uf
                yTrain = dat$y[idTrain]; xTrain <- dat$x[idTrain,]
                if(!is.null(filterFun)){
                    xTrain = filterFun(xTrain, yTrain)
                }
                predTest = predGlmnet(xTrain, yTrain, dat$x[!idTrain, colnames(xTrain)], nfolds = max(4,nInnerFolds-1),
                                      alpha = alpha, loc = loc[idTrain, ], ...)
                eOut = (predTest-dat$y[!idTrain])^2
                eBarOut = mean(eOut)
                b = var(eOut)/length(eOut) #Also like this in nestedcv package
                inFolds = sample(rep(unFoldsIn <- seq_len(nInnerFolds), length.out = sum(idTrain)))
                cat("Starting nested CV\t")
                errHatTilde = mean(unlist(sapply(unFoldsIn, function(inf){
                    idTrainIn = inFolds!=inf
                    xIn = xTrain[idTrainIn,]; yIn = yTrain[idTrainIn]
                        if(!is.null(filterFun)){
                            xIn = filterFun(xIn, yIn)
                            if(!ncol(xIn)){
                                return((yTrain[!idTrainIn]-mean(yTrain[!idTrainIn]))^2)
                            }
                        }
                    predTestIn = predGlmnet(xIn, yIn, xTrain[!idTrainIn,colnames(xIn)],
                                         nfolds = max(4,nInnerFolds-1), alpha = alpha, loc = loc[idTrain,][idTrainIn, ], ...)
                    (c(predTestIn)-yTrain[!idTrainIn])^2
                })))
                a = (errHatTilde-eBarOut)^2
                list("a" = a, "b" = b, "errHatTilde" = errHatTilde, "eOut" = eOut, "margVar" = var(dat$y[!idTrain]))
            })
            save(tmp, file = saveFile)
        } else load(saveFile)
        tmp
    })
    getSEsNested(cvSplitReps, nOuterFolds = nOuterFolds, n = nrow(dat$x))
}
```
