## Supplementary material for "The winner’s curse under dependence: repairing empirical Bayes using convoluted densities": Data, R-code and style file: nlpdenWrapper.R

```
#' A wrapper for the nlpden function, exploiting the Gaussian additive noise nature of the data for improved density estimation
nlpdenWrapper = function(z, sdVec, zObs = z, Plot = FALSE, returnFull = FALSE, breaks = NULL, ...){
    nlpEst = nlpden(z/sdVec, ...) #Standardize before estimating density
    if(Plot){
        plot(nlpEst, type ="l")
    }
    nlpEst$logDens = log(nlpEst$density_estimate)
    f = approxfun(nlpEst$x_axis, nlpEst$logDens)
    logDensDeriv = grad(f, zObsStand <- zObs/sdVec)
    if(returnFull){
        Ord = order(zObsStand)
        return(cbind("zObs" = zObsStand[Ord], "logDens" = approx(nlpEst$x_axis, nlpEst$logDens, zObsStand[Ord])$y,
                     "logDensDeriv" = logDensDeriv[Ord]))
    } else {
        return(rbind("deriv1" = logDensDeriv, "deriv2" = NA))
    }
}
```
