## Supplementary material for "The winner’s curse under dependence: repairing empirical Bayes using convoluted densities": Data, R-code and style file: normalize.R

```
#Normalization wrapper
normalize = function(mat, trans, design = formula("~1"), df = DataFrame("row.names" = rownames(mat))){
    mat = apply(mat, c(1,2), as.integer)
    out = if(trans=="Rel"){
        mat/rowSums(mat)
        } else if(trans %in% c("Vst", "Rlog")){
            require(DESeq2)
            object <- DESeqDataSetFromMatrix(t(mat), df, design)
            object = estimateSizeFactors(object, type = if(all(colSums(mat==0)>0)) "poscounts" else "ratio")
            object = if(trans=="Vst"){
                varianceStabilizingTransformation(object)
            } else if(trans=="Rlog"){
                rlog(object)
            }
            t(assay(object))
        } else if(trans=="Clr"){
            clrTransformMat(mat)
        }else {
           mat
            }
    dimnames(out) = dimnames(mat)
    return(out)
}
```
