## Supplementary material for "The winner’s curse under dependence: repairing empirical Bayes using convoluted densities": Data, R-code and style file: plotBoxRMSE.R

```
plotBoxRMSE = function(resList, trueMSE, Incl, returnDf = FALSE){
    rmseMat = t(sapply(which(sapply(resList, is.list)), function(i){
        colMeans((resList[[i]]$resMat-sqrt(if(is.matrix(trueMSE)) trueMSE[, i] else trueMSE))^2)
    }))
    plotExclude = c('tweedieEst', 'tweedieMSEBoot632','tweedieMSEdoubleBoot632', "tweedieBootEstsNlpden",
                    "tweedieBootParamEstsNlpden" , 'tweedieMSEdoubleBootOOB', 'tweedieMSEBootOOB', "estInSplit" , "catScores",
                    "rawBoot632", "rawBootOOB", "TanCorrectEstParam", grep("Stand", colnames(rmseMat), value = TRUE),
                    "rawsCatBoot", "rawsCatBootParam")
    rmseMat = rmseMat[, colnames(rmseMat) %in% methodLevels[methodLabels %in% Incl & !(methodLevels %in% plotExclude)]]
    moltRmse = melt(rmseMat, value.name = "RMSE", varnames = c("Rep", "Method"))
    moltRmse$Method = factor(moltRmse$Method, levels = methodLevels, labels = methodLabels)
    meansDf = aggregate(RMSE ~ Method , FUN = mean, data =  moltRmse)
    if(returnDf){
        return(moltRmse)
    }
    ggplot(data = moltRmse, aes_string(y = "RMSE", x = "Method", colour = "Method")) + geom_boxplot() +
        theme(axis.text.x = element_blank(), axis.ticks.x = element_blank()) +
        xlab("") + ylab("MSE of all estimates") + scale_y_log10() +
        geom_point(data = meansDf, shape = 5) +
        scale_colour_manual(values = colourVec[names(colourVec) %in% unique(moltRmse$Method)])
}
```
