## Supplementary material for "The winner’s curse under dependence: repairing empirical Bayes using convoluted densities": Data, R-code and style file: plotCoverage.R

```
plotCoverage = function(resList, extremes, Palette = "Paired", quantity = "FCR", Mode  ="dense",
                        Methods2Plot = c(plotNormal, "estOutSplit"), convertMethod = FALSE,
                        lineSize = 0.55, legSize = 8, returnDf = FALSE, extPlot = "small", methodLevelsIn = methodLevels, methodLabelsIn = methodLabels){
    coverageList = lapply(modes, function(mode){
        lapply(Sds, function(Sd){
            Sd = as.character(Sd)
            topMats = lapply(resList[[mode]][[Sd]], function(x) {
                out = lapply(nm <- names(x$ciList)[!sapply(x$ciList, is.null)], function(m){
                    Ord = order(x$resMat[, switch(m, "estOutSplit" = "estInSplit", m)])
                    Cov = x$ciList[[m]][,1] < x$resMat[, "meanVec"] & x$ciList[[m]][,2] > x$resMat[, "meanVec"]
                    switch(quantity, "FCR" = cbind("small" = cumsum(Cov[Ord][extremes]),
                                               "large" = cumsum(Cov[Ord][length(Cov)-extremes+1]))/seq_along(extremes),
                                 "RCC" = cbind("small" = Cov[Ord][extremes],
                                                  "large" = Cov[Ord][length(Cov)-extremes+1]))
                })
                names(out) = nm
                out
            })
            lapply(extremes, function(ext){
                extChar = as.character(ext)
                condMat = vapply(seq_along(resList[[mode]][[Sd]]), FUN.VALUE = matrix(0, ext, 2), function(i){
                    x = resList[[mode]][[Sd]][[i]]
                    cbind("small" = x$condMLests[[extChar]]$small[, 2] < x$resMat[as.integer(rownames(x$condMLests[[extChar]]$small)), "meanVec"] &
                                            x$condMLests[[extChar]]$small[, 3] > x$resMat[as.integer(rownames(x$condMLests[[extChar]]$small)), "meanVec"],
                                 "large" = x$condMLests[[extChar]]$large[, 2] < x$resMat[as.integer(rownames(x$condMLests[[extChar]]$large)), "meanVec"] &
                                     x$condMLests[[extChar]]$large[, 3] > x$resMat[as.integer(rownames(x$condMLests[[extChar]]$large)), "meanVec"])
                    })
                condOut = rowMeans(switch(quantity,
                       "FCR" = array(apply(condMat, c(2,3), cumsum)/seq_len(ext), dim = c(ext, 2, dim(condMat)[3])),
                       "RCC" = condMat), dims = 2)
                if(any(grepl(pattern = "rescale", names(resList[[1]][[1]])))){
                condMatRescale = vapply(seq_along(resList[[mode]][[Sd]]), FUN.VALUE = matrix(0, ext, 2), function(i){
                    x = resList[[mode]][[Sd]][[i]];trueZ = x$resMat[, "meanVec"]/sqrt(x$resMat[, "VarsTrue"])
                    cbind("small" = x$condML_rescale[[extChar]]$small[, 2] < x$resMat[as.integer(rownames(x$condML_rescale[[extChar]]$small)), "meanVec"] &
                              x$condML_rescale[[extChar]]$small[, 3] > x$resMat[as.integer(rownames(x$condML_rescale[[extChar]]$small)), "meanVec"],
                          "large" = x$condML_rescale[[extChar]]$large[, 2] < x$resMat[as.integer(rownames(x$condML_rescale[[extChar]]$small)), "meanVec"] &
                              x$condML_rescale[[extChar]]$large[, 3] > x$resMat[as.integer(rownames(x$condML_rescale[[extChar]]$small)), "meanVec"])
                })
                condOutRescale = rowMeans(switch(quantity,
                                                 "FCR" = array(apply(condMatRescale, c(2,3), cumsum)/seq_len(ext), dim = c(ext, 2, dim(condMatRescale)[3])),
                                                 "RCC" = condMatRescale), dims = 2)
                } else {condOutRescale = NULL}
                rbind(t(rowMeans(dims = 2, vapply(topMats, FUN.VALUE = matrix(0, 2, length(topMats[[1]])), function(x) sapply(x, function(yy) {
                    c("small" = yy[ext, "small"], "large" = yy[ext, "large"])
                })))), "condML" = condOut[ext, ], "condML_rescale" = condOutRescale[ext,])
            })
        })
    })
    moltErrorList = melt(coverageList, value.name = "Coverage")
    coverageListZ = lapply(modes, function(mode){
        lapply(Sds, function(Sd){
            Sd = as.character(Sd)
        topMats = lapply(resList[[mode]][[Sd]], function(x) {
            trueZ = x$resMat[, "meanVec"]/sqrt(x$resMat[, "VarsTrue"])
            out = lapply(nm <- names(x$ciListZ)[!sapply(x$ciListZ, is.null)], function(m){
                Ord = order(x$resMatZ[, switch(m, "estOutSplit" = "estInSplit", m)])
               Cov = x$ciListZ[[m]][,1] < trueZ & x$ciListZ[[m]][,2] > trueZ
                switch(quantity, "FCR" = cbind("small" = cumsum(Cov[Ord][extremes]),
                                           "large" = cumsum(Cov[Ord][length(Cov)-extremes+1]))/seq_along(extremes),
                       "RCC" = cbind("small" = Cov[Ord][extremes],
                                     "large" = Cov[Ord][length(Cov)-extremes+1]))
            })
            names(out) = nm
            out
        })
        lapply(extremes, function(ext){
            extChar = as.character(ext)
            condMat = vapply(seq_along(resList[[mode]][[Sd]]), FUN.VALUE = matrix(0, ext, 2), function(i){
                x = resList[[mode]][[Sd]][[i]];trueZ = x$resMat[, "meanVec"]/sqrt(x$resMat[, "VarsTrue"])
                cbind("small" = x$condMLestsZ[[extChar]]$small[, 2] < trueZ[as.integer(rownames(x$condMLestsZ[[extChar]]$small))] &
                          x$condMLestsZ[[extChar]]$small[, 3] > trueZ[as.integer(rownames(x$condMLestsZ[[extChar]]$small))],
                      "large" = x$condMLestsZ[[extChar]]$large[, 2] < trueZ[as.integer(rownames(x$condMLestsZ[[extChar]]$large))] &
                          x$condMLestsZ[[extChar]]$large[, 3] > trueZ[as.integer(rownames(x$condMLestsZ[[extChar]]$large))])
            })
            condOut = rowMeans(switch(quantity,
                                      "FCR" = array(apply(condMat, c(2,3), cumsum)/seq_len(ext), dim = c(ext, 2, dim(condMat)[3])),
                                      "RCC" = condMat), dims = 2)
            rbind(t(rowMeans(dims = 2, vapply(topMats, FUN.VALUE = matrix(0, 2, length(topMats[[1]])), function(x) sapply(x, function(yy) {
                c("small" = yy[ext, "small"], "large" = yy[ext, "large"])
            })))), "condML" = condOut[ext, ])
        })
    })
    })
    moltErrorListZ = melt(coverageListZ, value.name = "Coverage")
    names(moltErrorList)[c(1:2, 4:6)] = names(moltErrorListZ)[c(1:2, 4:6)] = c("Method", "extreme","Top_features", "Cor", "mode")
    moltErrorList$what = "Raw estimate";moltErrorListZ$what = "Test statistic"
    moltErrorListFull = rbind(moltErrorList, moltErrorListZ)
    moltErrorListFull = moltErrorListFull[moltErrorListFull$Method %in% Methods2Plot,]
    moltErrorListFull$Cor = as.numeric(moltErrorListFull$Cor)
    if(convertMethod)
        moltErrorListFull$Method = factor(moltErrorListFull$Method, ordered = TRUE, levels = methodLevelsIn, labels = methodLabelsIn)
    moltErrorListFull = droplevels(moltErrorListFull)
    moltErrorListFull$Top_features = as.integer(moltErrorListFull$Top_features)
    if(returnDf){
        moltErrorListFull$Performance_measure = "Coverage"
        moltErrorListFull$value = moltErrorListFull$Coverage
        moltErrorListFull$Coverage = NULL
        return(moltErrorListFull[moltErrorListFull$extreme == extPlot, ])
    } else {
    ggplot(data = moltErrorListFull[with(moltErrorListFull, extreme == extPlot & what == "Raw estimate" & mode == Mode), ],
           aes_string(y = "Coverage", colour = "Method", x = "Top_features")) +
            geom_line(size = lineSize) + facet_grid( ~ Cor, scales = "free_y") +
        xlab("Number of extreme estimates") +
        geom_hline(yintercept = 1-sigLevel, linetype = "dotted") +
        ylab(switch(quantity, "FCR" = "Coverage", "RCC" = "Rank conditional coverage")) +
        scale_colour_manual(values = colourVec[names(colourVec) %in% unique(moltErrorListFull$Method)])
    }
}
plotCoverageFDR = function(resList, Palette = "Paired", quantity = "FCR", Mode  ="dense",
                        Methods2Plot = c(plotNormal, "estOutSplit"), convertMethod = FALSE,
                        lineSize = 0.55, legSize = 8, returnDf = FALSE, extPlot = "small", methodLevelsIn = methodLevels, methodLabelsIn = methodLabels){
    coverageList = lapply(modes, function(mode){
        lapply(Sds, function(Sd){
            Sd = as.character(Sd)
            topMats = lapply(resList[[mode]][[Sd]], function(x) {
                extremes = getExtremes(x)
                out = lapply(nm <- names(x$ciList)[!sapply(x$ciList, is.null)], function(m){
                    Ord = order(x$resMat[, switch(m, "estOutSplit" = "estInSplit", m)])
                    Cov = x$ciList[[m]][,1] < x$resMat[, "meanVec"] & x$ciList[[m]][,2] > x$resMat[, "meanVec"]
                    switch(quantity, "FCR" = cbind("small" = cumsum(Cov[Ord][extremes]),
                                                   "large" = cumsum(Cov[Ord][length(Cov)-extremes+1]))/seq_along(extremes),
                           "RCC" = cbind("small" = Cov[Ord][extremes],
                                         "large" = Cov[Ord][length(Cov)-extremes+1]))
                })
                names(out) = nm
                out
            })
            condMat = vapply(seq_along(resList[[mode]][[Sd]]), FUN.VALUE = matrix(0, 1, 2), function(i){
                    extremes = NROW(topMats[[i]][[1]]);names(extremes) = extremes;extChar = as.character(extremes)
                    if(NCOL(topMats[[i]][[1]])==0){
                        return(NULL)
                    }
                    x = resList[[mode]][[Sd]][[i]]
                    cbind("small" = x$condMLests[[extChar]]$small[, 2] < x$resMat[as.integer(rownames(x$condMLests[[extChar]]$small)), "meanVec"] &
                              x$condMLests[[extChar]]$small[, 3] > x$resMat[as.integer(rownames(x$condMLests[[extChar]]$small)), "meanVec"],
                          "large" = x$condMLests[[extChar]]$large[, 2] < x$resMat[as.integer(rownames(x$condMLests[[extChar]]$large)), "meanVec"] &
                              x$condMLests[[extChar]]$large[, 3] > x$resMat[as.integer(rownames(x$condMLests[[extChar]]$large)), "meanVec"])
                })
                condOut = rowMeans(switch(quantity,
                                          "FCR" = array(apply(condMat, c(2,3), cumsum), dim = c(1, 2, dim(condMat)[3])),
                                          "RCC" = condMat), dims = 2)
                rbind(t(rowMeans(na.rm = TRUE, dims = 2, vapply(topMats, FUN.VALUE = matrix(0, 2, length(topMats[[1]])), function(x) sapply(x, function(yy) {
                    if(!NROW(yy) || NCOL(yy) <=1){
                        return(c("small" = NA, "large" = NA))
                    } else {
                        c("small" = yy[1, "small"], "large" = yy[1, "large"])
                    }
                })))), "condML" = condOut[1, ])
            })
    })
    moltErrorList = melt(coverageList, value.name = "value")
    names(moltErrorList)[c(1:2, 4:5)] = c("Method", "extreme", "Cor", "mode")
    moltErrorList$what = "Raw estimate"
    moltErrorList = moltErrorList[moltErrorList$Method %in% Methods2Plot,]
    moltErrorList$Cor = as.numeric(moltErrorList$Cor)
    if(convertMethod)
        moltErrorList$Method = factor(moltErrorList$Method, ordered = TRUE, levels = methodLevelsIn, labels = methodLabelsIn)
    moltErrorList = droplevels(moltErrorList)
    moltErrorList$Performance_measure = "Coverage"
    moltErrorList[moltErrorList$extreme == extPlot, ]
}
```
