## Supplementary material for "The winner’s curse under dependence: repairing empirical Bayes using convoluted densities": Data, R-code and style file: plotExtremes.R

```
plotExtremes = function(estsMat, meanVec, extremes = seq_len(30), includeOut = TRUE, measures = c("coverage", "width"),
                        Excl = c("estInSplit", methodLevels[!(methodLabels %in% Incl[Incl!="Separate estimation"])]),
                        Incl = namesCV, power = 1:2, boxPlot = FALSE, lineSize = 0.4, returnDf = FALSE){
    names(extremes) = extremes; names(measures) = measures
    p = length(meanVec)
    powList = lapply(power, function(pow){
        errorList = lapply(seq_along(estsMat), function(i){
            x = estsMat[[i]]$resMat
            meanVec = if(is.vector(meanVec)) meanVec else meanVec[,i]
            rownames(x) = names(meanVec) = seq_along(meanVec)
            powMat = apply(x[,!(colnames(x) %in% Excl)], 2, getTopRank, power = pow,
                        maxExt = max(extremes), true = meanVec, simplify = FALSE)
            c(powMat, if(includeOut) list("estOutSplit" = t(sapply(extremes, function(ext){
                errorLarge(ests = x[, "estOutSplit"], ests2rank = x[, "estInSplit"],
                                 trueVec = meanVec, ext, p = p, power = pow)
            }))))
        })
        moltErrorList = melt(errorList, value.name = "Diagnostic")
        names(moltErrorList)[c(1:2, 4:5)] = c("Top_features", "extreme", "Method", "Instance")
        moltErrorList$Top_features = as.integer(moltErrorList$Top_features)
        moltErrorList$Method = factor(moltErrorList$Method, levels = methodLevels, labels = methodLabels)
        moltErrorList = droplevels(moltErrorList[moltErrorList$Method %in% Incl,])
        #The means
        meansDf = aggregate(Diagnostic ~ Method + extreme + Top_features, FUN = mean, data =  moltErrorList)
        meansDf$measure = switch(pow, "1" = "Bias", "2" = "MSE")
        moltMeansDf = melt(meansDf, value.name = "Diagnostic")
        names(moltMeansDf)[2:3] = c("Which", "Top_features")
        return(list("moltErrorList" = moltErrorList, "meansDf" = meansDf))
    })
    if(covWidth <- !is.null(estsMat[[1]]$ciList)){
        covList = lapply(seq_along(estsMat), function(i){
            x = estsMat[[i]]
            meanVec = if(is.vector(meanVec)) meanVec else meanVec[,i]
            names(meanVec) = seq_along(meanVec)
            idNames = grep(names(x$ciList), pattern = "ashBoot", invert = TRUE)
            x$ciList[idNames] = lapply(x$ciList[idNames], function(xx) {rownames(xx) = names(meanVec);xx})
            rownames(x$resMat) = names(meanVec)
            covMat = lapply(nam <- names(x$ciList)[!(names(x$ciList) %in% c("estInSplit", "bootCorrectForde", methodLevels[!(methodLabels %in% Incl)]))], function(nn){
                lapply(measures, function(m){
                    getTopCov(x$resMat[,nn], x$ciList[[nn]], ret = m, maxExt = max(extremes), true = meanVec,
                              ests2rank = switch(nn, "estOutSplit" = x$resMat[,"estInSplit"], x$resMat[,nn]))
                })
            })
            names(covMat) = nam
            covMat
        })
        moltcovList = melt(covList, value.name = "Diagnostic")
        names(moltcovList)[c(1:2, 4:6)] = c("Top_features", "extreme", "measure", "Method", "Instance")
        moltcovList$Top_features = as.integer(moltcovList$Top_features)
        moltcovList$Method = factor(moltcovList$Method, levels = methodLevels, labels = methodLabels)
        moltcovList = droplevels(moltcovList[moltcovList$Method %in% Incl,])
        #The means
        meansDfEr = aggregate(Diagnostic ~ Method + extreme + Top_features + measure, FUN = mean, data =  moltcovList)
    } else {
        meansDfEr = NULL
    }
    P = if(boxPlot){
        with(powList[[1]], {
        ggplot(data = moltErrorList, aes_string(y = "Diagnostic", colour = "Method", x ="Method")) +
        geom_boxplot() + facet_grid(extreme~Top_features, scales = "free_y") +
        geom_point(data = meansDf, shape = 5) +
            theme(axis.text.x = element_blank()) +if(power[1]==2) {scale_y_log10()}
        })
    } else {
        meansDf = rbind(Reduce(f = rbind, lapply(powList, function(x) x$meansDf)), meansDfEr)
        meansDf$measure = factor(meansDf$measure, ordered = TRUE, levels = c("Bias", "MSE", measures))
        if(returnDf){return(meansDf)}
        ggplot(data = meansDf[meansDf$extreme == "small", ], aes(y = Diagnostic, colour = Method, x =Top_features)) +
            geom_line(linewidth = lineSize) + facet_grid(measure~. , scales = "free_y")
    }
    intDat = data.frame("measure" = c("Bias", "MSE", if(covWidth) measures), "yintercept" = c(0, NA, if(covWidth) c(1-sigLevel, NA)))
    intDat$measure = factor(intDat$measure, ordered = TRUE, levels = c("Bias", "MSE", measures))
    P + geom_hline(data = intDat, aes(yintercept = yintercept), linetype = "dotted") +
        ylab(if(length(power)==1) {switch(power, "1" = "Bias", "2" = "MSE")} else {"Performance measure"}) +
        xlab("Number of extreme estimates") +
        scale_colour_manual(values = colourVec[names(colourVec) %in% Incl])
}
```
