## Supplementary material for "The winner’s curse under dependence: repairing empirical Bayes using convoluted densities": Data, R-code and style file: plotMSEres.R

```
plotMSEres = function(resList, Methods2Plot = plotNormal,
                      Palette = "Paired", Dodge = 0.75, colVecAdd = NULL){
    mseList = lapply(modes, function(mode){
        lapply(Sds, function(Sd){
            Sd = as.character(Sd)
            sapply(resList[[mode]][[Sd]], function(x){
                colMeans((x$resMat[, colnames(x$resMat) != "meanVec"] - x$resMat[, "meanVec"])^2)
            })
        })
    })
    moltErrorList = melt(mseList, value.name = "MSE")
    names(moltErrorList)[c(1:2,4,5)] = c("Method", "Instance", "Cor", "mode")
    moltErrorList$Cor = as.numeric(moltErrorList$Cor)
    moltErrorList = moltErrorList[moltErrorList$Method %in% c("estOutSplit", Methods2Plot),]
    moltErrorList$Method = factor(moltErrorList$Method, ordered = TRUE,
                                  levels = methodLevels, labels = methodLabels)
    moltErrorList = droplevels(moltErrorList)
    moltErrorList$Group = paste(with(moltErrorList, paste(LETTERS[as.integer(Method)], Cor)))
    ggplot(data = moltErrorList,
           aes(y = MSE, colour = Method, group = Group)) +
        facet_grid(Cor ~ mode, scales = 'free_y') +
        geom_boxplot() +
        stat_summary(fun = mean, aes(group = Group, x = 0), geom = "point", shape = 5, size = 1, position = position_dodge(Dodge)) +
        scale_colour_manual(values = colourVec[names(colourVec) %in% c(levels(moltErrorList$Method), colVecAdd)]) +
        scale_y_log10() + ylab("MSE of entire parameter vector") +
        xlab("Number of extreme estimates") +
        theme(axis.title.x = element_blank(), axis.ticks.x = element_blank(), axis.text.x = element_blank())
}
```
