## Supplementary material for "The winner’s curse under dependence: repairing empirical Bayes using convoluted densities": Data, R-code and style file: plotPredCI.R

```
plotPredCI = function(estsMat, meanVec, extremes = seq_len(30), measures = c("coverage", "width"),
                        Excl = c("estInSplit", methodLevels[!(methodLabels %in% Incl[Incl!="Separate estimation"])]),
                        Incl = namesCV, power = 1:2, boxPlot = FALSE, lineSize = 0.4){
    names(extremes) = extremes; names(measures) = measures
    p = length(meanVec)
    errorList = lapply(seq_along(estsMat), function(i){
        x = estsMat[[i]]
        meanVec = if(is.vector(meanVec)) meanVec else meanVec[,i]
        names(meanVec) = seq_along(meanVec)
        x$ciList = lapply(x$ciList, function(xx) {rownames(xx) = names(meanVec);xx})
        rownames(x$resMat) = names(meanVec)
        covMat = lapply(nam <- names(x$ciList)[!(names(x$ciList) %in% Excl)], function(nn){
            lapply(measures, function(m){
                getTopCov(x$resMat[,nn], x$ciList[[nn]], ret = m, maxExt = max(extremes), true = meanVec)
            })
        })
        names(covMat) = nam
        covMat
    })
    moltErrorList = melt(errorList, value.name = "Diagnostic")
    names(moltErrorList)[c(1:2, 4:6)] = c("Top_features", "extreme", "measure", "Method", "Instance")
    moltErrorList$Top_features = as.integer(moltErrorList$Top_features)
    moltErrorList$Method = factor(moltErrorList$Method, levels = methodLevels, labels = methodLabels)
    moltErrorList = droplevels(moltErrorList[moltErrorList$Method %in% Incl,])
    #The means
    meansDf = aggregate(Diagnostic ~ Method + extreme + Top_features + measure, FUN = mean, data =  moltErrorList)
    P = {
        ggplot(data = meansDf, aes_string(y = "Diagnostic", colour = "Method", x ="Top_features")) +
            geom_line(linewidth = lineSize) + facet_grid(measure~ extreme, scales = "free_y")
    }
    intDat = data.frame("measure" = "coverage", "yintercept" = 1-sigLevel)
    P + geom_hline(data = intDat, aes_string(yintercept = "yintercept"), linetype = "dotted") +
        ylab('Diagnostic') +
        scale_colour_manual(values = colourVec[names(colourVec) %in% unique(meansDf$Method)])
}
getTopCov = function(ests, ci, maxExt, true, ret, ests2rank = ests){
    orderMat = order(ests2rank)
    sortMat = ci[orderMat,]
    p = length(ests)
    mat = switch(ret,
                 "coverage" = cbind("small" = sortMat[seq_len(maxExt), 1] < true[orderMat][seq_len(maxExt)] & sortMat[seq_len(maxExt), 2] > true[orderMat][seq_len(maxExt)],
                                    "large" = sortMat[seq(p, p-maxExt+1), 1] < true[orderMat][seq(p, p-maxExt+1)] &
                                        sortMat[seq(p, p-maxExt+1), 2] > true[orderMat][seq(p, p-maxExt+1)]),
                 "width" = cbind("small" = apply(sortMat[seq_len(maxExt), ], 1, diff),
                                 "large" = apply(sortMat[seq(p, p-maxExt+1), ], 1, diff)))
    tmp = apply(mat, 2, cumsum)/seq_len(maxExt)
    rownames(tmp) = seq_len(maxExt)
    tmp
}
```
