## Supplementary material for "The winner’s curse under dependence: repairing empirical Bayes using convoluted densities": Data, R-code and style file: plotRunningTDP.R

```
plotRunningTDP = function(resList, extremes, p, methodLevels = methodLevels,
                          methodLabels = methodLabels,
                        Methods2Plot = plotNormal[!(plotNormal %in% c("condML"))], Palette = "Paired", aggFun = mean,
                         lineSize = 0.45, legSize = 8, boxPlot = FALSE, ltySlope = "dotted", rgSlope = FALSE, returnDf = FALSE){
    getInTdp = function(ests, maxExt, trueSort){
        sortMat = sort(ests)
            t(sapply(seq_len(maxExt), function(i){
            c("small" = mean(names(sortMat)[seq_len(i)] %in% names(trueSort)[seq_len(i)]),
              "large" = mean(names(sortMat)[length(sortMat)-seq_len(i)+1] %in% names(trueSort)[length(sortMat)-seq_len(i)+1]))
            }))
    }
errorMatList = lapply(modes, function(mode){
    lapply(Sds, function(Sd){
        Sd = as.character(Sd)
        tdpMats = lapply(resList[[mode]][[Sd]], function(x) {
            apply(x$resMat[,Methods2Plot[!(Methods2Plot %in% c("condML", "condML_rescale", "split", "catScores"))]], 2, getInTdp,
                  maxExt = max(extremes), true = sort(x$resMat[, "meanVec"]), simplify = FALSE)
        })
        lapply(extremes, function(ext){
            extChar =  as.character(ext)
            lapply(seq_along(resList[[mode]][[Sd]]), function(i){
                x = resList[[mode]][[Sd]][[i]]
                cbind(sapply(tdpMats[[i]], function(yy) {
                    c(yy[ext, "small"], yy[ext, "large"])
                }),
                "split" = runningTDP(ests = x$resMat[, "estInSplit"], ests2rank = x$resMat[, "estInSplit"],
                                     trueRanks = rank(x$resMat[, "meanVec"]), ext, p = p))
            })
        })
        })
})
    moltErrorList = melt(errorMatList, value.name = "Running_TDP")
    errorMatListZ = lapply(modes, function(mode){
        lapply(Sds, function(Sd){
        Sd = as.character(Sd)
        tdpMats = lapply(resList[[mode]][[Sd]], function(x) {
            apply(x$resMatZ[,grep(pattern = "rescale", value = TRUE, invert = TRUE, Methods2Plot[!(Methods2Plot %in% c("condML", "split", "Zstatistic", "ash", "ashBoot", "ashBootParam", "VanZwet2021", "VanZwet2021conv"))])], 2, getInTdp,
                  maxExt = max(extremes), true = sort(x$resMat[, "meanVec"]), simplify = FALSE)
        })
        lapply(extremes, function(ext){
            extChar =  as.character(ext)
            lapply(seq_along(resList[[mode]][[Sd]]), function(i){
                x = resList[[mode]][[Sd]][[i]]
                cbind(sapply(tdpMats[[i]], function(yy) {
                    c(yy[ext, "small"], yy[ext, "large"])
                }),
                "split" = runningTDP(ests = x$resMatZ[, "estInSplit"], ests2rank = x$resMatZ[, "estInSplit"],
                                     trueRanks = rank(x$resMat[, "meanVec"]), ext, p = p))
            })
        })
    })
})
    moltErrorListZ = melt(errorMatListZ, value.name = "Running_TDP")
    names(moltErrorList)[c(1:2, 4:7)] = names(moltErrorListZ)[c(1:2, 4:7)] = c("extreme","Method", "Instance", "Top_features", "Cor", "mode")
    moltErrorList$what = "Raw estimate"; moltErrorListZ$what = "Test statistic"
    moltErrorListFull = rbind(moltErrorList, moltErrorListZ)
    moltErrorListFull = moltErrorListFull[moltErrorListFull$Method %in% Methods2Plot,]
    moltErrorListFull$Cor = as.numeric(moltErrorListFull$Cor)
    moltErrorListFull = droplevels(moltErrorListFull)
    moltErrorListFull$Group = paste(with(moltErrorListFull, paste(as.integer(Method), Cor)))
    moltErrorListFull$Top_features = as.integer(moltErrorListFull$Top_features)
    moltErrorListFull$Method = factor(moltErrorListFull$Method, ordered = TRUE,
                                      levels = methodLevels, labels = methodLabels)
    #The means
    meansDf = aggregate(Running_TDP ~ Method + extreme + Top_features+ Cor + what + mode, FUN = aggFun, data = moltErrorListFull)
    # Add random guessing slope
    Slope = 1/p
    if(boxPlot){
        ggplot(data = moltErrorListFull,
        aes_string(y = "Running_TDP", colour = "Method", x ="Cor", group = "Group")) +
        geom_boxplot() + facet_grid(extreme~Top_features, scales = "free_y") +
        geom_hline(yintercept = 0, linetype ="dotted") +
        stat_summary(fun = mean, aes_string(group = "Group", x = "Cor"), geom = "point", shape = 5, size = 1, position = position_dodge(0.19)) +
        scale_color_brewer(palette = Palette) + guides(colour = guide_legend(nrow = 3)) +
        theme(legend.position = 'top', legend.text = element_text(size = legSize))
    } else {
        if(returnDf){
            meansDf$Performance_measure = "TDR"
            meansDf$value = meansDf$Running_TDP
            meansDf$Running_TDP = NULL
            return(meansDf[meansDf$extreme=="small", ])
        } else {
    ggplot(data = meansDf[meansDf$extreme=="small", ], aes(y = Running_TDP, colour = Method, x =Top_features, linetype = what)) +
        geom_line(linewidth = lineSize) + facet_grid(Cor ~ ., scales = "free_y") +
        ylab("TDR") + xlab("Number of extreme estimates") +
        scale_linetype(name = "Estimand") +
        scale_colour_manual(values = colourVec[names(colourVec) %in% levels(moltErrorListFull$Method)]) +
         if(rgSlope) {geom_abline(slope = Slope, intercept = 0, linetype = ltySlope)}
        }
    }
}
plotRunningTDPfdr = function(resList, extremes, p, methodLevels = methodLevels,
                          methodLabels = methodLabels,
                          Methods2Plot = plotNormal[!(plotNormal %in% c("condML"))], Palette = "Paired", aggFun = mean,
                          lineSize = 0.45, legSize = 8, boxPlot = FALSE, ltySlope = "dotted", rgSlope = FALSE, returnDf = FALSE){
    getInTdp = function(ests, maxExt, trueSort){
        sortMat = sort(ests)
        t(sapply(seq_len(maxExt), function(i){
            c("small" = mean(names(sortMat)[seq_len(i)] %in% names(trueSort)[seq_len(i)]),
              "large" = mean(names(sortMat)[length(sortMat)-seq_len(i)+1] %in% names(trueSort)[length(sortMat)-seq_len(i)+1]))
        }))
    }
    errorMatList = lapply(modes, function(mode){
        lapply(Sds, function(Sd){
            Sd = as.character(Sd)
            tdpMats = lapply(resList[[mode]][[Sd]], function(x) {
                extremes = getExtremes(x)
                apply(x$resMat[,Methods2Plot[!(Methods2Plot %in% c("condML", "condML_rescale", "split", "catScores"))]], 2, getInTdp,
                      maxExt = max(extremes), true = sort(x$resMat[, "meanVec"]), simplify = FALSE)
            })
            tmp = lapply(seq_along(resList[[mode]][[Sd]]), function(i){
                extremes = NROW(tdpMats[[i]][[1]]);names(extremes) = extremes;extChar = as.character(extremes)
                x = resList[[mode]][[Sd]][[i]]
                if(NCOL(tdpMats[[i]][[1]])==0){
                    return(NULL)
                }
                cbind(sapply(tdpMats[[i]], function(yy) {
                    c(yy[extremes, "small"], yy[extremes, "large"])
                }),
                "split" = runningTDP(ests = x$resMat[, "estInSplit"], ests2rank = x$resMat[, "estInSplit"],
                                     trueRanks = rank(x$resMat[, "meanVec"]), extremes, p = p))
            })
            tmp[!sapply(tmp, is.null)]
        })
    })
    moltErrorList = melt(errorMatList, value.name = "value")
    names(moltErrorList)[c(1:2, 4:6)] = c("extreme","Method", "Instance", "Cor", "mode")
    moltErrorList$what = "Raw estimate"
    moltErrorList = moltErrorList[moltErrorList$Method %in% Methods2Plot,]
    moltErrorList$Cor = as.numeric(moltErrorList$Cor)
    moltErrorList = droplevels(moltErrorList)
    moltErrorList$Group = paste(with(moltErrorList, paste(as.integer(Method), Cor)))
    moltErrorList$Method = factor(moltErrorList$Method, ordered = TRUE,
                                      levels = methodLevels, labels = methodLabels)
    #The means
    meansDf = aggregate(value ~ Method + extreme + Cor + what + mode, FUN = aggFun, data = moltErrorList)
    meansDf$Performance_measure = "TDR"
    meansDf
}
```
