## Supplementary material for "The winner’s curse under dependence: repairing empirical Bayes using convoluted densities": Data, R-code and style file: plotSimpleRes.R

```
plotSimpleRes = function(resList, extremes, p,
                         Methods2Plot = plotNormal, Palette = "Paired",
                         lineSize = 0.55, legSize = 8, power = 1, boxPlot = FALSE,
                         extPlot = "small", returnDf = FALSE, methodLevels = methodLevels,
                         methodLabels = methodLabels, notMethodsPlot = c("condML","condML_rescale", "split", "catScores")){
    errorMatList = lapply(modes, function(mode){
        lapply(Sds, function(Sd){
            Sd = as.character(Sd)
            topMats = lapply(resList[[mode]][[Sd]], function(x) {
                apply(x$resMat[,Methods2Plot[!(Methods2Plot %in% notMethodsPlot)]], 2, getTopRank, power = power,
                           maxExt = max(extremes), true = x$resMat[, "meanVec"], simplify = FALSE)
                })
         lapply(extremes, function(ext){
                extChar =  as.character(ext)
                lapply(seq_along(resList[[mode]][[Sd]]), function(i){
                    x = resList[[mode]][[Sd]][[i]]
                    t(rbind(t(sapply(topMats[[i]], function(yy) {
                        c("small" = yy[ext, "small"], "large" = yy[ext, "large"])
                        })),
        "condML" = c("small" = mean((x$condMLests[[extChar]]$small[, "condMLests"] - x$resMat[as.integer(rownames(x$condMLests[[extChar]]$small)), "meanVec"])^power),
            "large" = mean((x$condMLests[[extChar]]$large[, "condMLests"] - x$resMat[as.integer(rownames(x$condMLests[[extChar]]$large)), "meanVec"])^power)),
        "condML_rescale" = if(!is.null(x$condML_rescale)) {c("small" = mean((x$condML_rescale[[extChar]]$small[, "condMLests"] - x$resMat[as.integer(rownames(x$condML_rescale[[extChar]]$small)), "meanVec"])^power),
                     "large" = mean((x$condML_rescale[[extChar]]$large[, "condMLests"] - x$resMat[as.integer(rownames(x$condML_rescale[[extChar]]$large)), "meanVec"])^power))},
        "split" = errorLarge(ests = x$resMat[, "estOutSplit"], ests2rank = x$resMat[, "estInSplit"],
                             trueVec = x$resMat[, "meanVec"], ext, p = p, power = power))
                )})
            })
    })
        })
    moltErrorList = melt(errorMatList, value.name = "Estimation_error")
    #Test statistics
    errorMatListZ = lapply(modes, function(mode){
        lapply(Sds, function(Sd){
            Sd = as.character(Sd)
            topMats = lapply(resList[[mode]][[Sd]], function(x) {
                apply(x$resMatZ[,grep(pattern = "rescale", value = TRUE, invert = TRUE,
                                      Methods2Plot[!(Methods2Plot %in% c("condML","condML_rescale", "split", "VanZwet2021", "VanZwet2021conv"))])],
                      2, getTopRank, power = power,
                      maxExt = max(extremes), true = x$resMat[, "meanVec"]/sqrt(x$resMat[, "VarsTrue"]), simplify = FALSE)
            })
            lapply(extremes, function(ext){
                extChar =  as.character(ext)
                lapply(seq_along(resList[[mode]][[Sd]]), function(i){
                    x = resList[[mode]][[Sd]][[i]]
                    trueZ = x$resMat[, "meanVec"]/sqrt(x$resMat[, "VarsTrue"])
                    t(rbind(t(sapply(topMats[[i]], function(yy) {
                        c("small" = yy[ext, "small"], "large" = yy[ext, "large"])
                    })),
                    "condML" = c("small" = mean((x$condMLestsZ[[extChar]]$small[, "condMLests"] - trueZ[as.integer(rownames(x$condMLestsZ[[extChar]]$small))])^power),
                                 "large" = mean((x$condMLestsZ[[extChar]]$large[, "condMLests"] - trueZ[as.integer(rownames(x$condMLestsZ[[extChar]]$large))])^power)),
                    "split" = errorLarge(ests = x$resMatZ[, "estOutSplit"], ests2rank = x$resMatZ[, "estInSplit"],
                                         trueVec = trueZ, ext, p = p, power = power))
                    )
                })
            })
        })
    })
    moltErrorListZ = melt(errorMatListZ, value.name = "Estimation_error")
    names(moltErrorListZ)[c(1:2, 4:7)] = names(moltErrorList)[c(1:2, 4:7)] = c("extreme","Method", "Instance", "Top_features", "Cor", "mode")
    moltErrorList$what = "Raw estimate";moltErrorListZ$what = "Test statistic"
    moltErrorListFull = rbind(moltErrorList, moltErrorListZ)

    moltErrorListFull = moltErrorListFull[moltErrorListFull$Method %in% Methods2Plot,]
    moltErrorListFull$Cor = as.numeric(moltErrorListFull$Cor)
    moltErrorListFull$Method = factor(moltErrorListFull$Method, ordered = TRUE,
                                    levels = methodLevels, labels = methodLabels)
    moltErrorListFull = droplevels(moltErrorListFull)
    moltErrorListFull$Group = paste(with(moltErrorListFull, paste(as.integer(Method), Cor)))
    moltErrorListFull$Top_features = as.integer(moltErrorListFull$Top_features)
    #The means
    meansDf = aggregate(Estimation_error ~ Method + extreme + Top_features+ Cor +what + mode, FUN = mean, data =  moltErrorListFull)
    meansDf$Group = factor(paste(with(meansDf, paste(Method, Cor))), ordered = TRUE)
    if(boxPlot){
        ggplot(data = moltErrorListFull,
        aes_string(y = "Estimation_error", colour = "Method", x ="Cor", group = "Group")) +
        geom_boxplot() + facet_grid(extreme~Top_features, scales = "free_y") +
        geom_hline(yintercept = 0, linetype = "dotted") +
        stat_summary(fun = mean, aes_string(group = "Group", x = "Cor"), geom = "point", shape = 5, size = 1, position = position_dodge(0.19)) +
        scale_color_brewer(palette = Palette) + guides(colour = guide_legend(nrow = 3)) +
        theme(legend.position = 'top', legend.text = element_text(size = legSize))
    } else {
        if(returnDf){
            meansDf$Performance_measure = switch(power, "1" = "Bias", "2" = "MSE")
            meansDf$value = meansDf$Estimation_error
            meansDf$Estimation_error = meansDf$Group = NULL
            return(meansDf[meansDf$extreme == extPlot, ])
        } else {
    ggplot(data = meansDf[meansDf$extreme == extPlot, ], aes(y = Estimation_error, colour = Method, x = Top_features)) +
        geom_line(linewidth = lineSize) + facet_grid(what ~ Cor, scales = "free_y") +
        ylab(switch(power, "1" = "Bias", "2" = "MSE")) +
        xlab("Number of extreme estimates") +
        scale_colour_manual(values = colourVec[names(colourVec) %in% levels(moltErrorListFull$Method)]) +
        if(power==1) geom_hline(yintercept = 0, linetype = "dotted")
        }
    }
}
getTopRank = function(ests, maxExt, true, power, orderVec){
    if(missing(orderVec))
        orderVec = order(ests)
    sortMat = ests[orderVec]
    tmp = apply(cbind("small" = (sortMat[seq_len(maxExt)]- true[orderVec][seq_len(maxExt)])^power,
                "large" = (sortMat[seq(length(sortMat), length(sortMat)-maxExt+1)]- true[orderVec][seq(length(sortMat), length(sortMat)-maxExt+1)])^ power),
          2, cumsum)/seq_len(maxExt)
    if(!is.matrix(tmp)){
        tmp = t(tmp)
    }
     rownames(tmp) = seq_len(maxExt)
    tmp
}
```
