## Supplementary material for "The winner’s curse under dependence: repairing empirical Bayes using convoluted densities": Data, R-code and style file: plotSimpleResFDR.R

```
#Make a plot,but only for significant features
plotSimpleResFDR = function(resList, p,
                         Methods2Plot = plotNormal, Palette = "Paired",
                         lineSize = 0.55, legSize = 8, power = 1, boxPlot = FALSE,
                         extPlot = "small", returnDf = FALSE, methodLevels = methodLevels,
                         methodLabels = methodLabels, notMethodsPlot = c("condML","condML_rescale", "split", "catScores")){
    errorMatList = lapply(modes, function(mode){
        mclapply(mc.cores = nCores, Sds, function(Sd){
            Sd = as.character(Sd)
            topMats = lapply(resList[[mode]][[Sd]], function(x) {
                extremes = getExtremes(x)
                apply(x$resMat[,Methods2Plot[!(Methods2Plot %in% notMethodsPlot)]], 2, getTopRank, power = power,
                      maxExt = extremes, true = x$resMat[, "meanVec"], simplify = FALSE)
            })
            tmp = lapply(seq_along(resList[[mode]][[Sd]]), function(i){
                x = resList[[mode]][[Sd]][[i]]
                if(NCOL(topMats[[i]][[1]])==0){
                    return(NULL)
                }
                extremes = NROW(topMats[[i]][[1]]);names(extremes) = extremes;extChar = as.character(extremes)
                 condMLests = corrCondML(raw = x$resMat[, "rawEst"], vars = apply(x$mat,2, var)/nrow(x$mat), dat = x$mat, extremes, ciReturn = TRUE)
                t(rbind(t(sapply(topMats[[i]], function(yy) {
                    c("small" = yy[extremes, "small"], "large" = yy[extremes, "large"])
                })),
                "condML" = c("small" = mean((condMLests[[extChar]]$small[, "condMLests"] - x$resMat[as.integer(rownames(condMLests[[extChar]]$small)), "meanVec"])^power),
                             "large" = mean((condMLests[[extChar]]$large[, "condMLests"] - x$resMat[as.integer(rownames(condMLests[[extChar]]$large)), "meanVec"])^power)),
                "split" = errorLarge(ests = x$resMat[, "estOutSplit"], ests2rank = x$resMat[, "estInSplit"],
                                     trueVec = x$resMat[, "meanVec"], extremes, p = p, power = power))
                )
                })
            tmp[!sapply(tmp, is.null)]
        })
    })
    moltErrorList = melt(errorMatList, value.name = "Estimation_error")
    names(moltErrorList)[c(1:2, 4:6)] = c("extreme","Method", "Instance", "Cor", "mode")
    moltErrorList$what = "Raw estimate";
    moltErrorList = moltErrorList[moltErrorList$Method %in% Methods2Plot,]
    moltErrorList$Cor = as.numeric(moltErrorList$Cor)
    moltErrorList$Method = factor(moltErrorList$Method, ordered = TRUE,
                                      levels = methodLevels, labels = methodLabels)
    moltErrorList = droplevels(moltErrorList)
    moltErrorList$Group = paste(with(moltErrorList, paste(as.integer(Method), Cor)))
    #The means
    meansDf = aggregate(Estimation_error ~ Method + extreme + mode + Cor +what, FUN = mean, data =  moltErrorList)
    meansDf$Group = factor(paste(with(meansDf, paste(Method, Cor))), ordered = TRUE)
    if(returnDf){
            meansDf$Performance_measure = switch(power, "1" = "Bias", "2" = "MSE")
            meansDf$value = meansDf$Estimation_error
            meansDf$Estimation_error = meansDf$Group = NULL
            return(meansDf[meansDf$extreme == extPlot, ])
     } else {
            ggplot(data = meansDf[meansDf$extreme == extPlot, ], aes(y = Estimation_error, colour = Method, x = Top_features)) +
                geom_line(linewidth = lineSize) + facet_grid(what ~ Cor, scales = "free_y") +
                ylab(switch(power, "1" = "Bias", "2" = "MSE")) +
                xlab("Number of extreme estimates") +
                scale_colour_manual(values = colourVec[names(colourVec) %in% levels(moltErrorList$Method)]) +
                if(power==1) geom_hline(yintercept = 0, linetype = "dotted")
        }
}
getExtremes = function(x){
    zStats = x$resMatZ[, "rawEst"]
    pVals = pnorm(zStats);pVals[zStats > 0] = pnorm(zStats[zStats > 0], lower.tail = FALSE)
    pAdj = p.adjust(pVals*2, method = "BH")
    sum(pAdj < sigLevel & zStats < 0) #Only small ranks matter
}
```
