## Supplementary material for "The winner’s curse under dependence: repairing empirical Bayes using convoluted densities": Data, R-code and style file: plotWidthCI.R

```
plotWidthCI = function(resList, extremes, Palette = "Paired",
                        Methods2Plot = c(plotNormal, "estOutSplit"),
                        lineSize = 0.55, legSize = 8, returnDf = FALSE, extPlot = "small",
                       methodLevels = methodLevels, methodLabels = methodLabels){
    widthList = lapply(modes, function(mode){
        lapply(Sds, function(Sd){
            Sd = as.character(Sd)
            topMats = lapply(resList[[mode]][[Sd]], function(x) {
                out = lapply(nm <- names(x$ciList)[!sapply(x$ciList, is.null)], function(m){
                    Ord = order(x$resMat[, switch(m, "estOutSplit" = "estInSplit", m)])
                    Width = x$ciList[[m]][,2] - x$ciList[[m]][,1]
                    cbind("small" = cumsum(Width[Ord][extremes]),
                              "large" = cumsum(Width[Ord][length(Width)-extremes+1]))/seq_along(extremes)
                })
                names(out) = nm
                out
            })
            lapply(extremes, function(ext){
                extChar = as.character(ext)
                lapply(seq_along(resList[[mode]][[Sd]]), function(i){
                    x = resList[[mode]][[Sd]][[i]]
                    t(rbind(t(sapply(topMats[[i]], function(yy) {
                        c(yy[ext, "small"], yy[ext, "large"])
                    })),
                    "condML" = c("small" = mean(x$condMLests[[extChar]]$small[, 3] - x$condMLests[[extChar]]$small[, 2]),
                                 "large" = mean(x$condMLests[[extChar]]$large[, 3] - x$condMLests[[extChar]]$large[, 2])),
                    "condML_rescale" = c("small" = mean(x$condML_rescale[[extChar]]$small[, 3] - x$condML_rescale[[extChar]]$small[, 2]),
                                                      "large" = mean(x$condML_rescale[[extChar]]$large[, 3] - x$condML_rescale[[extChar]]$large[, 2]))
                    ))
                    })
            })
        })
    })
    moltErrorList = melt(widthList, value.name = "Width")
    widthListZ = lapply(modes, function(mode){
        lapply(Sds, function(Sd){
            Sd = as.character(Sd)
            topMats = lapply(resList[[mode]][[Sd]], function(x) {
                out = lapply(nm <- names(x$ciListZ)[!sapply(x$ciListZ, is.null)], function(m){
                    Ord = order(x$resMatZ[, switch(m, "estOutSplit" = "estInSplit", m)])
                    Width = x$ciListZ[[m]][,2] - x$ciListZ[[m]][,1]
                    cbind("small" = cumsum(Width[Ord][extremes]),
                          "large" = cumsum(Width[Ord][length(Width)-extremes+1]))/seq_along(extremes)
                })
                names(out) = nm
                out
            })
            lapply(extremes, function(ext){
                extChar =  as.character(ext)
                lapply(seq_along(resList[[mode]][[Sd]]), function(i){
                    x = resList[[mode]][[Sd]][[i]]
                    t(rbind(t(sapply(topMats[[i]], function(yy) {
                        c(yy[ext, "small"], yy[ext, "large"])
                    })),
                    "condML" = c("small" = mean(x$condMLestsZ[[extChar]]$small[, 3] - x$condMLestsZ[[extChar]]$small[, 2]),
                                 "large" = mean(x$condMLestsZ[[extChar]]$large[, 3] - x$condMLestsZ[[extChar]]$large[, 2]))
                    ))
                })
            })
        })
    })
    moltErrorListZ = melt(widthListZ, value.name = "Width")
    names(moltErrorList)[c(1:2, 4:7)] =  names(moltErrorListZ)[c(1:2, 4:7)] = c("extreme","Method", "Instance", "Top_features", "Cor", "mode")
    moltErrorList$what = "Raw estimate";moltErrorListZ$what = "Test statistic"
    moltErrorListFull = rbind(moltErrorList, moltErrorListZ)
    moltErrorListFull = moltErrorListFull[moltErrorListFull$Method %in% Methods2Plot,]
    moltErrorListFull$Cor = as.numeric(moltErrorListFull$Cor)
    moltErrorListFull$Method = factor(moltErrorListFull$Method, ordered = TRUE,
                                      levels = methodLevels, labels = methodLabels)
    moltErrorListFull = droplevels(moltErrorListFull)
    moltErrorListFull$Top_features = as.integer(moltErrorListFull$Top_features)
    #The means
    meansDf = aggregate(Width ~ Method + extreme + Top_features + Cor + what + mode, FUN = mean, data =  moltErrorListFull)
    if(returnDf){
        meansDf$Performance_measure = "Width"
        meansDf$value = meansDf$Width
        meansDf$Width = NULL
        return(meansDf[meansDf$extreme == extPlot, ])
    } else {
        ggplot(data = meansDf, aes_string(y = "Width", colour = "Method", x = "Top_features")) +
                geom_line(size = lineSize) + facet_grid(extreme ~ Cor, scales = "free_y") +
            xlab("Number of extreme estimates") +
            scale_colour_manual(values = colourVec[names(colourVec) %in% unique(meansDf$Method)])
    }
}
plotWidthCIfdr = function(resList, Palette = "Paired",
                       Methods2Plot = c(plotNormal, "estOutSplit"),
                       lineSize = 0.55, legSize = 8, returnDf = FALSE, extPlot = "small",
                       methodLevels = methodLevels, methodLabels = methodLabels){
    widthList = lapply(modes, function(mode){
        lapply(Sds, function(Sd){
            Sd = as.character(Sd)
            topMats = lapply(resList[[mode]][[Sd]], function(x){
                extremes = getExtremes(x)
                out = lapply(nm <- names(x$ciList)[!sapply(x$ciList, is.null)], function(m){
                    Ord = order(x$resMat[, switch(m, "estOutSplit" = "estInSplit", m)])
                    Width = x$ciList[[m]][,2] - x$ciList[[m]][,1]
                    cbind("small" = cumsum(Width[Ord][extremes]),
                          "large" = cumsum(Width[Ord][length(Width)-extremes+1]))/seq_along(extremes)
                })
                names(out) = nm
                out
            })
            lapply(seq_along(resList[[mode]][[Sd]]), function(i){
                    x = resList[[mode]][[Sd]][[i]]
                    extremes = NROW(topMats[[i]][[1]]);names(extremes) = extremes;extChar = as.character(extremes)
                    if(NCOL(topMats[[i]][[1]])==0){
                        return(NULL)
                    }
                    t(rbind(t(sapply(topMats[[i]], function(yy) {
                        if(!NROW(yy) || NCOL(yy) <=1){
                            return(c("small" = NA, "large" = NA))
                        } else {
                            c("small" = yy[1, "small"], "large" = yy[1, "large"])
                        }
                    })),
                    "condML" = c("small" = mean(x$condMLests[[extChar]]$small[, 3] - x$condMLests[[extChar]]$small[, 2]),
                                 "large" = mean(x$condMLests[[extChar]]$large[, 3] - x$condMLests[[extChar]]$large[, 2]))
                    ))
                })
            })
    })
    moltErrorList = melt(widthList, value.name = "value")
    names(moltErrorList)[c(1:2, 4:6)] = c("extreme","Method", "Instance", "Cor", "mode")
    moltErrorList$what = "Raw estimate"
    moltErrorList = moltErrorList[moltErrorList$Method %in% Methods2Plot,]
    moltErrorList$Cor = as.numeric(moltErrorList$Cor)
    moltErrorList$Method = factor(moltErrorList$Method, ordered = TRUE,
                                      levels = methodLevels, labels = methodLabels)
    moltErrorList = droplevels(moltErrorList)
    #The means
    meansDf = aggregate(value ~ Method + extreme + Cor + what + mode, FUN = mean, data =  moltErrorList)
    meansDf$Performance_measure = "Width"
    meansDf
}
```
