## Supplementary material for "The winner’s curse under dependence: repairing empirical Bayes using convoluted densities": Data, R-code and style file: predLin.R

```
#Fit linear model and make prediction
predLin = function(x, y, xnew, loc = NULL, ...){
    if(!is.null(loc)){
        dat = data.frame("a" = y, "b" = c(x), loc)
        glsFit = try(gls(model = formula("a ~ b"), data = dat, na.action = na.exclude, ...), silent = TRUE)
    }
    if(is.null(loc) || inherits(glsFit, "try-error")) {
        predOLS(x, y, xnew)
    } else {
        glsFit$coef[1] + xnew * glsFit$coef[2]
    }
}
predGlmnet = function(x,y, xnew, pengls = FALSE, alpha, loc, innerCV = TRUE, ...){
    predFun = match.fun(paste0(if(innerCV) {"cv."}, if(pengls) "pengls" else "glmnet"))
    glmNetFit = if(pengls){
            predFun(data = data.frame(x, "y" = y, loc),
                  xNames = colnames(x), outVar = "y", alpha = alpha, ...)
        } else {
            predFun(x = x, y = y, alpha = alpha, ...)
    }
    predict(glmNetFit, newx =  xnew)
}
predOLS = function(x, y, xnew){
    lmFit = lm.fit(x = cbind(1, x), y = y)
    lmFit$coef[1] + xnew * lmFit$coef[2]
}
```
