## Supplementary material for "The winner’s curse under dependence: repairing empirical Bayes using convoluted densities": Data, R-code and style file: RCTshrinkage.R

```
#Fit mixture of 4 gaussians
fitMix = function(z, int = 1, k = 4, ...){
    flexmod = flexmix(formula(paste("z~", int)), k = k)#zero mean
    param = parameters(flexmod)
    rbind("sigma" = if(int==0) param else param["sigma",], "probs" = summary(flexmod)@comptab$prior,
          "mean" = if(int==0) 0  else param["coef.(Intercept)",])
}
#Do not go for unimodality, unlike ash
#Shrinkage according to van Zwet 2021
conditional <- function(fitMixMod, z, s,...) {
    condRes = lapply(seq_along(z), function(i){
        tau2 <- fitMixMod["sigma",]^2
        q <- fitMixMod["probs",]*dnorm(z[i],fitMixMod["mean",],sqrt(tau2+1))
        q <- q/sum(q)
        # conditional mixing probs
        m <- z[i]*tau2/(tau2+1)
        # conditional means
        v <- tau2/(tau2+1)
        # conditional variances
        sigma <- sqrt(v)
        # conditional std devs
        cbind("q" = q, "m" = m, "sigma" = sigma)# betahat = s*E(SNR | z)
    })
    betaHats = vapply(seq_along(z), FUN.VALUE = 0, function(i){
        "betahat" = s[i] * sum(condRes[[i]][, "q"] * condRes[[i]][, "m"])
    })
    names(betaHats) = names(condRes) = names(z)
    list("condRes" = condRes, "betaHats" = betaHats, "s" = s)
}
zwetConf = function(res, s, sigLevel = 0.05){
    #Confidence interval
    confInts = vapply(seq_along(res$condRes), FUN.VALUE = double(2), function(i){
        KScorrect::qmixnorm(p = c(sigLevel/2, 1-sigLevel/2), mean = res$condRes[[i]][, "m"],
                            sd = res$condRes[[i]][,"sigma"], pro = res$condRes[[i]][, "q"], expand=0.5)*s[i]
        #Multiply by S for confidence interval on beta
    })
}
zwetWrapper = function(z, s, zEsts = z, confInt = TRUE, int = 1, ...){
    fitMixMod = fitMix(zEsts, int = int, ...)
    res = conditional(fitMixMod, z, s)
    confInts = if(confInt){t(zwetConf(res, s))}
    list("res" = res, "confInts" = confInts)
}
```
