## Supplementary material for "The winner’s curse under dependence: repairing empirical Bayes using convoluted densities": Data, R-code and style file: simpleCV.R

```
#' Perform simple CV, returning a list of cv matrices per feature
simpleCV = function(dat, nOuterFolds, loc = NULL, nCores = 1, ...){
    folds = sample(rep(unFolds <- seq_len(nOuterFolds), length.out = nrow(dat$x)))
    mclapply(mc.cores = nCores, seq_len(ncol(dat$x)), function(j){
        sapply(unFolds, function(uf){
            idTrain = folds!=uf
            predTest = predLin(dat$x[idTrain,j], dat$y[idTrain],
                               dat$x[!idTrain,j], loc = loc[idTrain,], ...)
            (predTest-dat$y[!idTrain])^2
        })
    })
}
#' single CV for glmnet
singleCVglmnet = function(trainDat, nOuterFolds, pengls = FALSE, loc = NULL, alpha, ...){
    folds = sample(rep(unFolds <- seq_len(nOuterFolds), length.out = nrow(trainDat$x)))
    sapply(unFolds, function(uf){
        idTrain = folds!=uf
        glmNetFit = if(pengls){
            cv.pengls(data = data.frame(trainDat$x[idTrain,], "y" = trainDat$y[idTrain], loc[idTrain,]),
                      nfolds = nOuterFolds-1, xNames = colnames(trainDat$x), outVar = "y",  alpha = alpha,...)
        } else {
            cv.glmnet(x = trainDat$x[idTrain,], y = trainDat$y[idTrain], nfolds = nOuterFolds-1,  alpha = alpha, ...)
        }
        #Inner CV is considered part of model fitting
        (predict(glmNetFit, newx = trainDat$x[!idTrain,])-trainDat$y[!idTrain])^2
    })
}
```
