## Supplementary material for "The winner’s curse under dependence: repairing empirical Bayes using convoluted densities": Data, R-code and style file: singleCVEN.R

```
# Non-nested CV for EN
singleCVEN = function(penGrid, i, blockFolds, phenotypes, rlogBras3, coordsPlants, glsSt, folder, condVars = NULL, fold){
    resultFile = paste0("Results/", folder, "/", paste(penGrid[i, ], collapse = "_"),"fold", fold, ".RData")
    if(!file.exists(resultFile)){
        phenType = phenotypes[blockFolds$id, penGrid[i, "phenType"]]
        notNa = blockFolds$id[!is.na(phenType)]
        outerFoldVec =  switch(penGrid[i, "CV"],
                                 "random" = sample(blockFolds$fold[blockFolds$id %in% notNa]),
                                 "blocked" = blockFolds$fold[blockFolds$id %in% notNa])
        notNaExpr = rlogBras3[notNa,];phenType =  phenType[!is.na(phenType)]
        #Add variables for conditioning
        notNaExpr = cbind("condVar" = phenotypes[rownames(notNaExpr), condVars], notNaExpr)
        unFolds = unique(outerFoldVec); names(unFolds) = unFolds
        #A naive fit to provide a lambda sequence
        naieveFit <- cv.glmnet(x = as.matrix(notNaExpr), y = phenType,
                               foldid = outerFoldVec, alpha = penGrid[i, "alpha"])
        lambdas = naieveFit$lambda
        data = data.frame(notNaExpr, "phenType" = phenType, coordsPlants[rownames(notNaExpr), c("x", "y")])
        rownames(data) = rownames(notNaExpr)
        xNames = colnames(notNaExpr)
        innerPreds = lapply(unFolds, function(ifo){
            idif = outerFoldVec!=ifo #Leave out test fold
            Excl = filterFeat(data[idif,xNames], criterion = penGrid[i, "filter"], data[idif,"phenType"])
            if(length(Excl)==length(xNames)){
                return(matrix(ncol = length(lambdas), nrow = sum(!idif), rep(mean(data[idif,"phenType"]),sum(!idif)))) #If no predictors, return mean vector
            }
            innerFits = lapply(lambdas, function(lam){
                switch(penGrid[i, "algorithm"],
                       "pengls" = pengls(data = data[idif,], glsSt = glsSt, xNames = xNames,
                                         exclude = Excl+1, outVar = "phenType", lambda = lam, alpha = penGrid[i, "alpha"]),
                       "glmnet" = glmnet(x = data[idif,xNames], y = data[idif, "phenType"], exclude = Excl,
                                         lambda = lam, alpha = penGrid[i, "alpha"])
                )
            })
            coefs = vapply(FUN.VALUE = numeric(length(xNames)+1), innerFits, function(x) as.vector(coef(x)))
            preds = cbind(1, as.matrix(data[!idif,xNames, drop = FALSE])) %*% coefs
        })
        innerObs = lapply(unFolds, function(ifo){
            data[outerFoldVec==ifo, "phenType"] #only use test fold
        })
        MSEs = mapply(innerPreds, innerObs, FUN = function(p, o){
            colMeans((p-o)^2)
        })
        #Use lambda within 1se from minimum error
        cvEsts <- rowMeans(MSEs)
        maxId <- which.min(cvEsts) #minimum location
        sdMax <- sqrt(mean((MSEs[maxId,]-cvEsts[maxId])^2)/(nrow(MSEs)-1))
        cvId <- cvEsts < (cvEsts[maxId] + sdMax)
        seId <- which.max(cvId)
        innerPredsObs = mapply(innerPreds, innerObs, FUN = function(p, o) cbind("pred" = p[, seId], "obs" = o))
        ## END: inner loop
        #Now fit model with best lambda on complete dataset
        ExclIn = filterFeat(data[,xNames], criterion = penGrid[i, "filter"], data[,"phenType"])
        innerModel = try(silent = TRUE, switch(penGrid[i, "algorithm"],
                            "pengls" = pengls(data = data, glsSt = glsSt, xNames = xNames, exclude = ExclIn+1,
                                              outVar = "phenType", lambda = lambdas[seId], alpha = penGrid[i, "alpha"]),
                            "glmnet" = glmnet(x = data[,xNames], y = data[, "phenType"], exclude = ExclIn,
                                              lambda = lambdas[seId], alpha = penGrid[i, "alpha"])))
        singleModel = list("innerPredsObs" = innerPredsObs, "innerModel" = innerModel)
        save(singleModel, file = resultFile)
    } #else load(resultFile)
    #singleModel
}
```
