## Supplementary material for "The winner’s curse under dependence: repairing empirical Bayes using convoluted densities": Data, R-code and style file: TanCorrect.R

```
#'The correction by Tan2015 et al.
TanCorrect = function(rawEsts, bootEsts, quants, ciReturn = FALSE,
                      fullBootObj = NULL, testStatistic = FALSE){
    rankRaw = rank(rawEsts)
    orderBoot = apply(bootEsts, 1, order)
    sortBoot = apply(bootEsts, 1, sort)
    betaCor = vapply(FUN.VALUE = double(1), seq_along(rawEsts), function(k){ #Loop over ranks
        mean(sortBoot[k, ] - rawEsts[orderBoot[k,]])
    })
    out = rawEsts - betaCor[rankRaw]
    ciTan = if(ciReturn){
        tanCorrectIn = sapply(seq_len(dim(fullBootObj[[1]]$outIn)[3]), function(i){
            TanCorrect(rawEsts = bootEsts[i,], quants = quants, ciReturn = FALSE,
                        bootEsts = t(fullBootObj[[i]]$outIn[,"estsIn",]/
                                         (if(testStatistic) sqrt(fullBootObj[[i]]$outIn[,"varEstsIn",]) else 1)))$est[order(bootEsts[i,])]
        })
        biasIn = sapply(seq_len(nrow(bootEsts)), function(i){
            Ord = order(tanCorrectIn[,i])
            tanCorrectIn[Ord, i] - bootEsts[i, Ord]
        })
        biasQuants = t(apply(biasIn, 1, quantile, quants))
        ci = out + biasQuants[order(out),]
        names(ci) = names(out)
        ci
    }
    return(list("est" = out, "ci" = ciTan))
}
```
