## Supplementary material for "The winner’s curse under dependence: repairing empirical Bayes using convoluted densities": Data, R-code and style file: tweedieForm.R

```
#' Apply Tweedie's correction to the estimated AUC
#'
#' @param estimates Parameter estimates
#' @param varsEstimates Covariance matrix of parameter estimates
#' @param ... passed on to the density function
#' @return Tweedie's corrected estimates
tweedieForm = function(estimates, varsEstimates, estimatesForDens = estimates, n, smooth = FALSE,
                       densMethod = "Lindsey", analytical = TRUE, alpha1, trueMeans = NULL, trueVars = NULL, minAlpha1 = 0.05, ...) {
    densEstsLogDeriv = if(!analytical || alpha1 <= minAlpha1){
         estDens(estimatesForDens, zObs = estimates, densMethod = densMethod, sdVec = sqrt(varsEstimates), ...)
    } else {
        vapply(estimates, function(x) anaDerivLog(x, if(is.null(trueMeans)) estimates else trueMeans,
                                                  sqrt((if(is.null(trueVars)) varsEstimates else trueVars)*alpha1)),
               FUN.VALUE = double(2))
    }
    if(smooth){
        densEstsLogDeriv["deriv1", ] = splinPred(estimates, densEstsLogDeriv["deriv1", ], estimates, df = n)
    }
    tweedieCorEsts = estimates + varsEstimates * densEstsLogDeriv["deriv1", ]
    return(rbind("tweedieCorEsts" = tweedieCorEsts, "densEstsLogDeriv" = densEstsLogDeriv["deriv1", ],
                "densEstsLogDeriv2" = densEstsLogDeriv["deriv2",]))
}
estDens = function(densMethod, ...){
    switch(densMethod,
           "Lindsey" = LindseysMethod(...),
           "nlpden" = nlpdenWrapper(...),
           "kernel" = kernelEst(...),
           stop("Density estimation only implemented for 'Lindsey' and 'nlpden'"))
}
kernelEst = function(x, ...){
    deriv2kde = predict(kdde(estimates, deriv.order = 2),x = estimates)
    deriv1kde = predict(kdde(estimates, deriv.order = 1),x = estimates)
    deriv0kde = predict(kdde(estimates, deriv.order = 0),x = estimates)
    derivLogKde = deriv1kde/deriv0kde
    derivLogKde2 = (deriv2kde*deriv0kde - deriv1kde^2)/deriv0kde^2
    rbind("deriv1" = derivLogKde, "deriv2" = derivLogKde2)
}
```
