## Supplementary material for "The winner’s curse under dependence: repairing empirical Bayes using convoluted densities": Data, R-code and style file: SelectionBiasSup.Rmd

````
---
title: "Supplementary material for 'The winner's curse under dependence: repairing empirical Bayes using convoluted densities'"
author: "Stijn Hawinkel, Olivier Thas and Steven Maere"
output: 
  pdf_document:
    number_sections: true
    keep_tex: yes
    includes:
            in_header: selBias.sty
---

\beginsupplement{0}

```{r setup, include=FALSE}
knitr::opts_chunk$set(cache = TRUE, autodep = TRUE, warning = FALSE, cache.lazy = FALSE,
                      message = FALSE, echo = FALSE, eval = TRUE, tidy = TRUE,
                      fig.width = 7, fig.height = 4, purl = TRUE, 
                      fig.show = "hold", fig.pos = "p")
libs = c("ggplot2", "parallel", "reshape2", "nlme", "glmnet", "grid", "mvtnorm", 
         "xtable", "pengls", "numDeriv", "BiocParallel", "ks", "ashr", "nlpden",
         "st", "forcats", "winnerscurse", "KScorrect", "flexmix")
for (i in libs){
  library(i, character.only = TRUE, quietly = TRUE)
};rm(i, libs)
axisTitleSize = 11
theme_set(theme_bw())
theme_update(axis.title = element_text(size = axisTitleSize))
for(i in list.files("R")){source(file.path("R", i))};rm(i)
nCores = 3
lmeCon = lmeControl(opt = "optim", maxIter = 5e2, msVerbose = FALSE,
                   msMaxIter = 5e2, niterEM = 1e3, msMaxEval=1e3)
colourVec = c("rawEst" = "black", "split" = "#000099", "estOutSplit" = "grey25", "condML" = "#990099", 
              "FayeCorrectEst" = "#FF6600", "TanCorrectEst" = "#FFCC00", "TanCorrectEstParam" = "#CCCC99", "bootCorrectForde" = "#FF0099", "tweedieEst" = "#CCCC99", "tweedieEstsTrunc" = "#CCCC99",
              "tweedieConvEsts" = "#99CC00", "tweedieConvEstsTrunc" = "#99CC00","tweedieConvEstsBoot" = "#339900", "tweedieConvEstsTruncBoot" = "#660066", "ash" = "#990000", "ashConv" = "#FF0033", "VanZwet2021" = "#006666", "VanZwet2021conv" = "#00FFFF", "rawsCatBoot" = "#0099FF", "rawsCatBootParam" = "#009999", "decorrelated" = "#0099FF", "catScores" = "#0099FF", "Zstatistic" = "#666666")
methodLevels = names(colourVec)
methodLabels = c("Raw", "Separate estimation", "Separate estimation", "Conditional likelihood", 
                 "BR-squared", "Tan2015 (nonparametric)", "Tan2015 (parametric)", "winnerscurse", "Tweedie's formula", "Tweedie's formula\ntruncated", "Tweedie's formula\nconvoluted or bagged", "Tweedie's formula\nconvoluted or bagged\n(truncated)", 
                 "Tweedie's formula\nconvoluted or bagged(bootstrap SE)", "Tweedie's formula\nconvoluted or bagged\n(truncated, bootstrap SE)",  "Ash", "Ash convoluted", "VanZwet2021", "VanZwet2021 convoluted", "Cat scores\nnonparametric bootstrap", "Cat scores\nparametric bootstrap", "Decorrelated\nestimates", "Cat scores", "Z-statistic")
names(colourVec) = methodLabels
colourVec = c(colourVec, "Tweedie's formula\nconvoluted" = "#99CC00")
foo = lapply(c("Results", "simResults"), dir.create, showWarnings = FALSE) #Create paths to store results
options(tinytex.clean = FALSE) #Keep aux file for cross-referencing
```

\tableofcontents

The R-code for producing all output of this study is available from _Biostatistics_ online.

```{r simSetup}
n = 50 #Sample size
p = 1e3 #Number of features
nOuterFolds = 10
nInnerFolds = nOuterFolds - 1
sdY = 1 #Residual SD
alpha = 0.5
sigLevel = 0.05
Quants = c("lower"= -1, "upper" = 1)*qnorm(1-sigLevel/2)
probs <- c("lower" = sigLevel/2, "upper" = 1-sigLevel/2)
ciReps = 5e2 #Number of Monte-Carlo replicates for confidence interval coverage
MCestReps = 5e3 #Number of Monte-Carlo replicates to find true MSEs
cvSplits = 2e2 #Number of repeats of nested CV
signal = 2; fracSignal = 0.004
betas1e3 = c(rep(signal,p*fracSignal), rep(0, p*(1-fracSignal))) #Keep Betas fixed
tweedieReps = 1e2 #Number of times to repeat Tweedie's formula
cvSplitsBoot = 1;
bootReps = 1e2
nFolds = 10
sdMean = 0.2
extVec = seq_len(30); names(extVec) = extVec
ciRepsSingle = 5e1 #For univariate models there are several repeats => fewer reps suffice
betaList = list("p" = betas1e3)
Sds = c(0, 0.25, 0.5, 0.75); names(Sds) = Sds
```

# Convolution for improved density estimation under dependence

## The derivative of the logarithm of a mixture of normals \label{sec:logDeriv} 

The first and second derivative of the log density of an even mixture of $p$ normal distributions as in \eqref{eq:ftilde2} with means $\mu_j$, variances $\sigma_j$ and density $f_j(\gamma) = \frac{1}{\sigma_j\sqrt{2\pi}}\exp\left(-1/2(\frac{\gamma-\mu_j}{\sigma_j})^2\right)$ with $j=1,\hdots,p$ equal

\begin{equation}
\begin{aligned}
\frac{d \log f(\gamma)}{d\gamma} &= \frac{d \log p^{-1}\sum_{j=1}^pf_j(\gamma)}{d\gamma}\\
&= \frac{\sum_{j=1}^pf_j(\gamma)\frac{(\mu_j-\gamma)}{\sigma_j^2}}{\sum_{j=1}^pf_j(\gamma)}\\
\frac{d^2 \log f(\gamma)}{d\gamma^2} &= \frac{d}{d\gamma}\frac{\sum_{j=1}^pf_j(\gamma)\frac{(\mu_j-\gamma)}{\sigma_j^2}}{\sum_{j=1}^pf_j(\gamma)}\\ &= 
\frac{\sum_{j=1}^pf_j(\gamma)\left[\left(\frac{\mu_j-\gamma}{\sigma_j^2}\right)^2 - \frac{1}{\sigma_j^2} \right]}{\sum_{j=1}^pf_j(\gamma)}-\left(\frac{\sum_{j=1}^pf_j(\gamma)\frac{\mu_j-\gamma}{\sigma_j^2}}{\sum_{j=1}^pf_j(\gamma)}\right)^2
\label{eq:logDeriv2}
\end{aligned}
\end{equation}

## Scenarios in which convolution renders Tweedie's formula unbiased \label{sec:unbiased}

Although convolution yields an estimator $\tilde{f}$ with correct expected variance, unbiasedness of the estimation of the derivative of its log-density and hence of Tweedie's formula cannot be guaranteed in general. In computer simulations we see a great improvement in the bias though, suggesting that convolution is a simple and good working solution that drastically reduces bias in Tweedie's formula under many scenarios. Here we prove unbiasedness of our procedure in a few simple cases. We assume that M=1, so the covariance structure is determined by a single confounder that engenders a compound symmetric covariance structure, as is the case in our motivating example of estimating prediction loss for a single outcome using many possible predictors. In that case $\hat{k}$ is a normal distribution $\hat{k}(z) = r(z|\zeta \sqrt{\alpha_1}, 1-\alpha_1)$, with $\zeta$ standard normal. Since we work with z-values, the true $k$ is an even mixture of standard normals and thus itself standard normal: $k(z) = \phi(z)$.

If $g$ is a point mass at $\mu^*$, then $f(z) = r(z|\mu^*, 1)$ (remember that $f = g\ast k$). The convolution estimator \eqref{eq:ftilde} equals $\tilde{f}(z) = r(z|\zeta \sqrt{\alpha_1} +\mu^*, 1)$, the derivative of its log-density equals $\zeta\sqrt{\alpha_1}+\mu^*-z$ with expected value $\mu^*-z$, proving unbiasedness. If $g$ is a normal distribution with mean $\mu^*$ and variance $\sigma^2_g$, then $f(z) = r(z|\mu^*, 1 + \sigma^2_g)$. The convolution estimator equals $\tilde{f}(z) = r(z|\zeta \sqrt{\alpha_1} +\mu^*, 1+ \sigma^2_g)$, the derivative of its log-density $\frac{\zeta\sqrt{\alpha_1}+\mu^*-z}{1+ \sigma^2_g}$ with expected value $\frac{\mu^*-z}{1+ \sigma^2_g}$, again proving unbiasedness.

# Empirical validation

## Empirical validation of empirical properties

In the main text, the theoretical results from Sections \ref{sec:tweedieBiased}-\ref{sec:solution} are validated using Monte-Carlo simulations in Section \ref{sec:Tweedieconv}, as described in Section \ref{sec:simparam}. Here we provide exhaustive results, including not only the compound symmetric correlation structure but also block correlation.

### Naive estimator

The bias of the naive estimator discussed in Section \ref{sec:tweedieBiased} in the main text is shown in \fref{fig:biasLogDeriv} in the main text, the MSE and variance are shown in \fref{fig:varianceLogDeriv}.

```{r tryoutTweedie}
ps = c(5e2, 1e3, 5e3)
Grid = expand.grid("Sd" = Sds, "p" = ps)
ciReps = 5e2;extr = 4
zEval = seq(-extr, extr, length.out = 1e4)
if(!file.exists(theorFile <- "simResults/theorFile.RData")){
    logDerivs = mclapply(seq_len(nrow(Grid)), mc.cores = nCores, function(i){
            p = Grid[i, "p"];Sd = Grid[i, "Sd"]
             if(!file.exists(booFile <- paste0("simResults/theorFiles/Sd", Sd, "p", p, ".RData"))){
                mat = rmvnorm(n = ciReps, mean = rep(0, p), sigma = makeSigma(Sd, p))
                out = vapply(seq_len(ciReps), FUN.VALUE = double(8), function(j){
                        estDens(z = mat[j,], densMethod = "Lindsey", zObs = zEval, returnFit = TRUE)
                })
               save(out, file = booFile)
        } else load(booFile)
        out
    })
    meanVarianceDerivs = simplify2array(mclapply(mc.cores = nCores, logDerivs, function(x) {
        estLogDeriv = apply(x, 2, evalFitLogDeriv, zEval)
        diff = estLogDeriv+zEval
        cbind("Bias" = rowMeans(diff, na.rm = TRUE), 
              "MSE" = rowMeans(diff^2, na.rm = TRUE),
              "Variance" = rowMeans((estLogDeriv-rowMeans(estLogDeriv))^2, na.rm = TRUE))
    }))
    save(logDerivs, meanVarianceDerivs, file = theorFile)
} else load(theorFile) 
```

```{r logderivGraphs, include = FALSE}
#Mean
moltMeans = melt(meanVarianceDerivs, varnames = c("z", "what", "j"), value.name = "value")
moltMeans$z = zEval[moltMeans$z]
moltMeans = cbind(moltMeans, Grid[moltMeans$j,])
moltMeans$what = factor(moltMeans$what, ordered = TRUE, labels = c("Bias", "Variance", "MSE"), levels = c("Bias", "Variance", "MSE"))
ggplot(data = moltMeans[moltMeans$what=="Bias" & abs(moltMeans$z) < 1,], aes(x = z, y = value, col = factor(p))) +
    geom_line() + facet_grid(Sd~., scales = "free_y") +
    scale_colour_discrete(name = "Number of features") +
    geom_hline(yintercept = 0, linetype = "solid", linewidth = 0.3) +
    geom_abline(aes(slope = -Sd, intercept = 0), linetype = "dashed", linewidth = 0.3) +
    geom_line(aes(x = z, y = -Sd*z-Sd^2*z*(z^2-1)), linetype = "dotted", linewidth = 0.3, inherit.aes = FALSE) +
    ylab("Bias")
ggsave("ManuscriptSelBias/Graphs/LogDerivMean.pdf", height = 6.5, width = 8) 
rm(meanVarianceDerivs)
```

```{r logderivVar, fig.cap = "Variance and MSE of the naive estimator for the derivative of the log density based on the observed z-values (y-axis) as a function of z-value (x-axis), correlation $\\rho$ between features (rows) and number of features $p$ (colour) \\textbf{under compound symmetry}. The horizontal dashed line indicates $\\alpha_1 = \\rho$, the average correlation, whereas the dotted line indicates $\\alpha_1+2z^2\\alpha_2$: the first order and second order approximations to the variance as in \\eqref{eq:varResult} respectively. p=500 is omitted for legibility as it leads to very large variances.\\label{fig:varianceLogDeriv}"}
#Variance
ggplot(data = moltMeans[moltMeans$what %in% c("Variance", "MSE") & abs(moltMeans$z) < 1 & moltMeans$p >= 1e3,], 
       aes(x = z, y = value, col = factor(p))) +
    geom_line() +facet_grid(Sd~what, scales = "free_y") +
    geom_hline(aes(yintercept = Sd), linetype = "dashed", linewidth = 0.3) +
    geom_line(aes(x = z, y = Sd+Sd^2*z^2*2), linetype = "dotted", linewidth = 0.3, inherit.aes = FALSE) +
    scale_colour_discrete(name = "Number of features") +
    ylab("Variance or MSE")
```

\clearpage
 
### Convolution estimator

Exhaustive results for the convolution estimator from Section \ref{sec:solution} in the main paper are provided here.

#### Compound symmetry

The bias of the three estimators (naive, bagging and convolution) are shown in \fref{fig:biasLogDeriv2} in the main text. The more subtle difference between the convolution and bagging estimators is shown in \fref{fig:biasBagged} here, the variance and MSE are depicted in Figures \ref{fig:varLogDeriv2}-\ref{fig:mseLogDeriv2}.

```{r TweedieWithConvolution}
nTheor = 10;B = 20
ciReps = 5e2; repsG = 20
zEval = seq(-extr, extr, length.out = 1e3)
minAlpha = 0.05
if(!file.exists(theorFile2 <- "simResults/theorFile2.RData")){
    dir.create("simResults/theorFiles2")
    logDerivs2 = mclapply(seq_len(nrow(Grid)), mc.cores = nCores, mc.preschedule = FALSE, function(i){
            p = Grid[i, "p"];Sd = Grid[i, "Sd"]
            Vars = runif(p, varRange[1], varRange[2]);sds = sqrt(Vars)
            trueDensDerivLog = rowMeans(vapply(integer(repsG), FUN.VALUE = zEval, function(i){
                    mu = makeMean(p, sd = sdMean, mode = "dense")
                    vapply(zEval, FUN.VALUE = double(1), function(z){
                        anaDerivLog(z, mu, sd = sqrt(Vars/n))["deriv1"]
                    })
            }))
            if(!file.exists(booFile <- paste0("simResults/theorFiles2/Sd", Sd, "p", p, ".RData"))){
                mat = rmvnorm(n = ciReps*nTheor, mean = rep(0, p), sigma = makeSigma(Sd, p)*outer(sds, sds))
                out = vapply(seq_len(ciReps), FUN.VALUE = matrix(0, length(zEval), 3), function(j){
                         mu = makeMean(p, sd = sdMean, mode = "dense")
                        matIn = mat[seq_len(nTheor) + (j-1)*nTheor,] + rep(mu, nTheor)
                        naiveCoef = estDens(z = mv <- colMeans(matIn), densMethod = "Lindsey", returnFit = TRUE)
                        naiveEval = evalFitLogDeriv(naiveCoef, z = zEval)
                        bootMeans = vapply(integer(B), FUN.VALUE = mv, function(i){
                            colMeans(matIn[sample(n, replace = TRUE),])
                        })
                        bootCoef = estDens(z = c(bootMeans), densMethod = "Lindsey", returnFit = TRUE)
                        bootEval = evalFitLogDeriv(bootCoef, z = zEval)
                        alpha1 = getAlpha1(matIn)  #unlikely to be zero due to non-negative definiteness
                         SQR = sqrt(alpha1)
                         sdevs = apply(matIn, 2, sd)/sqrt(n)
                        a5eval = if(alpha1 < minAlpha){
                                naiveEval
                            } else {
                                vapply(zEval, FUN.VALUE = 0, function(z) anaDerivLog(z, mv,sdevs*SQR)["deriv1"])
                            }
                        mateval = cbind("naive" = naiveEval, "a5" = a5eval, "bagged" = bootEval)
                })
                diff = out - trueDensDerivLog
                bias = rowMeans(diff, dims = 2)
                mse = rowMeans(diff^2, dims = 2)
                variance = rowMeans((out - rowMeans(out))^2, dims = 2)
               save(bias, variance, mse, file = booFile)
        } else load(booFile)
        list("bias" = bias, "variance" = variance, "mse" = mse)
    })
    save(logDerivs2, file = theorFile2)
} else load(theorFile2)
```

```{r logderiv2Graphs, include = FALSE}
colValues = c("Naive" =  "#F8766D",  "Convolution" = "#00BA38", "Bagging" = "#619CFF")
#Mean
molt2 = melt(logDerivs2, varnames = c("z", "estimator"), value.name = "value")
names(molt2)[4:5] = c("what", "j")
molt2$z = zEval[molt2$z]
molt2 = cbind(molt2, Grid[molt2$j,])[molt2$estimator %in% c("naive", "a5", "bagged"),]
molt2$estimator = factor(molt2$estimator, ordered = TRUE, levels = c("naive", "a5", "a5smoothed", "bagged"), labels = c("Naive", "Convolution", "Convoluted smoothed", "Bagging"))
ggplot(data = molt2[molt2$what=="bias" & abs(molt2$z) < 4,], aes(x = z, y = value, col = factor(estimator))) +
    geom_line() + facet_grid(Sd~p, scales = "free_y") +
    scale_colour_manual(values = colValues, name = "Estimator") +
    geom_hline(yintercept = 0, linetype = "dotted", linewidth = 0.3) +
    ylab("Bias") + xlab(expression(gamma))
ggsave("ManuscriptSelBias/Graphs/LogDerivMean2.pdf", height = 6.5, width = 8)
```

```{r biasBag, fig.cap = "Bias in the estimation of the derivative of the log density of $f$ (y-axis) as a function of estimand $\\gamma$ (x-axis), correlation between features (rows), number of features $p$ (column) and estimator (colour) \\textbf{under compound symmetry}. Only the convolution and bagging estimators are shown (see \\fref{fig:biasLogDeriv2} in the main text for the naive estimator). The horizontal dotted line indicates 0 (no bias). This figures illustrates how bagging remedies the convolution estimator's small sample bias (low \\textit{p}) in the tails at zero correlation, but also that the bagging estimator is itself biased at high-correlation, no matter what the value of \\textit{p}. \\label{fig:biasBagged}"}
ggplot(data = molt2[molt2$what=="bias" & abs(molt2$z) < 4 & molt2$estimator %in% c("Convolution", "Bagging"),], aes(x = z, y = value, col = factor(estimator))) +
    geom_line() + facet_grid(Sd~p, scales = "free_y") +
    scale_colour_manual(values = colValues, name = "Estimator") +
    geom_hline(yintercept = 0, linetype = "dotted", linewidth = 0.3) +
    ylab("Bias") + xlab(expression(gamma))
```

```{r logderiv2GraphsVar, fig.cap = "Variance of the estimation of the derivative of the log density (y-axis) as a function of $\\gamma$ (x-axis), correlation between features (rows), number of features $p$ (column) and estimator (colour) \\textbf{under compound symmetry}. p=500 is omitted for legibility as it leads to very large variances; the y-axis is on the log-scale. At a correlation of 0, the naive and convolution estimators are identical and their lines overlap.\\label{fig:varLogDeriv2}"}
#Variance
ggplot(data = molt2[molt2$what=="variance" & abs(molt2$z) < 4 & molt2$p >= 1e3,], aes(x = z, y = value, col = factor(estimator))) +
    geom_line() + facet_grid(Sd~p, scales = "free_y") +
    scale_colour_manual(values = colValues, name = "Estimator") +
    scale_y_log10()+
    ylab("Variance")+ xlab(expression(gamma))
```

```{r logderiv2GraphsMSE, fig.cap = "MSE of the estimation of the derivative of the log density (y-axis) as a function of $\\gamma$ (x-axis), correlation between features (rows), number of features $p$ (column) and estimator (colour) \\textbf{under compound symmetry}. p=500 is omitted for legibility as it leads to very large MSEs; the y-axis is on the log-scale. At a correlation of 0, the naive and convolution estimators are identical and their lines overlap.\\label{fig:mseLogDeriv2}"}
ggplot(data = molt2[molt2$what=="mse" & abs(molt2$z) < 4 & molt2$p >= 1e3,], aes(x = z, y = value, col = factor(estimator))) +
    geom_line() + facet_grid(Sd~p, scales = "free_y") +
    scale_colour_manual(values = colValues, name = "Estimator") +
    scale_y_log10()+
    ylab("MSE")+ xlab(expression(gamma))
rm(molt2)
```

\clearpage

#### Block correlation \label{sec:block}

In a block correlation scenario, the number of features is fixed at p=2,000, and the correlation structure is a block correlation matrix with fixed correlation within blocks and no correlation between blocks of features. The block size is varied between 100, 500 and 1,000.

```{r blockCor}
blockSizes = c(100, 500, 1000)
p = 2e3
GridWeak = expand.grid("Sd" = Sds, "bs" = blockSizes)
zEval = seq(-extr, extr, length.out = 1e4)
if(!file.exists(theorFileWeak <- "simResults/theorFileWeak.RData")){
    dir.create("simResults/theorFilesWeak")
    logDerivsWeak = mclapply(seq_len(nrow(GridWeak)), mc.cores = nCores, mc.preschedule = FALSE, function(i){
            Sd = GridWeak[i, "Sd"];bs = GridWeak[i, "bs"]
            Vars = runif(p, varRange[1], varRange[2]);sds = sqrt(Vars)
            trueDensDerivLog = rowMeans(vapply(integer(repsG), FUN.VALUE = zEval, function(i){
                    mu = makeMean(p, sd = sdMean, mode = "dense")
                    vapply(zEval, FUN.VALUE = double(1), function(z){
                        anaDerivLog(z, mu, sd = sds/sqrt(n))["deriv1"]
                    })
            }))#Truly integrate over prior
            Sigma = bdiag(lapply(integer(p/bs), function(ii) makeSigma(Sd, bs)))
             if(!file.exists(booFile <- paste0("simResults/theorFilesWeak/bs", bs, "Sd", Sd, ".RData"))){
                mat = mvtnorm::rmvnorm(n = ciReps*n, mean = rep(0, p), sigma = as.matrix(Sigma*outer(sds, sds)))
                out = vapply(seq_len(ciReps), FUN.VALUE = matrix(0, length(zEval), 3), function(j){
                        mu = makeMean(p, sd = sdMean, mode = "dense")
                        matIn = mat[seq_len(n) + (j-1)*n,] + rep(mu, each = n)
                        naiveCoef = estDens(z = mv <- colMeans(matIn), densMethod = "Lindsey", returnFit = TRUE)
                        naiveEval = evalFitLogDeriv(naiveCoef, z = zEval)
                        bootMeans = vapply(integer(B), FUN.VALUE = mv, function(i){
                            colMeans(matIn[sample(n, replace = TRUE),])
                        })
                        bootCoef = estDens(z = c(bootMeans), densMethod = "Lindsey", returnFit = TRUE)
                        bootEval = evalFitLogDeriv(bootCoef, z = zEval)
                        alpha1 = getAlpha1(matIn) #unlikely to be zero due to non-negative definiteness
                        sdevs = apply(matIn, 2, sd)/sqrt(n)
                        a5eval = if(alpha1 < minAlpha){
                                naiveEval
                            } else {
                                vapply(zEval, FUN.VALUE = 0, function(z) anaDerivLog(z, mv,sdevs*sqrt(alpha1))["deriv1"])
                            }
                        mateval = cbind("naive" = naiveEval, "a5" = a5eval, "bagged"= bootEval)
                })
                diff = out - trueDensDerivLog
                bias = rowMeans(diff, dims = 2)
                mse = rowMeans(diff^2, dims = 2)
                variance = rowMeans((out - rowMeans(out))^2, dims = 2)
               save(bias, variance, mse, file = booFile)
        } else load(booFile)
        list("bias" = bias, "variance" = variance, "mse" = mse)
    })
    save(logDerivsWeak, file = theorFileWeak)
} else load(theorFileWeak)
```

```{r logderiv2GraphsBiasWeak, fig.cap = "Bias of the estimation of the derivative of the log density (y-axis) \\textbf{under block correlation} as a function of $\\gamma$ (x-axis), correlation between features (rows), correlation block size (column) and estimator (colour). At a correlation of 0, the naive and convolution estimators are identical and their lines overlap.\\label{fig:biasLogDerivWeak}"}
#Mean
moltMeansWeak = melt(logDerivsWeak, varnames = c("z", "estimator"), value.name = "value")
names(moltMeansWeak)[4:5] = c("what", "j")
moltMeansWeak$z = zEval[moltMeansWeak$z]
moltMeansWeak = moltMeansWeak[moltMeansWeak$estimator %in% c("naive", "a5", "bagged"),]
moltMeansWeak$estimator = factor(moltMeansWeak$estimator, ordered = TRUE, levels = c("naive", "a5", "a5smoothed", "bagged"), labels = c("Naive", "Convolution", "Convoluted smoothed", "Bagging"))
moltMeansWeak = cbind(moltMeansWeak, GridWeak[moltMeansWeak$j,])
ggplot(data = moltMeansWeak[moltMeansWeak$what=="bias" & abs(moltMeansWeak$z) < 4,], aes(x = z, y = value, col = factor(estimator))) +
    geom_line() + facet_grid(Sd~bs, scales = "free_y") +
    scale_colour_manual(values = colValues, name = "Estimator") +
    geom_hline(yintercept = 0, linetype = "solid", linewidth = 0.3) +
    ylab("Bias")
```

```{r logderiv2GraphsVarWeak, fig.cap = "Variance of the estimation of the derivative of the log density (y-axis) \\textbf{under block correlation} as a function of $\\gamma$ (x-axis), correlation between features (rows), correlation block size (column) and estimator (colour). The y-axis is on the log-scale. At a correlation of 0, the naive and convolution estimators are identical and their lines overlap.\\label{fig:varLogDerivWeak}"}
#Variance
ggplot(data = moltMeansWeak[moltMeansWeak$what=="variance" & abs(moltMeansWeak$z) < 4,], aes(x = z, y = value, col = factor(estimator))) +
    geom_line() + facet_grid(Sd~bs, scales = "free_y") +
    scale_colour_manual(values = colValues, name = "Estimator") +
    ylab("Variance") + scale_y_log10()
rm(moltMeansWeak, moltMeans)
```

\clearpage

## Empirical comparison of correction methods: parametric simulations

### Sources of correlation between estimates \label{sec:originCor}

\hl{We investigate causes of correlation between estimates in the following simulation study. Matrices of regressors $\mb X$ are generated as in the dense scenario of the parametric simulation in Section \ref{sec:simparam} in the main text, with different correlation strengths $\rho \in [0, 0.25, 0.5, 0.75]$. Next a coefficient vector $\bs\beta$ of length $p$ is generated through $p$ independent draws from a normal distribution with mean 0 and variance either 0, 0.04, 0.25 or 0.49, and the outcome vector $\mb y$ by independent draws from a normal distribution with variance 1 and means $\mb X\bs\beta$. Next $\mb y$ is regressed on every column of $\mb X$ using maximum likelihood, to estimate the slope $\beta_{yx}$. Conversely, every column of $\mb X$ is regressed on $\mb y$ to estimate the slope $\beta_{xy}$. In addition, the out-of-sample MSE of all regression models is estimated through cross-validation, leading to sets of estimates $\widehat{MSE}_{yx}$ for predicting y from x and $\widehat{MSE}_{xy}$ for predicting x from y. 500 Monte-Carlo instances are generated per correlation strength $\rho$, and all pairwise correlations between the $p$ estimates over these simulation instances are calculated and depicted in \fref{fig:corOrigin}. This reveals that the out-of-sample MSE is always correlated when estimated on the same outcome variable $y$ ($\widehat{MSE}_{yx}$). Yet the MSE with a common regressor $y$ for all prediction models ($\widehat{MSE}_{xy}$), and the slopes of regressing y on all columns of $\mb X$ ($\hat{\beta}_{yx}$), or all columns of $\mb X$ on y ($\hat{\beta}_{xy}$), are only correlated when the features in $\mb X$ are correlated. The correlations between $\hat{\beta}_{yx}$ and $\hat{\beta}_{xy}$ grow faster than the correlations between the regressors when there is a real relationship between the outcome and regressors, such that even moderately correlated regressors lead to considerably correlated estimators.}

```{r boxplotssim, fig.height = 8, fig.cap = "\\hl{Boxplots of pairwise correlations between estimators approximated by Monte-Carlo simulation (y-axis) as a function of the correlation between features (x-axis), for different estimated quantities (rows) and different signal strengths (columns). The horizontal dotted line indicates zero correlation, the diagonal dotted blue line has slope 1 and intercept 0.\\label{fig:corOrigin}}"}
p = 1e3;sdMeans = c(0, 0.2, 0.5, 0.7);names(sdMeans) = sdMeans
if(!file.exists(simFileCors <- "simResults/simCors.RData")){ 
    simCors = lapply(Sds, function(Sd){
        lapply(sdMeans, function(sdMean){  cat(sdMean, "\t")
            mclapply(integer(ciReps), mc.cores = nCores, function(i){
                x = rmvnorm(n = n, mean = rep(0, p), sigma = makeSigma(Sd, p))
                betas = rnorm(p, sd = sdMean)
               dat = list("x" = x, "y" = rnorm(n, mean = x %*% betas))
               Coefs = vapply(seq_len(p), FUN.VALUE = double(2), function(j){
                        coef1 = lm.fit(dat$x[, j], x =  cbind(1, dat$y))$coef[2]
                        coef2 = lm.fit(dat$y, x =  cbind(1, dat$x[, j]))$coef[2]
                        c(coef1, coef2)
               })
                MSEests1 = vapply(simpleCV(dat = dat, nOuterFolds = nFolds), FUN.VALUE = double(1), function(x) mean(unlist(x)))
                MSEests2 = {
                     folds = sample(rep(unFolds <- seq_len(nFolds), length.out = nrow(dat$x)))
                    vapply(seq_len(ncol(dat$x)), FUN.VALUE = 0, function(j){
                        mean(unlist(sapply(unFolds, function(uf){
                            idTrain = folds!=uf
                            predTest = predLin(dat$y[idTrain], dat$x[idTrain, j],
                                               dat$y[!idTrain])
                            (predTest-dat$x[!idTrain, j])^2
                        })))
                    })
                }
                out = rbind(Coefs, MSEests1, MSEests2)
                rownames(out) = c("EstimateXonY", "EstimateYonx", "MSEYonX", "MSEXonY")
                out
            })
        })
    })
    save(simCors, file = simFileCors)
} else load(simFileCors)
foo = lapply(simCors, function(xx){
    lapply(xx, function(yy){
    tmp = lapply(nm <- rownames(yy[[1]]), function(x){
        tmp = t(vapply(yy, FUN.VALUE = double(p), function(y) y[x,]))
        cm = cor(tmp, use = "pairwise.complete.obs")
        cm[upper.tri(cm)]
    })
    names(tmp) = nm;tmp
    })
})
foo2 = melt(foo); names(foo2) = c("Correlation", "Measure", "SignalStrength", "Sd")
foo2$Measure = factor(foo2$Measure, ordered = TRUE, levels =  c("EstimateYonx", "EstimateXonY", "MSEYonX", "MSEXonY"), labels = c("EstimateYonx", "EstimateXonY", "MSEYonX", "MSEXonY"))
facet_labels = c("EstimateXonY" = "beta[xy]", "EstimateYonx" = "beta[yx]", "MSEYonX" = "MSE[yx]", "MSEXonY" = "MSE[xy]")
foo2$SignalStrength = as.numeric(foo2$SignalStrength)^2
ggplot(data = foo2, aes(y = Correlation, x = Sd)) + geom_boxplot() + 
    facet_grid(Measure~SignalStrength, labeller = labeller(Measure = as_labeller(facet_labels, label_parsed))) +
    xlab("Correlation between features in X") + geom_hline(yintercept = 0, linetype = "dotted") + 
    ylab("Pairwise correlations between estimators") +
    geom_abline(slope = 0.25, intercept = -0.25, linetype = "dashed", colour = "blue")
rm(simCors, foo, foo2)
```

\clearpage

### Convolution repairs empirical Bayes methods

This section zooms in on the problems of empirical Bayes methods Tweedie's formula, ash and VanZwet2021 under dependence and how then can be repaired by convolution. The parametric simulation results in the dense scenario are shown in \fref{fig:varScenEmpBayes} in the main text, for the sparse scenario they are shown in \fref{fig:varScenBag}.

```{r baseSimCI}
Sigmas = lapply(Sds, function(Sd){
    Sigma = matrix(Sd, p, p)
    diag(Sigma) = 1
    Sigma
  })
varRange = c(2,10)
ciReps = 5e2
effSize = -1; effFrac = 0.1
modes = c("dense", "sparse");names(modes) = modes
if(!file.exists(baseSimCIVarFile <- "simResults/baseSimVarCI.RData")){
    baseTestVarCI = lapply(modes, function(mode){
        lapply(Sds, function(Sd){
            mclapply(seq_len(ciReps), mc.cores = nCores, function(j){
                    meanVec = makeMean(p, sd = sdMean, effFrac = effFrac, effSize = effSize, mode = mode)
                    names(meanVec) = seq_len(p)
                    VarsTrue = runif(p, varRange[1], varRange[2])
                    Sig = Sigmas[[as.character(Sd)]]*tcrossprod(sqrt(VarsTrue))
                    mat = rmvnorm(n = n, mean = meanVec, sigma = Sig)
                    rownames(mat) = seq_len(n)
                    getSimpleEstsMat(mat = mat, bootReps = bootReps, meanVec = meanVec, 
                                     extremes = extVec, VarsTrue = VarsTrue/n, ciReturn = TRUE)
            })
        })
    })
    save(baseTestVarCI, file = baseSimCIVarFile)
} else load(baseSimCIVarFile) 
plotNormal = c("rawEst", "FayeCorrectEst", "TanCorrectEst","TanCorrectEstParam",
               "tweedieConvEsts", "split", "condML", "ash",
               "ashConv", "catScores", "bootCorrectForde", "VanZwet2021" , "VanZwet2021conv")
plotNormalPaper = c("rawEst", "FayeCorrectEst", "TanCorrectEst", "VanZwet2021conv",
               "tweedieConvEsts", "split", "condML", "ashConv", "catScores", "bootCorrectForde")
```

```{r depBoot2, include  = FALSE}
depBootObj = lapply(Sds[c(1, 4)], function(Sd){
    lapply(seq_along(baseTestVarCI$dense[[as.character(Sd)]]), function(j){
            mat = baseTestVarCI$dense[[as.character(Sd)]][[j]]$mat #Get the matrices here
            alpha1 = getAlpha1(mat)
            list("cMeans" = colMeans(mat), "alpha1" = alpha1, "cVars" = apply(mat,2, var)/n)
    })
})
pdf("ManuscriptSelBias/Graphs/depBoot2.pdf", height = 8, width = 7) 
nBreaks = 25
xLims = c(-2.5, 2.5); nBreaks = 30; cexMain = 1.2; cexLab = 1.1
meanLty = "dashed";densLty = "solid"
par(mfrow = c(3, 2), mar = c(2, 4, 1.9, 0.5))
trueMarRMSE = density(unlist(lapply(depBootObj[["0"]], function(x) x$cMeans)))
zSeq = seq(xLims[1], xLims[2], length.out = 1e3)
for(i in seq_len(3)){
    if(i==3){
        par(mar = c(4.25, 4.2, 2, 0.5))
    }
    hist(cMeans <- depBootObj[["0"]][[i]]$cMeans, freq = FALSE, xlim = xLims, breaks = nBreaks, 
         xlab = if(i==3) "Estimate" else "", 
           main = if(i == 1) "Independent estimates", cex.main = cexMain, cex.lab = cexLab)
    abline(v = 0, col = "red", lty = meanLty);lines(trueMarRMSE, col = "blue", lty = densLty)
    lines(density(cMeans), col = "black", lty = densLty)
    alpha1 = depBootObj[["0"]][[i]]$alpha1
    evalMix = if(alpha1 > minAlpha){
        evalMixNorm(zSeq, mu = cMeans, sd = sqrt(depBootObj[["0"]][[i]]$cVars*alpha1))
    } else {
     predict(kde(cMeans), x = zSeq)
    }
    lines(zSeq, evalMix, col = "darkgreen", lty = densLty)
    hist(cMeansDep <- depBootObj[["0.75"]][[i]]$cMeans, freq = FALSE, xlim = xLims, cex.lab = cexLab, breaks = nBreaks, 
           xlab = if(i==3) "Estimate" else "", main = if(i == 1) "Dependent estimates", cex.main = cexMain); abline(v = 0, col = "red", lty = meanLty)
    alpha1 = depBootObj[["0.75"]][[i]]$alpha1
    lines(trueMarRMSE, col = "blue", lty = densLty)
    lines(density(cMeansDep), col = "black", lty = densLty)
    evalMix = if(alpha1 > minAlpha){
        evalMixNorm(zSeq, mu = cMeansDep, sd = sqrt(depBootObj[["0.75"]][[i]]$cVars*alpha1))
    } else {
     predict(kde(cMeansDep), x = cMeansDep)
    }
    lines(zSeq, evalMix, col = "darkgreen", lty = densLty)
}
par(mfrow = c(1,1), mar = c(4,4,4,4))
dev.off()
rm(depBootObj)
```

```{r onlyEmpBayes}
plotEmp = c("rawEst", "tweedieEst", "tweedieConvEsts", "ash", "ashConv", "VanZwet2021", "VanZwet2021conv")
if(!file.exists(empFilePaper <- "Results/empDfPaper.RData")){
    p1 = plotSimpleRes(baseTestVarCI, extremes = extVec, p = p, Methods2Plot = plotEmp, returnDf = TRUE)
    p2 = plotSimpleRes(baseTestVarCI, extremes = extVec, p = p, power = 2, Methods2Plot = plotEmp, returnDf = TRUE)
    p4 = plotRunningTDP(baseTestVarCI, extremes = extVec, p = p, Methods2Plot = plotEmp, returnDf = TRUE) 
    p5 = plotCoverage(baseTestVarCI, extremes = extVec, Methods2Plot = plotEmp, returnDf = TRUE)
    p6 = plotWidthCI(baseTestVarCI, extremes = extVec, Methods2Plot = plotEmp, returnDf = TRUE)
    empDf = rbind(p1, p2, p4, p5, p6)
    empDf$Method = factor(empDf$Method, ordered = TRUE, labels = methodLabels, levels = methodLevels)
    empDf = empDf[with(empDf, Performance_measure == "TDR" | what == "Raw estimate"),]
    empDf = empDf[with(empDf, !(grepl("Ash", Method) & what == "Test statistic")),]
    empDf$Performance_measure = factor(empDf$Performance_measure, ordered = TRUE, levels = c("Bias", "MSE", "TDR", "Coverage", "Width"))
    save(empDf, file = empFilePaper)
} else load(empFilePaper)
```

```{r bagPlot, fig.cap = "Simulation results for mean estimation in parametric simulation in dense scenario for different performance measures (side panels) for the raw estimates and naive and convolution empirical Bayes methods (colour) for increasing correlation between features (top panels). The x-axis shows the number of features with smallest estimates considered. The dotted horizontal line indicates zero bias or MSE or the nominal coverage of 95\\%, respectively. VanZwet2021 is natively resistent to dependence. Both Tweedie's formula's and ash's performance is deteriorated by dependence, but the inclusion of convolution improves their feature ranking, restores reasonable coverage for ash's confidence intervals and improves selection bias correction for both methods. For zero correlation, the convolution corrected versions mostly overlap with the naive versions. All measures shown are averages over 500 Monte-Carlo instances.\\label{fig:varScenEmpBayes}", fig.height = 8.5, fig.width = 10, include = FALSE}
Theme = theme(legend.position = "top") 
lineSize = 0.3
lineDf = data.frame("Performance_measure" = c("Bias", "MSE", "Coverage"), "value" = c(0, 0, 1-sigLevel))
lineDf$Performance_measure = factor(lineDf$Performance_measure, ordered = TRUE, levels = c("Bias", "MSE", "TDR", "Coverage", "Width"))
plotParEmpBayes = ggplot(data = empDf[(empDf$Method %in% methodLabels[match(table = methodLevels, plotEmp)]) & (empDf$what == "Raw estimate") & empDf$mode == "dense" ,], aes(y = value, colour = Method, x = Top_features)) + geom_line() +
    facet_grid(Performance_measure ~ Cor, scales = "free_y") +
    ylab("Performance measure") + xlab("Number of smallest estimates") +
    guides(colour = guide_legend(nrow = 1)) +
        scale_colour_manual(values = colourVec[names(colourVec) %in% levels(empDf$Method)]) +
        Theme + geom_hline(data = lineDf, aes(yintercept = value), linetype = "dotted")
pdf(paste0("ManuscriptSelBias/Graphs/VarScen50empBayes.pdf"), height = 10, width = 9.5)
print(plotParEmpBayes + geom_hline(data = lineDf, aes(yintercept = value), linetype = "dotted"))
dev.off();rm(plotParEmpBayes)
```

```{r bagPlot2, fig.cap = "\\hl{Simulation results for mean estimation in parametric simulation in the sparse scenario for different performance measures (side panels) for the raw estimates and naive and convolution empirical Bayes methods (colour) for increasing correlation between features (top panels). The x-axis shows the number of features with smallest estimates considered. The dotted horizontal line indicates zero bias or MSE or the nominal coverage of 95\\%, respectively. For zero correlation, the convolution corrected versions sometimes overlap with the naive versions.}\\label{fig:varScenBag}", fig.height = 8.5, fig.width = 10}
ggplot(data = empDf[(empDf$Method %in% methodLabels[match(table = methodLevels, plotEmp)]) & (empDf$what == "Raw estimate") & empDf$mode == "sparse",], 
       aes(y = value, colour = Method, x = Top_features)) + geom_line() +
  facet_grid(Performance_measure ~ Cor, scales = "free_y") +
        ylab("Performance measure") + xlab("Number of smallest estimates") +
    guides(colour = guide_legend(nrow = 2)) +
        scale_colour_manual(values = colourVec[names(colourVec) %in% levels(empDf$Method)]) +
        Theme + geom_hline(data = lineDf, aes(yintercept = value), linetype = "dotted")
rm(empDf)
```

Apart from estimating the derivative of the log density for Tweedie's formula, we also investigate the effect of convolution on ash and VanZwet2021. \fref{fig:ashIllustr} shows how convolution improves the estimator for the prior distribution of the ash method under dependence in a parametric simulation study under the dense scenario. \fref{fig:zwetIllustr} illustrates the improvement for the estimator of the z-value distribution for the VanZwet2021 method. Furthermore, \fref{fig:ashSimVar} shows how the widths of the respective prior and z-value distributions better approximate the width of the true distribution after convolution.

```{r ashresam, fig.cap = "Prior distribution of the estimand $\\gamma$ estimated by ash with increasing correlation between the features (plot titles) as in the dense scenario of parametric simulation. 4 Monte-Carlo instances are shown per correlation. The naive estimates are shown in dark red, the convoluted estimates are shown in bright red; solid lines when the prior was modelled as a mixture of uniforms (the default throughout this study) and dashed lines when the prior was modelled as a mixture of normals. The true prior distribution, a Gaussian with mean 0 and variance 0.04 as known from the data generation mechanism, is shown in blue. In scenarios with correlation between features, convolution is seen to yield better estimates of the prior distribution. The y-axis is on the log-scale.\\label{fig:ashIllustr}", fig.height = 6.75}
numSamAsh = 10; 
if(!file.exists(depBootAshFile <- "simResults/depBootAshVanZwet.RData")){
    depBootAshVZ = lapply(Sds, function(Sd){
        mclapply(integer(ciReps), mc.cores = nCores, function(i){
            meanVec = makeMean(p, sd = sdMean, mode = "dense"); names(meanVec) = seq_len(p)
            VarsTrue = runif(p, varRange[1], varRange[2])
            Sig = Sigmas[[as.character(Sd)]]*tcrossprod(sqrt(VarsTrue))
            mat = rmvnorm(n = n, mean = meanVec, sigma = Sig)
            sigmahat = apply(mat, 2, sd)/sqrt(n)
            muhat = colMeans(mat); names(muhat) = seq_len(p)
            ashObj = ash(muhat, sigmahat, method = "shrink", mode = "estimate")
            #Now convolute
            alpha1 = getAlpha1(mat)
            ashObjConv = getConvAsh(ashObj = ashObj, alpha1 = alpha1, raws = muhat, ses = sigmahat, numSamAsh = numSamAsh)
            ashObjNormal = ash(muhat, sigmahat, method = "shrink", mode = "estimate", mixcompdist = "normal")
            ashObjConvNorm = getConvAsh(ashObj = ashObjNormal, alpha1 = alpha1, raws = muhat, 
                                        ses = sigmahat, numSamAsh = numSamAsh, mixcompdist = "normal")
            zwetObj = fitMix(muhat/sigmahat)
            zwetObjConv = if(alpha1 < 0){
                zwetObj
            } else {
                zEsts = rnorm(p*numSamAsh, muhat, sigmahat*sqrt(alpha1))
                fitMix(zEsts/sigmahat)
            }
            list("zwetObj" = zwetObj, "zwetObjConv" = zwetObjConv,
                 "ashObj" = ashObj, "ashObjConv" = ashObjConv, "ashObjNormal" = ashObjNormal, 
                 "ashObjConvNorm" = ashObjConvNorm, "meanVec" = meanVec)
        })
    })
    save(depBootAshVZ, file = depBootAshFile)
} else load(depBootAshFile)
lengthDens = 3e2; minDens = 1e-5
xLim = c(-1, 1)*2
Seq <- seq(xLim[1], xLim[2], length.out = lengthDens)
par(mfrow = c(4,4), mar = c(4, 4, 2, 0.5))
for(Sd in Sds){
    for(i in 1:4){
        with(depBootAshVZ[[as.character(Sd)]][[i]], {
            dens = get_density(ashObj, Seq)
            dens$y[dens$y < minDens] = minDens
            densBoot = get_density(ashObjConv, Seq)
            densBoot$y[densBoot$y < minDens] = minDens
            plot(dens, type = "l", ylab = "Density", xlab = bquote(gamma), main = paste("Cor:", Sd), 
             xlim = xLim, log = "y", ylim = c(1e-5, max(dens$y)), col = colourVec["Ash"])
            lines(densBoot, col = colourVec["Ash convoluted"])
            lines(densBoot$x, dnorm(densBoot$x, sd = sdMean), col = "blue") #True prior
            densNorm = get_density(ashObjNormal, Seq)
            densNorm$y[densNorm$y < minDens] = minDens
            densBootNorm = get_density(ashObjConvNorm, Seq)
            densBootNorm$y[densBootNorm$y < minDens] = minDens
            lines(densNorm, col = colourVec["Ash"], lty = "dashed")
            lines(densBootNorm, col = colourVec["Ash convoluted"], lty = "dashed")
        })
    }
}
par(mfrow = c(1,1))
```

```{r plotZwetSNR, fig.cap = "Distribution of the z-values estimated by VanZwet2021 under increasing correlation between the features (plot titles) as in the paramtric simulation study on mean estimation. 4 Monte-Carlo instances are shown per correlation. The naive estimates are shown in teal, the convoluted estimates are shown in cyan. The true distribution of the z-values, as known from the data generation mechanism, is shown in blue. In scenarios with correlation between features, convolution is seen to yield better estimates of the prior distribution.\\label{fig:zwetIllustr}", fig.height = 6.5}
xLim = c(-1, 1)*4
Seq <- seq(xLim[1], xLim[2], length.out = lengthDens)
trueZdistr = rowMeans(vapply(seq(varRange[1], varRange[2], length.out = lengthDens), FUN.VALUE = double(lengthDens), function(vv){
    dnorm(Seq, mean = 0, sd = sqrt(sdMean^2 + vv/n)/sqrt(vv/n))
}))
par(mfrow = c(4,4), mar = c(4, 4, 2, 0.5))
densList = lapply(names(depBootAshVZ), function(x){
    lapply(depBootAshVZ[[x]][sample(length(depBootAshVZ[[x]]), 4)], function(y){
        zwetObj = evalZwetDens(y$zwetObj, z = Seq)
        zwetObjConv = evalZwetDens(y$zwetObjConv, z = Seq)
        plot(Seq, trueZdistr/sum(trueZdistr), col = "blue", type = "l", main = x, 
             xlab = "z", log = "", xlim = xLim, ylab = "density", ylim = c(0, max(zwetObj/sum(zwetObj))))
        lines(Seq, zwetObj/sum(zwetObj), col = colourVec["VanZwet2021"], lty = "dashed")
        lines(Seq, zwetObjConv/sum(zwetObjConv), col = colourVec["VanZwet2021 convoluted"], lty = "dashed")
    })
})
trueVar = sdMean^2/(mean(varRange)/n) + (mean(varRange)/(n-1))*(n-1)/(n-3) + 1 #Variance of the means + t-distribution variance + 1 for z-statistics
```

```{r estZwetVars, fig.cap = paste0("\\textbf{Left:} Average variance of the estimated prior distribution of ash (y-axis) as a function of correlation strength (x-axis) for the naive and convolution ash estimator, in dark and bright red respectively. The vertical dashed line indicates the variance of the true prior (", sdMean^2, "). \\textbf{Right:} Average variance of the estimated z-value distribution (y-axis) as a function of correlation strength (x-axis) for naive and convolution VanZwet2021 estimator, in teal and cyan respectively. The vertical dashed line indicates the variance of the true z-value distribution (", signif(trueVar, 3) ,"). At a correlation of 0, the estimated variances without and with convolution coincide for both methods. The simulation settings are the same as in the parametric simulation study in the 'dense' scenario. \\label{fig:ashSimVar}"), fig.width = 6, fig.height = 3.5}
avAsh = colMeans(sapply(depBootAshVZ, function(x) sapply(x, function(y) getVarianceUniform(y[["ashObj"]]))))
avConv = colMeans(sapply(depBootAshVZ, function(x) sapply(x, function(y) getVarianceUniform(y[["ashObjConv"]]))))
avZwet = colMeans(sapply(depBootAshVZ, function(x) sapply(x, function(y) zwetOverallVar(y[["zwetObj"]]))))
avZwetConv = colMeans(sapply(depBootAshVZ, function(x) sapply(x, function(y) zwetOverallVar(y[["zwetObjConv"]]))))
par(mfrow = c(1,2), mar = c(4, 4, 3, 3))
plot(Sds, avAsh, col = colourVec["Ash"], ylab = "Average estimated variance", xlab = "Correlation between features", ylim = range(c(avAsh, avConv)), main = "ash", xaxt='n')
axis(side = 1, at = Sds, labels = TRUE, tcl = -0.2)
points(Sds, avConv, col = colourVec["Ash convoluted"])
abline(h = sdMean^2, lty = "dashed")
plot(Sds, main = "VanZwet2021", avZwet, col = colourVec["VanZwet2021"], ylab = "Average estimated variance", xlab = "Correlation between features", ylim = range(c(avZwet, avZwetConv)))
points(Sds, avZwetConv, col = colourVec["VanZwet2021 convoluted"])

abline(h = trueVar, lty = "dashed")
par(mfrow = c(1,1))
rm(depBootAshVZ)
```

\clearpage

### Exhaustive simulation results

```{r varPaper0, include = FALSE}
if(!file.exists(fullFilePaper <- "Results/fullDfPaper.RData")){
    p1 = plotSimpleRes(baseTestVarCI, extremes = extVec, p = p, Methods2Plot = plotNormal, returnDf = TRUE)
    p2 = plotSimpleRes(baseTestVarCI, extremes = extVec, p = p, power = 2, Methods2Plot = plotNormal, returnDf = TRUE)
    p4 = plotRunningTDP(baseTestVarCI, extremes = extVec, p = p, Methods2Plot = plotNormal, returnDf = TRUE) 
    p5 = plotCoverage(baseTestVarCI, extremes = extVec, Methods2Plot = c(plotNormal, "estOutSplit"), returnDf = TRUE)
    p6 = plotWidthCI(baseTestVarCI, extremes = extVec, Methods2Plot = c(plotNormal, "estOutSplit"), returnDf = TRUE)
    fullDf = rbind(p1, p2, p4, p5, p6)
    fullDf$Method = factor(fullDf$Method, ordered = TRUE, labels = methodLabels, levels = methodLevels)
    fullDf = fullDf[with(fullDf, Performance_measure == "TDR" | what == "Raw estimate"),]
    fullDf = fullDf[with(fullDf, !(grepl("Ash", Method) & what == "Test statistic")),]
    fullDf$Performance_measure = factor(fullDf$Performance_measure, ordered = TRUE, levels = c("Bias", "MSE", "TDR", "Coverage", "Width"))
    save(fullDf, file = fullFilePaper)
} else load(fullFilePaper)
linValues = c("Raw estimate" = "solid", "Test statistic" = "dashed", "Rescaled test statistic" = "dotdash")
plotPar = ggplot(data = fullDf[with(fullDf, mode == "dense" & Method %in% methodLabels[match(table = methodLevels, plotNormalPaper)] & what %in% c("Raw estimate", "Rescaled test statistic")),], aes(y = value, colour = Method, x = Top_features)) +
        geom_line(linewidth = lineSize) + facet_grid(Performance_measure ~ Cor, scales = "free_y") +
        ylab("Performance measure") + xlab("Number of smallest estimates") +
        guides(colour = guide_legend(nrow = 2)) +
        scale_colour_manual(values = colourVec[names(colourVec) %in% levels(fullDf$Method)]) +
        Theme + geom_hline(data = lineDf, aes(yintercept = value), linetype = "dotted")
pdf(paste0("ManuscriptSelBias/Graphs/VarScen50.pdf"), height = 10.5, width = 9)
print(plotPar)
dev.off();rm(plotPar)
```

```{r plotSparse, fig.cap = "\\label{fig:varScenSparse}Parametric simulation results in the sparse scenario for different performance measures (side panels) and methods (colour) for increasing correlation between features (top panels). The x-axis shows the number of features with smallest estimates considered. For the true discovery rate (TDR), the feature ranking is the same for conditional likelihood as for the raw estimates, so in this case only the raw estimates' results are shown. The dotted horizontal line indicates zero bias or MSE or the nominal coverage of 95\\%.", fig.height = 11, fig.width = 10}
plotParSparse = ggplot(data = fullDf[with(fullDf,mode == "sparse" & Method %in% methodLabels[match(table = methodLevels, plotNormalPaper)] & what == "Raw estimate"),], aes(y = value, colour = Method, x = Top_features)) +
        geom_line(linewidth = lineSize) + facet_grid(Performance_measure ~ Cor, scales = "free_y") +
        ylab("Performance measure") + xlab("Number of smallest estimates") +
 guides(colour = guide_legend(nrow = 3)) +
        scale_colour_manual(values = colourVec[names(colourVec) %in% levels(fullDf$Method)]) +
        Theme
print(plotParSparse + geom_hline(data = lineDf, aes(yintercept = value), linetype = "dotted"))
rm(fullDf, plotParSparse)
```

```{r mseOverall, fig.cap = "Boxplots of MSE of all estimates for parametric simulation study in the different scenarios (top panels) for different methods (colour) and increasing correlation between features (side panels) over 500 Monte-Carlo instances. The conditional likelihood method is not included as it does not calculate corrected estimates for all features.\\label{fig:mseAll}", fig.height = 6}
plotMSEres(baseTestVarCI, Methods2Plot = c(plotNormal, "estOutSplit"), colVecAdd = "Conditional likelihood") 
```

```{r varSup, fig.cap = "Full simulation results for parametric simulation study in the \\textbf{dense scenario} for different performance measures (side panels), different methods (colour) and increasing correlation between features (top panels). The x-axis shows the number of features with smallest estimates considered. The linetype indicates whether the result is based on raw estimates, test statistics or rescaled test statistics. The dotted horizontal line indicates zero bias or MSE or the nominal coverage of 95\\%. The feature ranking is the same for conditional likelihood as for the raw estimates, and when using test statistics also identical to ash with or without convolution, so in these cases only the results for the raw estimates are shown. Rescaling the corrected test statistics to the scale of the raw estimates does not systematically improve performance in terms of bias and MSE of top estimates with respect to correcting the raw estimates for any of the methods considered (separate estimation was omitted here as it is identical to the raw version). Nonparametric and parametric bootstrapping for Tan2015 yields virtually the same result in this simple case.\\label{fig:varScenSup}", fig.height = 12, fig.width = 10}
rescNames = paste0(setdiff(plotNormal, c("rawEst", "split")), "_rescale")
plotNormalResc = c(plotNormal, rescNames) 
plotNormalResc = plotNormalResc[!(plotNormalResc %in% c("catScores_rescale", "VanZwet2021_rescale", "VanZwet2021conv_rescale"))]
methodLevelsResc = c(methodLevels, rescNames)
methodLabelsResc = c(methodLabels, rescNames)
if(!file.exists(fullFileSup <- "Results/fullDfSup.RData")){
    baseTestVarCIresc = lapply(baseTestVarCI, function(xx){
        lapply(xx, function(x){
            lapply(x, function(y){
                #Raw estimates
                tmp = y$resMatZ[, setdiff(plotNormal, c("rawEst", "condML", "split", "VanZwet2021conv"))] * (ses <- sqrt(y$resMat[, "Vars"]))
                colnames(tmp) = paste0(colnames(tmp), "_rescale")
                y$resMat = cbind(y$resMat, tmp)
                ## condML
                condMLrescale = lapply(y$condMLests, function(xx){
                    lapply(xx, function(yy){
                        yy* ses[rownames(yy)]
                    })
                })
                #Confidence intervals
                ciListScale = lapply(y$ciListZ, function(x) x*ses)
                names(ciListScale) = paste0(names(ciListScale), "_rescale")
                y$ciList = c(y$ciList, ciListScale[colnames(tmp)])
                c(y, list("condML_rescale" = condMLrescale))
            })
        })
    })
    rm(baseTestVarCI)
    p1 = plotSimpleRes(baseTestVarCIresc, extremes = extVec, p = p, Methods2Plot = plotNormalResc, returnDf = TRUE, methodLevels = methodLevelsResc, methodLabels = methodLabelsResc)
    p2 = plotSimpleRes(baseTestVarCIresc, extremes = extVec, p = p, power = 2, Methods2Plot = plotNormalResc, returnDf = TRUE, methodLevels = methodLevelsResc, methodLabels = methodLabelsResc)
    p4 = plotRunningTDP(baseTestVarCIresc, extremes = extVec, p = p, Methods2Plot = plotNormalResc, returnDf = TRUE, methodLevels = methodLevelsResc, methodLabels = methodLabelsResc) 
    p5 = plotCoverage(baseTestVarCIresc, extremes = extVec, Methods2Plot = c(plotNormalResc, "estOutSplit"), returnDf = TRUE, methodLevels = methodLevelsResc, methodLabels = methodLabelsResc)
    p6 = plotWidthCI(baseTestVarCIresc, extremes = extVec, Methods2Plot = c(plotNormalResc, "estOutSplit"), returnDf = TRUE, methodLevels = methodLevelsResc, methodLabels = methodLabelsResc)
    fullDfSup = rbind(p1, p2, p4, p5, p6)
    fullDfSup = fullDfSup[with(fullDfSup, !(grepl("Ash", Method) & what == "Test statistic")),]
    save(fullDfSup, file = fullFileSup)
} else load(fullFileSup) 
fullDfSup2 = fullDfSup[with(fullDfSup, (Performance_measure == "TDR" | what == "Raw estimate") ),]
fullDfSup2$what[grep(fullDfSup2$Method, pattern = "rescale")] = "Rescaled test statistic"
fullDfSup2$what = factor(fullDfSup2$what, levels = names(linValues), labels = names(linValues), ordered = TRUE)
fullDfSup2$Method = factor(gsub(pattern = "_rescale", "", fullDfSup2$Method), levels = c(methodLevelsResc, methodLabelsResc), labels = c(methodLabelsResc, methodLabelsResc), ordered = TRUE)
fullDfSup2$Performance_measure = factor(fullDfSup2$Performance_measure, ordered = TRUE, levels = c("Bias", "MSE", "TDR", "Coverage", "Width"))
fullDfSup2b = fullDfSup2[!with(fullDfSup2, Performance_measure != "TDR" & Method == "Cat scores"),]
ggplot(data = fullDfSup2b[with(fullDfSup2, mode == "dense"),], aes(y = value, colour = Method, x = Top_features, linetype = what)) +
        geom_line(linewidth = lineSize) + facet_grid(Performance_measure ~ Cor, scales = "free_y") +
        ylab("Performance measure") + xlab("Number of smallest estimates") +
        scale_linetype_manual(name = "Estimate", values = linValues) + 
        guides(colour = guide_legend(nrow = 4), linetype = guide_legend(nrow = 3)) +
        scale_colour_manual(values = colourVec[names(colourVec) %in% levels(fullDfSup2$Method)]) +
        Theme + geom_hline(data = lineDf, aes(yintercept = value), linetype = "dotted")
```

```{r fullScenSparse, fig.cap = "\\hl{Full simulation results for parametric simulation study in the \\textbf{sparse scenario} for different performance measures (side panels), different methods (colour) and increasing correlation between features (top panels). The x-axis shows the number of features with smallest estimates considered. The linetype indicates whether the result is based on raw estimates, test statistics or rescaled test statistics. The dotted horizontal line indicates zero bias or MSE or the nominal coverage of 95\\%. The feature ranking is the same for conditional likelihood as for the raw estimates, and when using test statistics also identical to ash with or without convolution, so in these cases only the results for the raw estimates are shown. Rescaling the corrected test statistics to the scale of the raw estimates does not systematically improve performance in terms of bias and MSE of top estimates with respect to correcting the raw estimates for any of the methods considered (separate estimation was omitted here as it is identical to the raw version). Nonparametric and parametric bootstrapping for Tan2015 yields virtually the same result in this simple case.\\label{fig:varScenSupSparse}}", fig.height = 12, fig.width = 10}
ggplot(data = fullDfSup2b[with(fullDfSup2, mode == "sparse"),], aes(y = value, colour = Method, x = Top_features, linetype = what)) +
        geom_line(linewidth = lineSize) + facet_grid(Performance_measure ~ Cor, scales = "free_y") +
        ylab("Performance measure") + xlab("Number of smallest estimates") +
        scale_linetype_manual(name = "Estimate", values = linValues) + 
        guides(colour = guide_legend(nrow = 4), linetype = guide_legend(nrow = 3)) +
        scale_colour_manual(values = colourVec[names(colourVec) %in% levels(fullDfSup2$Method)]) +
        Theme + geom_hline(data = lineDf, aes(yintercept = value), linetype = "dotted")
```

```{r fulldfdiff, fig.cap = "\\hl{Full simulation results for parametric simulation study: difference in TDR between (rescaled) test statistics and raw estimates for different scenarios (side panels), different methods (colour) and increasing correlation between features (top panels). Positive values indicate higher TDR for the (rescaled) test statistic. The x-axis shows the number of features with smallest estimates considered. The linetype indicates which difference is depicted. The dotted horizontal line indicates zero: no difference. The feature ranking is the same for conditional likelihood as for the raw estimates, and when using test statistics also identical to ash with or without convolution, so in these cases no difference is shown. \\label{fig:varScenSupTDR}}", fig.height = 7, fig.width = 10}
tmp = table(fullDfSup2[fullDfSup2$Performance_measure == "TDR",]$what, fullDfSup2[fullDfSup2$Performance_measure == "TDR",]$Method)
tmp = tmp[rowSums(tmp)>0, colSums(tmp)>0]
fullDfDiff = aggregate(data = fullDfSup2[fullDfSup2$Performance_measure != "TDR",],
                       x = value ~ Method + mode + extreme + Top_features + Cor + Performance_measure, FUN = function(y){
    y[2]-y[1]
})
fullDfDiff$what = "Test statistic - raw"
fullDfDiffTdr1 = aggregate(data = fullDfSup2[fullDfSup2$Performance_measure == "TDR" & fullDfSup2$Method %in% c("BR-squared", "Tan2015 (nonparametric)", "Tan2015 (parametric)", "winnerscurse", "Tweedie's formula\nconvoluted or bagged"),], x = value ~ Method + extreme + mode + Top_features + Cor + mode + Performance_measure, FUN = function(y){
        c(y[2]-y[1], y[3]-y[1])
})
fullDfDiffTdr11 = rbind(cbind(fullDfDiffTdr1[, 1:6], "value" = fullDfDiffTdr1$value[, 1], "what" = "Test statistic - raw"), cbind(fullDfDiffTdr1[, 1:6], "value" = fullDfDiffTdr1$value[, 2], "what" = "Rescaled test statistic - raw"))
fullDfDiffTdr2 = aggregate(data = fullDfSup2[fullDfSup2$Performance_measure == "TDR" & fullDfSup2$Method %in% c("Separate estimation", "Raw"),], x = value ~ Method + extreme + Top_features + Cor + mode + Performance_measure, FUN = function(y){
        y[2]-y[1]
})
fullDfDiffTdr2$what = "Test statistic - raw"
fullDfDiffTdr3 = aggregate(data = fullDfSup2[fullDfSup2$Performance_measure == "TDR" & fullDfSup2$Method %in% c("Ash", "Ash convoluted"),], x = value ~ Method + extreme + mode + Top_features + Cor + Performance_measure, FUN = function(y){
        y[2]-y[1]
})
fullDfDiffTdr3$what = "Rescaled test statistic - raw"
fullDfDiff2 = rbind(fullDfDiff, fullDfDiffTdr11, fullDfDiffTdr2, fullDfDiffTdr3)
fullDfDiff2$what = factor(fullDfDiff2$what, levels = c("Test statistic - raw", "Rescaled test statistic - raw"), ordered = TRUE)
ggplot(data = fullDfDiff2[fullDfDiff2$Performance_measure == "TDR",], aes(y = value, colour = Method, x = Top_features, linetype = what)) +
        geom_line(linewidth = lineSize) + facet_grid(mode ~ Cor, scales = "free_y") +
        ylab("Difference in TDR") + xlab("Number of smallest estimates") +
        scale_linetype_discrete(name = "Difference") + 
        guides(colour = guide_legend(nrow = 3), linetype = guide_legend(nrow = 2)) +
        scale_colour_manual(values = colourVec[names(colourVec) %in% levels(fullDfSup2$Method)]) +
        Theme + geom_hline(aes(yintercept = 0), linetype = "dotted")
```

```{r fulldfdiffMSE, fig.cap = "\\hl{Full simulation results for parametric simulation study: difference in MSE between rescaled test statistics and raw estimates for different scenarios (side panels), different methods (colour) and increasing correlation between features (top panels). Positive values indicate higher MSE for the (rescaled) test statistic. The x-axis shows the number of features with smallest estimates considered. The dotted horizontal line indicates zero: no difference.\\label{fig:varScenSupMSE}}", fig.height = 7, fig.width = 10}
#MSE
ggplot(data = fullDfDiff2[fullDfDiff2$Performance_measure == "MSE",], aes(y = value, colour = Method, x = Top_features)) +
        geom_line(linewidth = lineSize) + facet_grid(mode ~ Cor, scales = "free_y") +
        ylab("Difference in MSE: rescaled test statistic - raw") + xlab("Number of smallest estimates") +
        guides(colour = guide_legend(nrow = 3), linetype = guide_legend(nrow = 2)) +
        scale_colour_manual(values = colourVec[names(colourVec) %in% levels(fullDfSup2$Method)]) +
        Theme + geom_hline(aes(yintercept = 0), linetype = "dotted")
```

\clearpage

### The rank conditional coverage \label{sec:rcc}

The rank conditional coverage (RCC) is shown in \fref{fig:rcc} for the parametric simulation study (dense scenario), reflecting the average coverage at each rank separately. Yet there is no material difference between the coverage results of the RCC and overall coverage, except that the RCC is more variable.

```{r ciRCC, fig.cap = "Rank conditional coverage (RCC) of 95\\% confidence intervals (y-axis) for estimates of different methods (colour) for different ranks (x-axis) for increasing correlation (top panels) and different simulation scenarions (side panels) in parametric simulation. \\label{fig:rcc}", fig.height = 6, fig.width = 9}
rccDf = plotCoverage(baseTestVarCI, extremes = extVec, Methods2Plot = c(plotNormal, "estOutSplit"), quantity = "RCC", convertMethod = TRUE, returnDf = TRUE)
ggplot(data = rccDf[with(rccDf, extreme == "small" & what == "Raw estimate"), ],
       aes_string(y = "value", colour = "Method", x = "Top_features")) +
        geom_line(size = lineSize) + facet_grid(mode ~ Cor, scales = "free_y") +
    xlab("Number of extreme estimates") +
    geom_hline(yintercept = 1-sigLevel, linetype = "dotted") +
    ylab("Rank conditional coverage") +
    scale_colour_manual(values = colourVec[names(colourVec) %in% unique(rccDf$Method)])
rm(baseTestVarCI) 
```

\clearpage

### Selection by controlling the false discovery rate \label{sec:tdr}

\hl{So far, only the top $h$ estimates were considered, with $h$ running from 1 to 30. Here we look at the selection of features while controlling the false discovery rate (FDR) using the procedure by \textcite{Benjamini1995}. All features with an adjusted p-value below `r sigLevel` and point estimate below 0 (since we are interested in small estimates) were retained and the relevant performance measures calculated for them. The results are shown in \fref{fig:tdr}, and reveal that for the sparse scenario, the winner's curse effect is weak, the TDR is high and the empirical coverage of the confidence intervals comes close to their nominal level for most methods. The reason is that in most simulations, all 0.1$p$ features with mean drawn from a distribution with non-zero mean are selected, and none of the other $0.9p$. In the dense scenario, all methods perform much worse, as it becomes harder to correctly rank the features compared to the sparse scenario. A drawback of the FDR approach is that it is only applicable when a null value for $\gamma$ is available.}

```{r fdrSel, fig.cap = paste("\\hl{Simulation results for mean estimation in parametric simulation for different performance measures (side panels) for different selection bias correction methods (colour) and increasing correlation between features (top panels) in different scenarios (point shapes). The performance measures were calculated on all features with adjusted p-values below", sigLevel, "and with point estimate smaller than 0. The dotted horizontal line indicates zero bias or MSE or the nominal coverage of 95\\%, respectively. \\label{fig:tdr}}"), fig.height = 8.5, fig.width = 10}
if(!file.exists(fdrFile <- "Results/fdrFile.RData")){
    p1fdr = plotSimpleResFDR(baseTestVarCI, p = p, Methods2Plot = plotNormalPaper, returnDf = TRUE)
    p2fdr = plotSimpleResFDR(baseTestVarCI, p = p, power = 2, Methods2Plot = plotNormalPaper, returnDf = TRUE)
    p3fdr = plotRunningTDPfdr(baseTestVarCI, extremes = extVec, p = p, Methods2Plot = plotNormalPaper, returnDf = TRUE) 
    p4fdr = plotCoverageFDR(baseTestVarCI, Methods2Plot = plotNormalPaper, returnDf = TRUE) 
    p5fdr = plotWidthCIfdr(baseTestVarCI, Methods2Plot = plotNormalPaper, returnDf = TRUE) 
    fdrDf = rbind(p1fdr, p2fdr, p3fdr, p4fdr, p5fdr)
    fdrDf$Method = factor(fdrDf$Method, ordered = TRUE, labels = methodLabels, levels = methodLevels)
    fdrDf$Performance_measure = factor(fdrDf$Performance_measure, ordered = TRUE, levels = c("Bias", "MSE", "TDR", "Coverage", "Width"))
    save(fdrDf, file = fdrFile)   
} else load(fdrFile)
ggplot(data = fdrDf[fdrDf$extreme == "small", ], aes(y = value, colour = Method, x = Method, shape = mode)) +
    geom_point() + facet_grid(Performance_measure ~ Cor, scales = "free_y") +
    xlab("Method") + ylab("Performance measure") + scale_shape_discrete(name = "Scenario") +
    scale_colour_manual(values = colourVec[names(colourVec) %in% levels(fdrDf$Method)])+ 
    geom_hline(data = lineDf, aes(yintercept = value), linetype = "dotted") +
    theme(axis.text.x = element_blank())
```

\clearpage

## Empirical comparison of correction methods: nonparametric simulations

The results of the non-parametric simulation are shown in \fref{fig:nonPar} in the main text and for the overall MSE in \fref{fig:nonparamMSE}. The separate estimation and \textit{winnerscurse} methods have the highest overall MSE, ash and VanZwet2021 the lowest.

```{r loadBRas1}
load("Data/geneExpr1.RData")
load("Data/sampleData1.RData")
phenId1 = c("leaf.8.width.(76.DAS)", "leaf.8.length.(76.DAS)", "leaf.count.(74.DAS)",
            "total.seed.count", "plant.height.(278.DAS)", "total.shoot.dry.weight.(wo.seeds)")
namPhen = names(phenId1) = c("Leaf 8 width", "Leaf 8 length", "Leaf count",
                             "Total seed count", "Plant height", "Total shoot dry weight")
#Choose some phenotypes with high single feature prediction
rlogBras1 = rlogBras1[rownames(phenData1),]
Loc1 = phenData1[, c("x", "y")]
Nugget = 0.25; Range = max(dist(Loc1))/2;corVals = c(Range, Nugget)
csGaus = corGaus(form = ~ x + y, nugget = TRUE, value = corVals)
```

```{r bras1trialSplit}
numSplits = 20
topFeatsSplit = 1e3 
if(!file.exists(UniMSEcvFileSplit <- "Results/UniMSECVsplit.RData")){ 
    UniMSECVsplit = lapply(phenId1, function(phen){
        dir.create(saveFolder <- paste0("Results/UniCVsplit/", phen))
            if(!file.exists(uniFile <- paste0("Results/UniCVsplit/", phen, ".RData"))){
            cat("Phenotype", phen, "\n")
            id <-!is.na(phenData1[,phen])
            dat = list("x" = rlogBras1[id,], "y" = phenData1[id,phen])
            Loc1b = Loc1[id,]
            #Use top 5000 features
            relExpr = colSums(dat$x)
            dat$x = dat$x[,relExpr >= sort(relExpr, decreasing = TRUE)[topFeatsSplit]]
            n = length(dat$y)
            splitsList = lapply(seq_len(numSplits), function(i){
                id = sample(n, n/2)
                datIn = datOut = dat
                datIn$y = datIn$y[id];datIn$x = datIn$x[id,]
                datOut$y = datOut$y[-id];datOut$x = datOut$x[-id,]
                nesCV = nestedCV(dat = datIn, cvSplits = cvSplits, nCores = nCores, loc = Loc1[id,], verbose = TRUE,
                 control = lmeCon, nOuterFolds = nOuterFolds, correlation = csGaus, saveFolder = saveFolder)
                boCV = bootCV(datIn, n/2, bootReps, cvSplits = cvSplitsBoot, loc = Loc1[id,],correlation = csGaus,
                              control = lmeCon, mseOut = TRUE, verbose = TRUE, nCores = nCores)
                CVout = nestedCV(dat = datOut, cvSplits = cvSplits, nCores = nCores, loc = Loc1[-id,], verbose = TRUE,
                 control = lmeCon, nOuterFolds = nOuterFolds, correlation = csGaus, saveFolder = saveFolder)
                #Splitting
                idInSplit = sample(n/2, n/4)
                datInSplit = datOutSplit = datIn
                LocSplit = Loc1[id,]
                datInSplit$y = datIn$y[idInSplit];datInSplit$x = datIn$x[idInSplit,]
                datOutSplit$y = datIn$y[-idInSplit];datOutSplit$x = datIn$x[-idInSplit,]
                CVinSplit = simpleCV(datInSplit, n/4, correlation = csGaus, loc = LocSplit[idInSplit,], control = lmeCon)
                CVoutSplit = nestedCV(datOutSplit, cvSplits = cvSplits, nrow(datOutSplit$x), correlation = csGaus,
                                          loc = LocSplit[-idInSplit,], control = lmeCon,
                                          nOuterFolds = nOuterFolds, nCores = nCores, saveFolder = saveFolder, verbose = FALSE)
                outList = list("nesCV" = nesCV, "boCV" = boCV, "CVout" = CVout, "CVinSplit" = CVinSplit, 
                               "CVoutSplit" = CVoutSplit, "id" = id)
                })
            save(splitsList, file = uniFile)
            } else {load(uniFile)}
        splitsList
    })
    QuantCut = 0.5
    estsListSplit = lapply(names(phenId1), function(phen){
        cat(phen, "\t")
        phenRes = mclapply(mc.cores = 2, UniMSECVsplit[[phen]], function(y){
                rawEsts = sqrt(sapply(y$nesCV, function(x) x["Bates", "MSEhat"]))
                outEsts = sqrt(sapply(y$CVout, function(x) x["Bates", "MSEhat"]))
                outSEests = sapply(y$CVout, function(x) x["Bates", "SE"])/(2*outEsts)
                seEsts = sapply(y$nesCV, function(x) x["Bates", "SE"])/(2*rawEsts)
                splitEstsIn = sqrt(sapply(y$CVinSplit, function(xx) mean(unlist(xx))))
                splitEstsOut = sqrt(sapply(y$CVoutSplit, function(xx) xx["Bates", "MSEhat"]))#Naive
                bootEstsIn = sqrt(t(sapply(y$boCV, function(x) x[,"MSE"])))
                bootEstsOut = sqrt(t(sapply(y$boCV, function(x) x[,"MSEout"])))
                bootCorrectTan = TanCorrect(rawEsts, bootEstsIn)$est
                bootCorrectForde = forde(rawEsts, Vars <- seEsts^2, allowInflate = TRUE)
                bootCorrectFaye = FayeCorrect(rawEsts, bootEstsIn, bootEstsOut)$est
                alpha1 = getAlpha1(bootEstsIn)
                tweedieCorrectConvList = tweedieForm(rawEsts, varsEstimates = Vars, alpha1 = alpha1)
                tweedieCorrectConv = tweedieCorrectConvList["tweedieCorEsts", ]
                tweedieCorrectConv[tweedieCorrectConv<0] = 0
                tweedieCorrectConvTrunc = tweedieCorrectConv
                cutPoint = quantile(rawEsts, QuantCut)
                idCut <- (rawEsts > cutPoint)
                tweedieCorrectConvTrunc[idCut] = rawEsts[idCut]
                varBoot = apply(bootEstsIn, 2, var)
                tweedieCorrectConvBootList = tweedieForm(rawEsts, varsEstimates = varBoot, alpha1 = alpha1)
                tweedieCorrectConvBoot = tweedieCorrectConvBootList["tweedieCorEsts", ]
                tweedieCorrectConvBoot[tweedieCorrectConvBoot<0] = 0
                tweedieCorrectConvBootTrunc = tweedieCorrectConvBoot
                tweedieCorrectConvBootTrunc[idCut] = rawEsts[idCut]
                ashObj = ash(rawEsts, seEsts, method = "shrink", mode = "estimate")
                ashObjConv = getConvAsh(rawEsts, seEsts, alpha1, ashObj = ashObj)
                # cat scores
                mr = mean(rawEsts)
                rawsCatBoot = crossprod.powcor.shrink(bootEstsIn, rawEsts-mr, alpha = -1/2, verbose = FALSE)+mr
               #Good estimation of correlation matrix
                id = rawsCatBoot<0; rawsCatBoot[id] = rawEsts[id]
                zwetRes = zwetWrapper(z = (rawEsts-mr)/seEsts, s = seEsts)
                zwetResConv = if(alpha1 > 0){
                    zEsts = rnorm(length(rawEsts)*10, rawEsts, seEsts*sqrt(alpha1))
                    zwetWrapper(z = (rawEsts-mr)/seEsts, s = seEsts, zEsts = (zEsts-mr)/seEsts)
                } else {
                    zwetRes
                }
                resMat = cbind("rawEst" = rawEsts, "split" = splitEstsIn, "estOutSplit" = splitEstsOut, 
                      "TanCorrectEst" = bootCorrectTan, "FayeCorrectEst" = bootCorrectFaye, "outEst" = outEsts, 
                      "bootCorrectForde" = bootCorrectForde, "VanZwet2021" = zwetRes$res$betaHats + mr,
                      "VanZwet2021conv" = zwetResConv$res$betaHats + mr, "outSEests" = outSEests,
                      "tweedieConvEsts" = tweedieCorrectConv, "tweedieConvEstsTrunc" = tweedieCorrectConvTrunc, 
                      "tweedieConvEstsBoot" = tweedieCorrectConvBoot, "tweedieConvEstsTruncBoot" = tweedieCorrectConvBootTrunc, 
                      "ash" = ashObj$result$PosteriorMean, "ashConv" = ashObjConv$result$PosteriorMean, "rawsCatBoot" = c(rawsCatBoot))
                #Normalized test statistics
                rawEstsZ = rawEsts/seEsts
                bootCorrectForde = forde(rawEstsZ, 1, allowInflate = TRUE)
                tweedieCorrectConvZ = tweedieForm(rawEsts, varsEstimates = 1, alpha1 = alpha1)["tweedieCorEsts", ]
                tweedieCorrectConvZ[tweedieCorrectConvZ<0] = 0
                tweedieCorrectConvTruncZ = tweedieCorrectConvZ
                cutPoint = quantile(rawEstsZ, QuantCut)
                tweedieCorrectConvTruncZ[rawEstsZ > cutPoint] = rawEstsZ[rawEstsZ > cutPoint]
                ashObjZ = ash(rawEstsZ, 1, method = "shrink", mode = "estimate")
                ashObjConvZ = getConvAsh(rawEsts, seEsts, alpha1, ashObj = ashObjZ)
                resMatZ = seEsts*cbind("bootCorrectForde_rescaled" = bootCorrectForde,
                      "tweedieConvEsts_rescaled" = tweedieCorrectConvZ, "tweedieConvEstsTrunc_rescaled" = tweedieCorrectConvTruncZ, 
                      "ash_rescaled" = ashObjZ$result$PosteriorMean, "ashConv_rescaled" = ashObjConvZ$result$PosteriorMean[seq_along(ashObjZ$result$PosteriorMean)])
                resMat = cbind(resMat, resMatZ)
                rawCI = rawEsts + outer(seEsts, Quants)
                sepCI = outEsts + outer(outSEests, Quants)
                tweedieConvEstsTruncCi = buildCredInt(tweedieCorrectConvList, n = n, Vars, Quants)
                tweedieConvEstsTruncBootCi = buildCredInt(tweedieCorrectConvBootList, n = n, varBoot, Quants)
                tweedieConvEstsTruncCi[idCut,] = tweedieConvEstsTruncBootCi[idCut, ] = rawCI[idCut,]
                ciList = list("rawEst" = rawCI, "estOutSplit" = sepCI,
                 "tweedieConvEstsTrunc" = tweedieConvEstsTruncCi,
                 "tweedieConvEstsTruncBoot" = tweedieConvEstsTruncBootCi,
                 "ash" = ashci(ashObj, level = sigLevel, trace = FALSE),
                 "ashConv" = ashci(ashObjConv, level = sigLevel, trace = FALSE),
                 "rawsCatBoot" = c(rawsCatBoot) + outer(seEsts, Quants),
                 "VanZwet2021" = zwetRes$confInts + mr, "VanZwet2021conv" = zwetResConv$confInts + mr
                )
                list("resMat" = resMat, "ciList" = ciList)
            })
        MSEres = rowMeans(sapply(phenRes, function(x){
            colMeans(na.rm = TRUE, (x$resMat[, "outEst"]-x$resMat[, !(colnames(x$resMat) %in% c("outEst", "split", "outSEests"))])^2)
        }))
        BiasRes = sapply(extVec, function(ext){
            rowMeans(na.rm = TRUE, sapply(phenRes, function(x){
                        c(apply(x$resMat[, !(colnames(x$resMat) %in% c("outEst", "split", "estOutSplit", "outSEests"))], 2, errorLarge, p = nrow(x$resMat), 
                              trueVec = x$resMat[, "outEst"], ext = ext, power = 1)["small",],
                          "estOutSplit" = unname(errorLarge(x$resMat[, "estOutSplit"], ests2rank = x$resMat[, "split"], p = nrow(x$resMat), 
                              trueVec = x$resMat[, "outEst"], ext = ext, power = 1)["small"]))
            }))
        })
        MSEtopRes = sapply(extVec, function(ext){
            rowMeans(na.rm  = TRUE, sapply(phenRes, function(x){
                        c(apply(x$resMat[, !(colnames(x$resMat) %in% c("outEst", "split", "estOutSplit", "outSEests"))], 2, errorLarge, p = nrow(x$resMat), trueVec = x$resMat[, "outEst"], outSEs = x$resMat[, "outSEests"], ext = ext, power = 2)["small",],
                          "estOutSplit" = unname(errorLarge(x$resMat[, "estOutSplit"], ests2rank = x$resMat[, "split"], p = nrow(x$resMat),
                              trueVec = x$resMat[, "outEst"], ext = ext, power = 2)["small"]))
            }))
        })
         TDP = sapply(extVec, function(tdpTop){
             rowMeans(na.rm = TRUE, sapply(phenRes, function(x){
                    rownames(x$resMat) = seq_len(nrow(x$resMat))
                    apply(x$resMat[, !(colnames(x$resMat) %in% c("outEst", "estOutSplit", "outSEests"))], 2,function(y){
                       rankObs = rank(y)
                       trueRanks = rank(x$resMat[, "outEst"])
                        "small" = mean(names(y)[rankObs <= tdpTop] %in% names(trueRanks)[trueRanks <= tdpTop]) 
                    })
            }))
         })
        WidthRes = sapply(extVec, function(ext){
            rowMeans(sapply(phenRes, function(x){
                 tmp = vapply(names(x$ciList), FUN.VALUE = double(ext), function(y){
                        id = rank(x$resMat[, y]) <= ext    
                        x$ciList[[y]][id, 2] - x$ciList[[y]][id, 1]
                 })
                 if(is.matrix(tmp)) colMeans(tmp) else tmp
            }))
        })
         list("MSEtotal" = MSEres, "Bias" = BiasRes, "MSE" = MSEtopRes, "TDP" = TDP, "phenRes" = phenRes, "WidthRes" = WidthRes)
    });names(estsListSplit) = names(phenId1)
    save(UniMSECVsplit, estsListSplit, file = UniMSEcvFileSplit)
} else load(UniMSEcvFileSplit) 
```

```{r nonparPlot, include = FALSE}
moltNonpar = melt(lapply(estsListSplit, function(x) x[c("Bias", "MSE", "TDP", "WidthRes")]), varnames = c("Method", "Top_features"))
names(moltNonpar)[4:5] = c("Performance_measure", "Phenotype") 
moltNonpar$Performance_measure[moltNonpar$Performance_measure == "TDP"] = "TDR"
moltNonpar$Performance_measure[moltNonpar$Performance_measure == "WidthRes"] = "Width" 
moltNonpar$what = "Raw estimate"
moltNonpar$what[grep(moltNonpar$Method, pattern = "rescale")] = "Rescaled test statistic"
moltNonpar$what = factor(moltNonpar$what, levels = names(linValues), labels = names(linValues), ordered = TRUE)
moltNonpar$Method = factor(gsub(pattern = "_rescaled", "", moltNonpar$Method), levels = methodLevels, labels = methodLabels, ordered = TRUE)
moltNonpar$Phenotype = factor(moltNonpar$Phenotype, levels = names(UniMSECVsplit), labels = names(UniMSECVsplit), ordered = TRUE)
#Rename, cause not really cat scores
moltNonpar$Method[moltNonpar$Method== "Cat scores\nnonparametric bootstrap"] = "Decorrelated\nestimates"
moltNonpar = moltNonpar[with(moltNonpar, !(Method %in% c("Tweedie's formula\nconvoluted or bagged", "Tweedie's formula\nconvoluted or bagged\n(truncated, bootstrap SE)", "Tweedie's formula\nconvoluted or bagged(bootstrap SE)"))),]
moltNonpar = droplevels(moltNonpar[!with(moltNonpar, grepl(x = as.character(Method), pattern = "Decorrelated\nestimates") & Performance_measure != "TDR"),])
#Center bias and MSE
moltNonpar = moltNonpar[moltNonpar$what == "Raw estimate",]
idMSE = moltNonpar$Performance_measure=="MSE";idBias = moltNonpar$Performance_measure=="Bias";idWidth = moltNonpar$Performance_measure=="Width"
moltNonpar[idMSE, "value"] = 
        unsplit(lapply(split(moltNonpar[idMSE, "value"], moltNonpar$Phenotype[idMSE]), function(x){
            (x-min(x, na.rm = TRUE))/diff(range(x, na.rm = TRUE))
        }), moltNonpar$Phenotype[idMSE])
moltNonpar[idBias, "value"] = 
        unsplit(lapply(split(moltNonpar[idBias, "value"], moltNonpar$Phenotype[idBias]), function(x){
            x/max(abs(x), na.rm = TRUE)
        }), moltNonpar$Phenotype[idBias])
moltNonpar[idWidth, "value"] = 
        unsplit(lapply(split(moltNonpar[idWidth, "value"], moltNonpar$Phenotype[idWidth]), function(x){
            (x-min(x, na.rm = TRUE))/diff(range(x, na.rm = TRUE))
        }), moltNonpar$Phenotype[idWidth])
ggplot(data = moltNonpar, aes(x = Top_features, y = value, colour = Method)) + geom_line() + 
    scale_colour_manual(values = colourVec[names(colourVec) %in% unique(moltNonpar$Method)])+
    facet_grid(Performance_measure ~ Phenotype, scales = "free_y") + geom_hline(yintercept = 0, linetype = "dotted") +
    geom_hline(linetype = "dotted", aes(yintercept = yintercept), data = data.frame(Performance_measure = c("MSE", "Width"), yintercept = 1)) +
    ylab("Performance measure") + theme(legend.position = "top") + xlab("Number of smallest estimates") +
    guides(colour = guide_legend(nrow = 3))
ggsave("ManuscriptSelBias/Graphs/nonPar.pdf", height = 10, width = 9)
```

```{r figMSE, fig.cap = "Log10 of the ratio of the average MSE over repeated splits of all estimates of each method to the average MSE of all the raw estimates (y-axis) for different methods (colour) and outcome phenotypes (x-axis) in the nonparametric simulation study. Values below 0 indicate more accurate estimation of the collection of all estimates compared to the raw estimates. \\label{fig:nonparamMSE}", fig.width = 8, fig.height = 5.5}
mat = sapply(estsListSplit, function(x) x$MSEtotal) 
mseMolt = melt(mat, value.name = "MSE", varnames = c("Method", "Phenotype"))
mseMolt = mseMolt[grep(mseMolt$Method, pattern = "rescale", invert = TRUE),]
mseMolt$Method = factor(mseMolt$Method, levels = c(methodLevels, "tweedieCorrectBootTrunc"), labels = c(sub("Tweedie's formula\nconvoluted$", "Tweedie's formula\nconvoluted", methodLabels), "Tweedie's formula\nconvoluted (bootstrap SE)"))
dp = mseMolt[with(mseMolt, !(Method %in% c("Tweedie's formula\nconvoluted or bagged", "Cat scores\nnonparametric bootstrap", "Tweedie's formula\nconvoluted or bagged(bootstrap SE)", "Tweedie's formula\nconvoluted or bagged\n(truncated, bootstrap SE)"))),]
dp$MSEstand = log10(sapply(seq_len(nrow(dp)), function(i){
    dp$MSE[i]/dp$MSE[dp$Method=="Raw" & dp$Phenotype == dp$Phenotype[i]]
}))
ggplot(data = dp, aes(y = MSEstand, colour = Method, x = Phenotype, group = Method)) + geom_point() + geom_line() + scale_colour_manual(values = colourVec[names(colourVec) %in% unique(dp$Method)]) +
    theme(axis.title.x = element_blank(), legend.position = "top") + guides(colour = guide_legend(nrow = 3)) +
    ylab("log10(Ratio to raw MSE)")
```

### Correlation between RMSE estimates

```{r corRMSE, fig.height = 5, fig.width = 8.75, fig.cap = "Boxplots of correlation between RMSE estimators for the nonparametric simulation study, estimated through the empirical correlation over repeated sample splits. The limited spread of the estimates corroborates the proposed compound symmetric correlation working model. \\label{fig:corRMSEests}"}
corMatsSplit = lapply(names(UniMSECVsplit), function(phen){ 
    corMat = cor(t(sapply(UniMSECVsplit[[phen]], function(y){
            sqrt(sapply(y$nesCV, function(x) x["Bates", "MSEhat"]))
    })))
    corMat[lower.tri(corMat)]
});names(corMatsSplit) = names(UniMSECVsplit)
corMolt = melt(corMatsSplit)
par(mar = c(6,5,4,4))
labs = names(UniMSECVsplit)
labs[6] = "Total shoot\ndry weight"; labs[4] = "Total seed\ncount"
corMolt$L1 = factor(corMolt$L1, ordered = TRUE, levels = names(corMatsSplit), 
                    labels = labs)
boxplot(data = corMolt, value ~ L1, ylab = "Correlation between RMSE estimators", 
        xlab = "", horizontal = FALSE, las = 1, cex.axis = 0.75)
abline(h = c(0,1), lty = "dotted")
rm(UniMSECVsplit) 
```

\clearpage

# Case study on \emph{B. napus} field trial \label{sec:fieldTrial}

## Analysis

The selection bias correction methods investigated were applied to the single-gene RMSE estimates of the 5,000 most expressed genes in the \textit{B. napus} dataset of \textcite{DeMeyer2022}. The prediction models were fitted using generalised least squares (GLS) with Gaussian autocovariance structure \parencite{Pinheiro2021}. In addition, a multigene linear model was fitted using a custom version of elastic net (EN) with mixing parameter $\alpha = 0.5$ which accounts for spatial autocorrelation across the field in the same way as GLS, as implemented in the $pengls$ package \parencite{Hawinkel2022}. In some cross-validation instances, the fit can diverge leading to extremely large RMSE estimates; these instances were removed before calculating the RMSE and its standard error. As in the nonparametric simulation study, the truncated Tweedie estimates are used.

## Results \label{sec:caseMethod}

The selection bias correction methods investigated were applied to the single-gene RMSE estimates of the \textit{Brassica napus} dataset. \fref{fig:brasPredReal} in the main text shows the smallest raw and corrected estimates together with the EN estimate, Table \ref{tab:taxNames} reveals that the top-predictive selected feature is often not the same one after correction, and varies across correction methods. \fref{fig:corresp} depicts the correspondence in feature ranking between methods in terms of spearman correlation between feature estimates (top right) and proportion overlap between top 30 features (bottom left). The cat scores and especially the separate estimation method coincide least with the feature ranking of the other methods (note that the latter method represents a single random split and results may vary when the split is repeated). The overall feature ranking of the raw estimates, Tan2015, VanZwet2021, BR-squared, Tweedie's formula and ash exhibits strong correspondence, except for total seed count where the bootstrap methods' feature ranking anticorrelate with that of some of the the other methods. \fref{fig:scatterReal} plots the corrected versus the raw estimates, revealing a worrying overcorrection of large raw estimates for Tweedie's formula with convolution. For this reason we switched to the truncated Tweedie estimates. The other corrections, on the other hand, appear reliable and no modification was applied, although the ash and VanZwet2021 methods exhibit very strong shrinkage for some phenotypes, leading to all features having virtually the same corrected RMSE estimate.

```{r bras1trial}
# Don't use blocked CV for comparability
topFeats = 5e3
if(!file.exists(UniMSEcvFile <- "Results/UniMSECV.RData")){
    UniMSECV = lapply(phenId1, function(phen){
            id <-!is.na(phenData1[,phen])
            dat = list("x" = rlogBras1[id,], "y" = phenData1[id,phen])
            #Use top 5000 features
            relExpr = colSums(dat$x)
            dat$x = dat$x[,relExpr >= sort(relExpr, decreasing = TRUE)[topFeats]][, seq_len(topFeats)]
            nesCV = nestedCV(dat = dat, cvSplits = cvSplits, nCores = nCores, loc = Loc1[id,], verbose = TRUE,
                             control = lmeCon, nOuterFolds = nOuterFolds, correlation = csGaus, saveFolder = saveFolder)
            boCV = bootCV(dat, nOuterFolds, bootReps, cvSplits = cvSplits, loc = Loc1[id,], control = lmeCon, 
                          mseOut = TRUE, verbose = TRUE, correlation = csGaus, nCores = nCores)
                            id = sample(n, n/2)
            #Separate estimation
            datIn = datOut = dat
            id2 = sample(nrow(dat$x), nrow(dat$x)/2)
            datIn$y = datIn$y[id2];datIn$x = datIn$x[id2,]
            CVin = simpleCV(datIn, nrow(datIn$x), loc = Loc1[id, ][id2, ], nCores = nCores, correlation = csGaus, control = lmeCon)
            #Find SE for smallest one
           rmseIn = sqrt(sapply(CVin, function(xx) mean(unlist(xx))))
            #Find SE for smallest one
            datOut$y = datOut$y[-id2];datOut$x = datOut$x[-id2, idMin <- which.min(rmseIn), drop = FALSE]
            CVout = nestedCV(datOut, nrow(datOut$x), loc = Loc1b[id, ][-id2, ], cvSplits = cvSplits,
                             correlation = csGaus, control = lmeCon, nCores = nCores)
            #Find the overall regression parameters
            dat$y = dat$y - mean(dat$y)
            betas = apply(dat$x, 2, function(x){
                lm.fit(x = cbind(x), y = dat$y)$coef
            })
            list("nesCV" = nesCV, "boCV" = boCV, "CVin" = CVin, "CVout" = CVout, "rmseIn" = rmseIn, "betas" = betas)
    })
    save(UniMSECV, file = UniMSEcvFile)
}
if(!file.exists(estsListCIfile <- "Results/estsListsCI.RData")){
    load(UniMSEcvFile)  
    estsListCI = mclapply(names(phenId1), mc.cores = 1, function(phen){
        cat(phen)
        rawEsts = sqrt(sapply(UniMSECV[[phen]]$nesCV, function(x) x["Bates", "MSEhat"]))
        seEsts = sapply(UniMSECV[[phen]]$nesCV, function(x) x["Bates", "SE"])/(2*rawEsts)
        rawCi = rawEsts + outer(seEsts, Quants)
        bootEstsIn = sqrt(t(sapply(UniMSECV[[phen]]$boCV, function(x) x[,"MSE"])))
        bootEstsOut = sqrt(t(sapply(UniMSECV[[phen]]$boCV, function(x) x[,"MSEout"])))
        bootCorrectTan = TanCorrect(rawEsts, bootEstsIn, ciReturn = FALSE)$est
        #Include Faye correction here
        bootCorrectFaye = FayeCorrect(rawEsts, bootEstsIn, bootEstsOut)$est
        #Separate estimation
        split = UniMSECV[[phen]]$rmseIn
        estOutSplit = seOut = rep(NA, length(rawEsts))
        estOutSplit[idMin <- which.min(split)] = sqrt(UniMSECV[[phen]]$CVout[[1]]["Bates", "MSEhat"])
        seOut[idMin] = UniMSECV[[phen]]$CVout[[1]]["Bates", "SE"]/(2*estOutSplit[idMin])
        estOutSplitCi = estOutSplit + outer(seOut, Quants)
        alpha1 = getAlpha1(bootEstsIn)
        tweedieCorrectBoot = tweedieForm(rawEsts, varsEstimates = seEsts^2, alpha1 = alpha1)
        tweedieCorrectBoot["tweedieCorEsts",][tweedieCorrectBoot["tweedieCorEsts",]<0] = 0
        tweedieCredInt = buildCredInt(tweedieCorrectBoot, vars = seEsts^2, quants = Quants)
        ashObj = ash(rawEsts, seEsts, method = "shrink", mode = "estimate")
        bootCorrectForde = forde(rawEsts, seEsts^2, allowInflate = TRUE)
        ashCI = ashci(ashObj, level = 1-sigLevel, betaindex = seq_along(rawEsts), trace = FALSE)
         ashConv = getConvAsh(rawEsts, seEsts, alpha1 = alpha1, ashObj = ashObj)
        ashConvCI = ashci(ashConv, level = 1-sigLevel, trace = FALSE)
        #Van Zwet
        mr = mean(rawEsts)
        zwetRes = zwetWrapper(z = (rawEsts-mr)/seEsts, s = seEsts)
        zwetResConv = if(alpha1 >0){
            zEsts = rnorm(p*10, rawEsts, seEsts*sqrt(alpha1))
            zwetWrapper(z = (rawEsts-mr)/seEsts, s = seEsts, zEsts = (zEsts-mr)/seEsts)
        } else {zwetRes}
        # cat scores
        rawsCatBoot = crossprod.powcor.shrink(bootEstsIn, rawEsts-mr, alpha = -1/2, verbose = FALSE)+mr
        rownames(ashCI) = rownames(ashConvCI[seq_along(rawEsts),]) = rownames(estOutSplitCi) =names(rawEsts)
        list("ests" = data.frame("rawEst" = rawEsts, "tweedieConvEsts" = tweedieCorrectBoot["tweedieCorEsts",], 
                   "TanCorrectEst" = bootCorrectTan, "phenotype" = phen, "bootCorrectForde" = bootCorrectForde, "VanZwet2021" = zwetRes$res$betaHats + mr, "VanZwet2021conv" = zwetResConv$res$betaHats + mr, "ashConv" = ashConv$result$PosteriorMean[seq_along(rawEsts)], "estOutSplit" = estOutSplit, "rawsCatBoot" = c(rawsCatBoot), "FayeCorrectEst" = bootCorrectFaye, "ash" = ashObj$result$PosteriorMean),
             "ci" = list("Raw"  = rawCi, "Tweedie's formula\nconvoluted" = tweedieCredInt, "Ash convoluted" = ashConvCI,
             "Ash" = ashCI, "Separate estimation" = estOutSplitCi, "VanZwet2021" = zwetRes$confInts + mr, "VanZwet2021 convoluted" = zwetResConv$confInts + mr))
    })
    names(estsListCI) = names(phenId1) 
    save(estsListCI, file = estsListCIfile)
} else load(estsListCIfile) 
```

```{r bras1trialMulti}
Loc1b = Loc1; colnames(Loc1b) = c("xLoc", "yLoc")
#Multivariate model (gls)
if(!file.exists(multiGlsMSEcvFile <- "Results/multiGlsMSECV.RData")){
    multiGlsMSECV = lapply(phenId1, function(phen){
        dat = list("x" = rlogBras1, "y" = phenData1[,phen])
        dat$y = dat$y[id <- !is.na(dat$y)];dat$x = dat$x[id,]
        #Use top 5000 features
relExpr = colSums(dat$x)
dat$x = dat$x[,relExpr >= sort(relExpr, decreasing = TRUE)[topFeats]]
        penglsInit = pengls(maxIter = 1e2, cbind(as.data.frame(dat), Loc1b), glsSt = csGaus, xNames = colnames(dat)[colnames(dat)!="y"], alpha = alpha, outVar = "y", lambda = 10)#Initiate fit
        csGausFit = corGaus(form = ~ xLoc + yLoc, nugget = TRUE,
                            value = c(exp(penglsInit$gls$Coef[1]), exp(penglsInit$gls$Coef[2])/(1+exp(penglsInit$gls$Coef[2]))))
        multiGlsMod = nestedCVglmnet(dat, loc = Loc1b, nOuterFolds, alpha = alpha, cvSplits = cvSplits,
                                      pengls = TRUE, cvType = "random", loss = "MSE", glsSt = csGausFit)
    })
    save(multiGlsMSECV, file = multiGlsMSEcvFile) 
} else load(multiGlsMSEcvFile)
mseMult <- sapply(multiGlsMSECV, function(x) if(x["Bates", "MSEhat"]>0) x["Bates", "MSEhat"] else x["Naive", "MSEhat"])
estsList = lapply(estsListCI, function(x) x$ests)  
estsListMolt = melt(Reduce(f = rbind, estsList), id.vars = c("phenotype"), variable.name = "Method", value.name = "estimate")
highDimEst = data.frame("estimate" = sqrt(mseMult), phenotype = names(mseMult)) 
estsListMolt$Method = factor(estsListMolt$Method, levels = methodLevels, labels = methodLabels)
estsListMolt$phenotype = factor(estsListMolt$phenotype, levels = namPhen, labels = namPhen, ordered = TRUE)
```

```{r predFigPaper, fig.cap = "\\label{fig:histBras}Histograms of RMSE estimates of single-gene models for predicting different phenotypes (plot titles) of the \\emph{Brassica napus} field trial. The RMSE estimate of the multi-gene model is shown as a dashed blue vertical line, the smallest raw estimate of the single-gene models (before winner's curse correction) is shown as dotted vertical line.", fig.height = 6}
par(mfcol = c(3,2), mar = c(4,4,3,0.5))
xlabs = paste(names(phenId1), c("(cm)", "(cm)", "(leaves)", "(seeds)", "(cm)", "(g)"));names(xlabs) = names(phenId1)
namPhen = names(phenId1)
for(i in namPhen){
    dat = estsList[[i]]$rawEst
    hist(dat, breaks = 50, main = gsub("_", " ", i), xlab = xlabs[[i]], 
         xlim = range(c(sqrt(mseMult[[i]]), dat)), freq = FALSE)
    abline(v = sqrt(mseMult[[i]]), col = "blue", lty = "dashed")
    abline(v = min(estsList[[i]]$rawEst), col = "black", lty = "dotted")
}
par(mfrow = c(1,1)) 
```

```{r plotBetasVsMSE, include = FALSE}
load(UniMSEcvFile)
pdf("ManuscriptSelBias/Graphs/BetasVsRMSE.pdf", height = 5.5)
par(mfrow = c(2,3))
foo = lapply(names(UniMSECV), function(x){
    y = sqrt(sapply(UniMSECV[[x]]$nesCV, function(x) x["Bates", "MSEhat"]))
    features = intersect(names(y), names(UniMSECV[[x]]$betas))
    plot(y[features], abs(UniMSECV[[x]]$betas[features]), main = x, xlab = "RMSE", ylab = "Absolute value of the slope")
})
par(mfrow = c(1,1))
dev.off()
```

```{r tableMin}
#Only consider smallest 50% raw estimates for Tweedie
estsList2 = lapply(estsList, function(x){
    quant50 = x[, "rawEst"] > quantile(x[, "rawEst"], 0.5)
    x$tweedieConvEstsTrunc = x[, "tweedieConvEsts"]
    x[quant50, "tweedieConvEstsTrunc"] = x[quant50, "rawEst"]
    x
}) 
genes = sapply(estsListCI, function(x) rownames(x$ci$Raw))
estsListMolt2 = melt(data.frame(Reduce(f = rbind, estsList2), "gene" = unlist(genes)), 
                     id.vars = c("phenotype", "gene"), variable.name = "Method", value.name = "estimate")
estsListMolt2 = estsListMolt2[!(estsListMolt2$Method %in% c("tweedieConvEsts", "tweedieCorrectBootRaw")),]
estsListMolt2$Method = factor(estsListMolt2$Method, levels = methodLevels, labels = methodLabels)
estsListMolt2$phenotype = factor(estsListMolt2$phenotype, levels = namPhen, labels = namPhen, ordered = TRUE)
smallestEst = aggregate(estimate ~ phenotype + Method, data = estsListMolt2, FUN = min, na.rm = TRUE)
smallestEst$estimate[smallestEst$Method == "Separate estimation"] = sapply(estsList2, function(x){
    min(x$estOutSplit, na.rm = TRUE)
}) 
smallestEstWhich = aggregate(estimate ~ phenotype + Method, data = estsListMolt2, FUN = which.min)
smallestEstWhich$estimate[smallestEstWhich$Method == "Separate estimation"] = sapply(estsList2, function(x){
    which.min(x$estOutSplit)
})
ciSmallest = vapply(FUN.VALUE = double(2), seq_len(nrow(smallestEstWhich)), function(i){
    x = smallestEstWhich[i,]
    if(x[["Method"]] ==  "Tweedie's formula\nconvoluted or bagged\n(truncated)"){
        x[["Method"]] = "Tweedie's formula\nconvoluted"
    }
    if(x[["Method"]] %in% c("Tan2015 (nonparametric)", "Cat scores\nnonparametric bootstrap", "BR-squared", "winnerscurse")){return(c(NA, NA))}
    estsListCI[[x[["phenotype"]]]]$ci[[as.character(x[["Method"]])]][x[["estimate"]], ]
})
ciSmallestPasted = apply(round(ciSmallest, 2), 2, paste, collapse = " - ")
ciCast = dcast(Method ~ phenotype, data = cbind("ci" = ciSmallestPasted, smallestEstWhich[, c("phenotype", "Method")]), value.var ="ci")
dfMins00 = dcast(Method ~ phenotype, data = smallestEst, value.var ="estimate")
dfMins0 = dfMins00; dfMins0[, -1] = round(dfMins0[, -1], 2)
dfMins = sapply(seq_len(nrow(dfMins0)), function(i) paste0(dfMins0[i, -1], " (", ciCast[i, -1], ")"))
dfMins = cbind(as.character(dfMins0$Method), t(dfMins))
dimnames(dfMins) = dimnames(ciCast)
dfMins = as.data.frame(dfMins)
dfMins$Method = factor(dfMins$Method, levels = c(methodLabels, "Elastic net"), 
                       labels = c(methodLabels, "Elastic net"), ordered = TRUE)
elNetRMSEs = sqrt(mseMult)
seHat = sapply(multiGlsMSECV, function(x) x["Bates", "SE"])/(2*elNetRMSEs)
ciEl = elNetRMSEs + outer(seHat, Quants)
elNetCI = paste0(as.character(round(elNetRMSEs, 2)), " (", 
                round(ciEl[, "lower"], 2), " - ", round(ciEl[, "upper"], 2), ")")
```

```{r tableFig, include = FALSE}
#Will be much clearer in a figure
stripSize = 7 
tabDf = rbind(
    data.frame("phenotype" = names(elNetRMSEs), "Method" = "Elastic net", "estimate" = elNetRMSEs, ciEl),
    data.frame(smallestEst, t(ciSmallest))
)
tabDf$Method = factor(tabDf$Method, levels = c("Elastic net", methodLabels), labels = c("Elastic net", methodLabels), ordered = TRUE)
Dodge = .04; Width = .025;pointSize = 1.7
namPhenUnits = paste(namPhen, c("(cm)", "(cm)", "(leaves)", "(seeds)", "(cm)", "(g)"))
tabDf$phenotype = factor(tabDf$phenotype, levels = namPhen, labels = namPhenUnits, ordered = TRUE) 
ggplot(data = tabDf[!(tabDf$Method %in% c("Cat scores\nnonparametric bootstrap")),], aes(y = estimate, col = Method, x = 1)) + 
    geom_point(position = position_dodge(Dodge), size = pointSize) +
    facet_grid(phenotype ~ ., scales = "free_y")+ 
    scale_colour_manual(values = c(colourVec, "Elastic net" = "#CC6699")) +
    geom_errorbar(aes(ymin=lower, ymax=upper), width = Width, position=position_dodge(Dodge)) +
    geom_hline(col = "#CC6699", aes(yintercept = estimate), data = tabDf[tabDf$Method== "Elastic net", c("phenotype", "estimate")],
               linetype = "dotted") +
    ylab("RMSE estimate") +
    xlab("") +
    theme(axis.text.x = element_blank(), axis.ticks.x = element_blank(), strip.text.y = element_text(size = stripSize))
ggsave("ManuscriptSelBias/Graphs/BrasPredReal.pdf", height = 8, width = 7.5) 
```

```{r taxTab, results = "asis"}
#Use 5000 most expressed genes
relExpr = colSums(rlogBras1) 
taxNames = names(relExpr)[relExpr >= sort(relExpr, decreasing = TRUE)[topFeats]]
taxList = sapply(namPhen, function(phen){
    sapply(c("rawEst","estOutSplit","TanCorrectEst","FayeCorrectEst","rawsCatBoot","tweedieConvEstsTrunc","ashConv", "VanZwet2021"), function(x){
        taxNames[which.min(estsList2[[phen]][,x])]
    })
})
rownames(taxList) = methodLabels[match(rownames(taxList), methodLevels)]
rownames(taxList)[5] = "Cat scores"; rownames(taxList)[6] = "Tweedie's formula"; rownames(taxList)[7] = "Ash" 
rownames(taxList)[3] = "Tan2015"
taxList = apply(taxList, c(1,2), function(x) gsub("Bna", "", x))
print(row.names = FALSE, size = "tiny", xtable(t(taxList), caption = "\\label{tab:taxNames}Gene with smallest RMSE estimate for different methods. The preposition 'Bna' was omitted from the gene names for brevity. Convolution was used forall empirical Bayes methods (Tweedie's formula, ash and VanZwet2021), and nonparametric bootstrapping for the Tan2015 method."), comment = FALSE)
```

```{r overlap30andSpearman, fig.height = 9, fig.cap = "Pairwise correspondence between feature ranking of different methods (x- and y-axes) for the \\textit{B. napus} case study, summarized by proportion of overlapping features in the top 30 ranked features (bottom left) and by spearman correlation between all RMSE estimates (top right). The entries are coloured from low (orange) to high (blue).\\label{fig:corresp}"}
#Find spearman correlation between features
methodLevelsSpear = c("rawEst", "FayeCorrectEst", "TanCorrectEst", "bootCorrectForde",
                      "tweedieConvEstsTrunc", "ashConv", "rawsCatBoot", "VanZwet2021conv")
names(methodLevelsSpear) = methodLevelsSpear
spearCors = lapply(estsList2, function(x){
    corMat = cor(x[, methodLevelsSpear], method = "spearman")
    smallestEsts = apply(x[, methodLevelsSpear], 2, function(y){
        seq_len(topFeats)[rank(y) <= max(extVec)]
    })
    vennTab = sapply(methodLevelsSpear, function(meth1){
        sapply(methodLevelsSpear, function(meth2){
            length(intersect(smallestEsts[, meth1], smallestEsts[, meth2]))/max(extVec)
        })
    })
    list("Spearman_correlation" = corMat, "Overlap_top30" = vennTab)
})
moltSpearCors = melt(spearCors)
names(moltSpearCors) = c("Method_x", "Method_y", "value", "what", "Phenotype")
#Now keep only upper triangle for Spearman correlation, and lower triangle for overlap
moltSpearCors$Method_x = factor(moltSpearCors$Method_x, levels = methodLevels, labels = methodLabels, ordered = TRUE)
moltSpearCors$Method_y = factor(moltSpearCors$Method_y, levels = methodLevels, labels = gsub("\n", " ", methodLabels), ordered = TRUE)
moltSpearCors = moltSpearCors[with(moltSpearCors, {
    ((as.integer(Method_x) > as.integer(Method_y) & what == "Spearman_correlation") |
    (as.integer(Method_x) < as.integer(Method_y) & what == "Overlap_top30"))
}),]
#Revert levels to get proper ordering
moltSpearCors$Method_y = fct_rev(moltSpearCors$Method_y)
moltSpearCors$value = round(moltSpearCors$value, 2)
moltSpearCors$col[moltSpearCors$what == "Spearman_correlation"] = (moltSpearCors$value[moltSpearCors$what == "Spearman_correlation"]+1)/2
moltSpearCors$col[moltSpearCors$what == "Overlap_top30"] = moltSpearCors$value[moltSpearCors$what == "Overlap_top30"]
moltSpearCors$Phenotype = factor(moltSpearCors$Phenotype, levels = namPhen, labels = namPhen, ordered = TRUE) 
ggplot(moltSpearCors, aes(x = Method_x, y = Method_y, fill = col, label = value)) +
    geom_tile() + geom_text(size = 3) + facet_grid(Phenotype ~ .) +
    theme(panel.grid = element_blank(), axis.text.x = element_text(angle = 90), 
          strip.text.y = element_text(size = stripSize)) +
    xlab("") + ylab("") + 
    scale_fill_gradient(low = "#FF9933", high = "#3300FF", guide = "none")
```

```{r processReal, fig.height = 9.3, fig.cap = "Scatterplots of raw vs. corrected RMSE estimates for the six phenotypes considered (units are shown between brackets in the plot titles). Nonparametric bootstrapping was employed where applicable. Tweedie's formula often overshrinks large raw estimates, whereas ash and VanZwet2021 sometimes shrink all estimates to virtually the same value. The cat scores and \\textit{winnerscurse} estimates have high variability. The black solid line has intercept 0 and slope 1.\\label{fig:scatterReal}"}
par(mfrow = c(4,2), mar = c(4, 4, 3, 3));Pch = 1;Cex = 0.3 
for(phen in namPhen){
    plot(pch = Pch, cex = Cex, estsListCI[[phen]]$ests$TanCorrectEst, x = rawEsts <- estsListCI[[phen]]$ests$rawEst, xlab = "Raw estimate", ylab = "Corrected estimate", main = xlabs[phen], ylim = range(na.rm = TRUE, c(estsListCI[[phen]]$ests[, c("tweedieConvEsts", "TanCorrectEst", "ashConv", "estOutSplit", "FayeCorrectEst", "ash", "bootCorrectForde", "VanZwet2021", "rawsCatBoot")])), col = colourVec["Tan2015 (nonparametric)"])
    points(pch = Pch, cex = Cex, estsListCI[[phen]]$ests$tweedieConvEsts, x = rawEsts, col = colourVec["Tweedie's formula\nconvoluted"])
    points(pch = Pch, cex = Cex, estsListCI[[phen]]$ests$ashConv, x = rawEsts, col = colourVec["Ash convoluted"])
    points(pch = Pch, cex = Cex, estsListCI[[phen]]$ests$ash, x = rawEsts, col = colourVec["Ash"])
    points(pch = Pch, cex = Cex, estsListCI[[phen]]$ests$FayeCorrectEst, x = rawEsts, col = colourVec["BR-squared"])
    points(pch = Pch, cex = Cex, estsListCI[[phen]]$ests$estOutSplit, x = rawEsts, col = colourVec["Separate estimation"])
    points(pch = Pch, cex = Cex, estsListCI[[phen]]$ests$rawsCatBoot, x = rawEsts, col = colourVec["Cat scores\nnonparametric bootstrap"])
    points(pch = Pch, cex = Cex, estsListCI[[phen]]$ests$bootCorrectForde, x = rawEsts, col = colourVec["winnerscurse"])
     points(pch = Pch, cex = Cex, estsListCI[[phen]]$ests$VanZwet2021, x = rawEsts, col = colourVec["VanZwet2021"])
    abline(0,1) #Tan's correction appears more natural
} 
legVec = c("Separate estimation", "BR-squared", "Tan2015 (nonparametric)", "Tweedie's formula\nconvoluted", "Ash", "Ash convoluted", "Cat scores\nnonparametric bootstrap", "winnerscurse", "VanZwet2021")
plot(type = "n", 0, 0, xlab = "", ylab = "", axes = FALSE)
par(mar = c(4, 4, 3, 4.7))
legend("center", ncol = 2, legend = legVec, col = colourVec[legVec], pch = Pch, cex = 0.8) 
```

\clearpage

# Summary statistics of the quadratic loss distribution: the MSE and RMSE \label{supsec:rmse}

In this section we describe the estimation of the RMSE and its standard error (SE) through cross-validation. We perform K-fold cross-validation, assuming for simplicity that $K$ divides $n$. Each fold with index $k = 1,\hdots, K$ contains $n/K$ observations, indicated by the set $\mathcal{I}_k$. With $\mathcal{I}$ the set of all observations, $\mathcal{I}\backslash\mathcal{I}_k$ is referred to as the training folds and $\mathcal{I}_k$ as the left-out fold. A model $m$ with parameters $\bs\theta$ is trained on the training folds, yielding an estimate $\bs\theta_k$. This model can be used to make predictions on the left-out fold, with vector of predictions $\mb{\hat{y}}_k = m(\mb X_k\mid \bs\theta_k)$. Repeating this for all folds $k$ yields predictions $\hat{y}_i$, which are then evaluated by comparing them with the true values $\mb y_i$ using a loss function $r$. We choose the squared loss function $r(y, \hat{y}) = (y-\hat{y})^2$, and will abbreviate $r(y_i, \hat{y}_i) = r_i$. The interest is in the distribution of this loss on test datasets, and especially its expectation, the mean squared error (MSE). Cross-validation (CV) is a way to estimate this loss, so the MSE is estimated as $\widehat{MSE} = \frac{1}{n} \sum_{i=1}^n r_i$, on the assumption that new data would be drawn from the same distribution as the original data. \textcite{Bates2023} proposed a nested cross-validation paradigm to estimate the SE on $\widehat{MSE}$. Yet rather than the mean squared error (MSE), we choose to work with the square root of the MSE or root mean squared error (RMSE) as estimand $\gamma$, because this one more closely approximates the normal sampling distribution, as assumed by several selection bias correction methods employed. The variance of the RMSE as function of the variance of the MSE is approximated by the delta method \parencite{Gauss1823} as:

\begin{equation}
\Var{\widehat{RMSE}} = \text{Var}\left(\sqrt{\widehat{MSE}}\right) \approx \Var{\widehat{MSE}} \left(\frac{d \sqrt{\widehat{MSE}}}{\widehat{dMSE}} \right)^2 = \frac{\Var{\widehat{MSE}}}{4 \widehat{MSE}}
\label{eq:varRMSE}
\end{equation}

# Computation times \label{sec:benchComp}

A computational benchmark of the different methods for the mean estimation  in the parametric simulations is shown in Table \ref{tab:compTimes} for _p_=300 features and a sample size of _n_=50. For the Tan2015 and BR-squared methods, _B_ = 100 bootstrap instances were drawn. For the conditional likelihood method, the 30 smallest and largest corrected estimates were calculated; all other methods find all corrected estimates. When also confidence intervals were calculated (right column), an additional _B_=100 inner bootstraps were run for the Tan2015 and BR-squared methods. Computations were run on a single Intel 2.6 GHz core for 20 repeats, and average computation times over the repeats are reported. Computation times increase roughly quadratically with _B_ for the bootstrap methods because of the nested bootstrap, and increase quickly with _p_ for the parametric bootstrap for the need to estimate the covariance matrix. The bootstrap confidence intervals quickly become computationally unfeasible for larger number of features and more complicated estimation procedures.

```{r compTimes}
benchReps = 20; pComp = 3e2; bootReps = 1e2; quants = c(0.025, 0.975);bootRepsAsh = 10
if(!file.exists(benchCompFile <- "simResults/benchComp.RData")){
    benchComp = vapply(seq_len(benchReps), FUN.VALUE = matrix(0, 14, 2), function(j){
                    meanVec = rnorm(pComp, sd = sdMean); names(meanVec) = seq_len(pComp)
                    mat = mvtnorm::rmvnorm(n = n, mean = meanVec, sigma = Sig <- Sigmas[[as.character(0.5)]][seq_len(pComp), seq_len(pComp)])
                    id = seq_len(n)
                    #Raw
                    pointEsts = c("rawEst" = system.time(colMeans(mat))[3],
                    #Tweedie
                    "tweedieConvEsts" = system.time({
                        rawEsts = colMeans(mat)
                        alpha1 = getAlpha1(mat)
                        tweedieForm(rawEsts, varsEstimates = apply(mat, 2, var)/n, alpha1 = alpha1)
                    })[3],
                    "tweedieConvEstsBoot" = system.time({
                        rawEsts = colMeans(mat)
                        bootEsts = sapply(integer(bootRepsAsh), function(b){
                            id0 = sample(id, replace = TRUE)
                            colMeans(mat[id0,])
                        })
                        alpha1 = getAlpha1(t(bootEsts))
                        tweedieForm(rawEsts, varsEstimates = apply(mat, 2, var)/n, alpha1 = alpha1)
                    })[3],
                    "TanCorrectEst" = system.time({
                        rawEsts = colMeans(mat)
                        bootEsts = sapply(integer(bootReps), function(b){
                            id0 = sample(id, replace = TRUE)
                            colMeans(mat[id0,])
                        })
                        TanCorrect(rawEsts, t(bootEsts))
                    })[3],
                    "TanCorrectEstParam" = system.time({
                        rawEsts = colMeans(mat)
                        paramBootMat = rmvnorm(n = n*bootReps, mean = rawEsts, sigma = cov(mat))
                        bootEsts = sapply(seq_len(bootReps), function(i){
                            colMeans(paramBootMat[seq_len(n)+(i-1)*n,])
                        })
                        TanCorrect(rawEsts, t(bootEsts))
                    })[3],
                    #Faye
                    "FayeCorrectEst" = system.time({
                        rawEsts = colMeans(mat)
                        bootEsts = lapply(integer(bootReps), function(b){
                            id0 = sample(id, replace = TRUE)
                            cbind("estsIn" = colMeans(mat[id0,]), "estsOut" = colMeans(mat[-id0,]))
                        })
                        FayeCorrect(rawEsts, t(sapply(bootEsts, function(x) x[,"estsIn"])), 
                                    t(sapply(bootEsts, function(x) x[,"estsOut"])))
                    })[3],
                    #Split
                    "estOutSplit" = system.time({
                        idSplit = sample(n, size = n/2)
                        estInSplit = colMeans(mat[idSplit,]);estOutSplit = colMeans(mat[-idSplit,])
                    })[3],
                    #Conditional likelihood
                    "condML" = system.time({
                        corrCondML(raw = colMeans(mat), vars = apply(mat,2, var)/nrow(mat), 
                                   dat = mat, extVec, ciReturn = FALSE)
                    })[3],
                     #ash
                    "ash" = system.time({
                        ash(betahat = colMeans(mat), sebetahat = apply(mat, 2, sd)/sqrt(nrow(mat)), method = "shrink", mode = "estimate")
                    })[3],
                    "ashConv" = system.time({
                       ests = colMeans(mat)
                       ses = apply(mat, 2, sd)/sqrt(nrow(mat))
                       alpha1 = getAlpha1(mat)
                    ashObj = getConvAsh(ests, ses, alpha1 = alpha1, ashObj = ash(betahat = ests, ses, method = "shrink", mode = "estimate"))
                    })[3],
                    "catScores" = system.time({shrinkcat.stat(mat, L = rep(0, n), verbose = FALSE)})[3],
                    "bootCorrectForde" = system.time({
                        forde(colMeans(mat), apply(mat, 2, var)/nrow(mat))
                        })[3],
                    "VanZwet2021" = system.time({
                       ests = colMeans(mat)
                       s = apply(mat, 2, sd)/sqrt(nrow(mat))
                       mr = mean(ests)
                       zwetWrapper(z = (ests-mr)/s, s = s, confInt = FALSE)
                    })[3],
                    "VanZwet2021conv" = system.time({
                         ests = colMeans(mat);                       mr = mean(ests)
                       s = apply(mat, 2, sd)/sqrt(nrow(mat))
                       alpha1 = getAlpha1(mat)
                        zwetResConv = if(alpha1 < 0){
                            zwetWrapper(z = (ests-mr)/s, s = s, confInt = FALSE)
                        } else {
                            zEsts = rnorm(p*bootRepsAsh, ests, s*sqrt(alpha1))
                            zwetWrapper(z = (ests-mr)/s, s = s, zEsts = (zEsts-mr)/s, confInt = FALSE)
                        }
                    })[3])
                    ### CONFIDENCE INTERVALS
                     #Raw
                    confInts = c("rawEst" = system.time({
                        colMeans(mat) + outer(apply(mat, 2, sd)/sqrt(n), Quants)
                        })[3],
                    #Tweedie
                    "tweedieConvEsts" = system.time({
                        rawEsts = colMeans(mat)
                         alpha1 = getAlpha1(mat)
                        tweedieForm(rawEsts, varsEstimates = apply(mat, 2, var)/n, alpha1 = alpha1, 
                        ciReturn = TRUE)
                    })[3],
                    "tweedieConvEstsBoot" = system.time({
                        rawEsts = colMeans(mat)
                        bootEsts = sapply(integer(bootRepsAsh), function(b){
                            id0 = sample(id, replace = TRUE)
                            colMeans(mat[id0,])
                        })
                        alpha1 = getAlpha1(t(bootEsts))
                        tweedieForm(rawEsts, varsEstimates = apply(mat, 2, var)/n, alpha1 = alpha1, 
                        ciReturn = TRUE)
                    })[3],
                    #Tan
                    "TanCorrectEst" = system.time({
                        rawEsts = colMeans(mat)
                         bootEsts = lapply(integer(bootReps), function(b){
                                id0 = sample(id, replace = TRUE)
                                out = cbind("estsIn" = colMeans(mat[id0,]), "estsOut" = colMeans(mat[-id0,]))
                                outIn = vapply(seq_len(bootReps), FUN.VALUE = matrix(0, pComp, 2), function(b){
                                        idIn = sample(id, replace = TRUE)
                                        cbind("estsIn" = colMeans(mat[id0,][idIn,]), "estsOut" = colMeans(mat[id0,][-idIn,]))
                                    })
                                list("out" = out, "outIn" = outIn)
                        })
                        TanCorrect(rawEsts, t(sapply(bootEsts, function(x) x$out[, "estsIn"])), 
                                   fullBootObj = bootEsts, ciReturn = TRUE, quants = quants)
                    })[3],
                    "TanCorrectEstParam" = system.time({
                        rawEsts = colMeans(mat)
                        paramBootMat = rmvnorm(n = n*bootReps, mean = rawEsts, sigma = cov(mat))
                        bootEstsParam = lapply(seq_len(bootReps), function(i){
                            out = cbind("estsIn" = colMeans(matIn <- paramBootMat[seq_len(n)+(i-1)*n,]))
                             paramBootMatIn = rmvnorm(n = n*bootReps, mean = out[, 'estsIn'], sigma = cov(matIn))
                             outIn = vapply(seq_len(bootReps), FUN.VALUE = matrix(0, ncol(mat), 2), function(b){
                                    cbind("estsIn" = colMeans(paramBootMatIn[seq_len(n)+(b-1)*n,]),
                                          "estsOut" = NA)
                                })
                            list("out" = out, "outIn" = outIn)})
                        TanCorrect(rawEsts, t(sapply(bootEstsParam, function(x) x$out[, "estsIn"])), 
                                   ciReturn = TRUE, fullBootObj = bootEstsParam, quants = quants)
                    })[3],
                    #Faye
                    "FayeCorrectEst" = system.time({
                        rawEsts = colMeans(mat)
                        bootEsts = lapply(integer(bootReps), function(b){
                                id0 = sample(id, replace = TRUE)
                                out = cbind("estsIn" = colMeans(mat[id0,]), "estsOut" = colMeans(mat[-id0,]))
                                outIn = vapply(seq_len(bootReps), FUN.VALUE = matrix(0, pComp, 2), function(b){
                                        idIn = sample(id, replace = TRUE)
                                        cbind("estsIn" = colMeans(mat[id0,][idIn,]), "estsOut" = colMeans(mat[id0,][-idIn,]))
                                    })
                                list("out" = out, "outIn" = outIn)
                        })
                        FayeCorrect(rawEsts, t(sapply(bootEsts, function(x) x$out[,"estsIn"])), 
                                    t(sapply(bootEsts, function(x) x$out[,"estsOut"])), ciReturn = TRUE, 
                                    fullBootObj = bootEsts, zQuants = Quants)
                    })[3],
                    #Split
                    "estOutSplit" = system.time({
                        idSplit = sample(n, size = n/2)
                        estInSplit = colMeans(mat[idSplit,]);estOutSplit = colMeans(mat[-idSplit,])
                        sdOut = apply(mat[-idSplit,], 2, sd)/sqrt(n/2)
                        CIestOutSplit = estOutSplit + outer(sdOut, Quants)
                    })[3],
                    #Conditional likelihood
                    "condML" = system.time({
                        raws = colMeans(mat)
                        corrCondML(raw = raws, vars = apply(mat,2, var)/nrow(mat), dat = mat, extVec, ciReturn = TRUE)
                    })[3],
                    #ash
                    "ash" = system.time({
                        ashObj = ash(betahat = colMeans(mat), sebetahat = apply(mat, 2, sd)/sqrt(nrow(mat)), 
                                     method = "shrink", mode = "estimate")
                        ashci(ashObj, level = 1-sigLevel, trace = FALSE)
                    })[3],
                    "ashConv" = system.time({
                        ests = colMeans(mat)
                       ses = apply(mat, 2, sd)/sqrt(nrow(mat))
                     alpha1 = getAlpha1(mat)
                    ashObj = getConvAsh(ests, ses, alpha1 = alpha1, ashObj = ash(betahat = ests, ses, method = "shrink", mode = "estimate"))
                        ashci(ashObj, level = 1-sigLevel, trace = FALSE)
                    })[3], "catScores" = NA, "bootCorrectForde" = NA,
                                      "VanZwet2021" = system.time({
                       ests = colMeans(mat)
                       s = apply(mat, 2, sd)/sqrt(nrow(mat))
                       mr = mean(ests)
                       zwetWrapper(z = (ests-mr)/s, s = s, confInt = TRUE)
                    })[3],
                    "VanZwet2021conv" = system.time({
                         ests = colMeans(mat);mr = mean(ests)
                       s = apply(mat, 2, sd)/sqrt(nrow(mat))
                       alpha1 = getAlpha1(mat)
                        zwetResConv = if(alpha1 < 0){
                            zwetWrapper(z = (ests-mr)/s, s = s, confInt = TRUE)
                        } else {
                            zEsts = rnorm(p*bootRepsAsh, ests, s*sqrt(alpha1))
                            zwetWrapper(z = (ests-mr)/s, s = s, zEsts = (zEsts-mr)/s, confInt = TRUE)
                        }
                    })[3])
                    cbind("pointEsts" = pointEsts, "confInts" = confInts)
            })
    save(benchComp, file = benchCompFile)
} else load(benchCompFile) 
```

```{r compTimesTable, results = "asis"}
benchCompMean = apply(benchComp, c(1,2), mean) 
colnames(benchCompMean) = c("Point estimate", "Confidence interval")
rownames(benchCompMean) = gsub("\\(CV SE\\)", "", as.character(factor(gsub(".elapsed", "", rownames(benchCompMean)), levels = methodLevels, labels = methodLabels)))
rownames(benchCompMean) = gsub("bootstrap SE", "bootstrap correlation", rownames(benchCompMean))
print.xtable(comment = FALSE, xtable(benchCompMean[c("Raw", "Separate estimation", "Conditional likelihood", "BR-squared", "Tan2015 (nonparametric)", "Tan2015 (parametric)", "winnerscurse", "Tweedie's formula\nconvoluted or bagged", "Ash", "Ash convoluted", "VanZwet2021", "VanZwet2021 convoluted", "Cat scores"),], display = rep("e", 3), digits = 2, caption = "Average computation times in seconds for calculation of point estimates and confidence intervals for different selection bias correction methods (rows) over 20 repeats.\\label{tab:compTimes}")) 
```

\clearpage

# Recommendations table

\begin{table}
\begin{tabularx}{1.05\textwidth}{|>{\setlength\hsize{0.35\hsize}\setlength\linewidth{\hsize}}X|>{\setlength\hsize{1.3\hsize}\setlength\linewidth{\hsize}}X|>{\setlength\hsize{1.35\hsize}\setlength\linewidth{\hsize}}X|}
\hline
Method & Strengths & Weaknesses \\
\hline
Separate estimation  & \begin{itemize}
\item{Immune to selection bias when estimand is independent of the sample size}
\item{Nominal coverage of the confidence interval}
\item{Mostly robust to dependence}
\end{itemize} &
\begin{itemize}
 \item{Variable estimates}
  \item{Poor feature ranking}
 \item{Biased when estimand depends on sample size (e.g. prediction loss)}
 \item{Random result because of single data split}
\end{itemize}\\
\hline
Conditional likelihood  &
\begin{itemize}
\item{Reasonable correction of selection bias under independence}
\end{itemize} &
\begin{itemize}
\item{Overcorrects selection bias in presence of dependence between estimates}
\item{Variable estimates}
\item{Only applicable to likelihood estimation}
\item{Computationally intensive}
\item{Corrected estimates depend on the number of top features considered}
\end{itemize}\\
\hline
BR-squared &
\begin{itemize}
\item Reasonable performance under dependence
\end{itemize} &
\begin{itemize}
\item Poor performance under independence
\item Computationally intensive
\item Ill-suited to small sample size datasets
\end{itemize}\\
\hline
Tan2015 &
\begin{itemize}
\item Good correction of selection bias regardless of dependence
\item No distributional assumptions
\end{itemize} &
\begin{itemize}
\item Computationally intensive
\item Ill-suited to small sample size datasets
\end{itemize}\\
\hline
\textit{winnerscurse} &
\begin{itemize}
\item Short computation times
\item Relies on estimates and standard errors only, no access to original data needed
\end{itemize} &
\begin{itemize}
\item Changeable correction of winner's curse
\item No measure of uncertainty on corrected estimates
\item High-dimensionality required
\end{itemize}\\
\hline
Tweedie's formula &
\begin{itemize}
\item Accurate and fast estimation with reduced selection bias
\item Good feature ranking performance
\item Robust to dependence thanks to convolution
\end{itemize} &
\begin{itemize}
\item Sensitive to error in estimation of estimator variance and marginal log density, which can lead to overshrinkage
\item High-dimensionality required
\end{itemize}\\
\hline
ash &
\begin{itemize}
\item Accurate point estimates with low selection bias thanks to convolution
\item Good feature ranking performance
\end{itemize} &
\begin{itemize}
\item Assumption of unimodal prior can be violated
\item Underestimates posterior variance, even with convolution
\item High-dimensionality required
\end{itemize}\\
\hline
VanZwet2021 &
\begin{itemize}
\item Robust to dependence thanks to convolution
\item Relies on estimates and standard errors only, no access to original data needed
\end{itemize} &
\begin{itemize}
\item Assumptions on prior distribution can be violated
\item High-dimensionality required
\end{itemize}\\
\hline
Cat scores &
\begin{itemize}
\item Improved feature ranking under strong dependence using test statistics
\end{itemize} &
\begin{itemize}
\item Slight drop in performance in absence of strong dependence
\item Changeable performance when applied to raw estimates
\end{itemize}\\
\hline
\end{tabularx}
\caption{\label{tab:overview}Selection bias correction methods considered in this study with strengths and weaknesses.} 
\end{table}
\clearpage

# Software \label{sec:soft}

The software versions used are detailed below.

```{r sessionInfo}
sessionInfo()
```

\printbibliography
````
